# Earliest Evidence of Elephant Butchery at Olduvai Gorge (Tanzania) Reveals the Evolutionary Impact of Early Human Megafaunal Exploitation

**DOI:** 10.1101/2025.08.28.672811

**Authors:** Manuel Domínguez-Rodrigo, Enrique Baquedano, Abel Moclán, David Uribelarrea, José Ángel Correa Cano, Fernando Diez-Martín, Alejandro Velázquez-Tello, Elia Organista, Eduardo Mendez-Quintas, Marina Vegara-Riquelme, Agness Gidna, Audax Mabulla

## Abstract

The role of megafaunal exploitation in early human evolution remains debated. Occasional use of large carcasses by early hominins has been considered by some as opportunistic, possibly a fallback dietary strategy, and for others a more important survival strategy. At Olduvai Gorge, evidence for megafaunal butchery is scarce in the Oldowan of Bed I, but becomes more frequent and widespread after 1.8 Ma in Bed II, coinciding with the emergence of Acheulean technologies, but not functionally related to the main Acheulian tool types. Here, we present the earliest direct evidence of proboscidean butchery, including a newly documented elephant butchery site (EAK). This shift in behavior is accompanied by larger, more complex occupation sites, signaling a profound ecological and technological transformation. Rather than opportunistic scavenging, these findings suggest a strategic adaptation to megafaunal resources, with implications for early human subsistence and social organization. The ability to systematically exploit large prey represents a unique evolutionary trajectory, with no direct modern analogue, since modern foragers do so only episodically.

## Introduction

Spatially concurrent lithic tools and proboscidean bones have a long history in human evolution. Such concurrency (sometimes in conjunction with other variables) has been frequently interpreted as the result of hominin butchery (Adrien Hannus, 2018; Agam and Barkai, 2018, 2016; Altamura et al., 2020; Barkai, 2019; Ben-Dor and Barkai, 2020; Berthelet and Chavaillon, 2001; Boschian and Saccà, 2015; Chavaillon et al., 1987; Chavaillon and Berthelet, n.d.; Delagnes et al., 2006; Gaudzinski et al., 2005; Gaudzinski-Windheuser et al., 2023a, 2023b; Gomez et al., 1978; Goren-Inbar et al., 1994; Haynes, 2022; Konidaris et al., 2021; Leakey, 1971; Lemorini et al., 2023; Lev and Barkai, 2016; Mussi and Villa, 2008; Reshef and Barkai, 2015; Rocca et al., 2021; Saccà, 2012; Santucci et al., 2016; Solodenko et al., 2015; Villa et al., 2005; Yravedra et al., 2019, 2010; Rabinovich et al., 2012). Spatial concurrency is insufficient to establish a causal relationship between tools and megafaunal remains. Given the time-averaged nature of a large part of the Paleolithic record, the deposition of tools and elephants could have occurred sequentially and unrelatedly in several sites; especially taking into account that most of these sites were in paleo-floodplains where elephants and hippos were intensively active and formed natural taphocoenoses. Examples of this abound in the paleobiological record. For example, at BK (upper Bed II, Olduvai Gorge, Tanzania), some *Pelorovis* carcasses were naturally deposited on the same alluvial bar and among the fauna processed by hominins, thereby creating an artificial spatial concurrence between independent depositional events and agencies (Organista et al., 2016). In the Middle Pleistocene deposits of Torralba and Ambrona (Spain), artifact-littered areas occur in spatial association with naturally-deposited elephant carcasses, only a few of which were also exploited to a limited degree by hominins, as demonstrated by a few proboscidean remains bearing cut marks (Villa, 1990; Villa et al., 2005). Post-depositional/sedimentation processes also might have artificially created associations of faunal remains and unrelated lithic artifacts (Villa, 1990; Villa et al., 2005).

Direct taphonomic evidence of hominin exploitation of megafauna^1^ (in the form of cutmarked or dynamically-percussed bone) has been so far the main argument to establish a causal link between hominin agency and megafaunal remains (Tappen et al., 2022). If restricted to the Early Pleistocene, most sites containing partial carcasses of megafauna (e.g., elephant, hippopotamus) either show poor preservation, do not exhibit any of these anthropogenic traces, or these modifications cannot be securely identified (Delagnes et al., 2006; Domínguez-Rodrigo et al., 2007; Haynes, 2022, 1991; Isaac and Isaac, 2023; Leakey, 1971). In contrast, for the Middle and Upper Pleistocene, there is more abundant evidence of hominin butchery of megafaunal remains, especially of elephants (Gaudzinski-Windheuser et al., 2023a, 2023b; Haynes, 2022; Yravedra et al., 2010; Rabinovich et al., 2012); see review of taphonomic evidence in Agam and Barkai, (2018), Haynes (2022) and Konidaris et al. (2021). The question that we address here is if such a behavior has its roots in the early Pleistocene.

Once hominin agency is empirically demonstrated at taphonomically-supported butchery sites, a second inferential step is attributing carcass butchery to hunting or scavenging. Occasionally, the occurrence of embedded flint in proboscidean remains has been taken as direct evidence of predatory behaviors (Agam and Barkai, 2018). This evidence is marginal and chronologically restricted to the late Upper Pleistocene with composite hafted tools. Recently, a supportive argument used to detect proboscidean hunting lies on the intensity of cut marks on carcasses and mortality profiles dominated by male prime adults (Gaudzinski-Windheuser et al., 2023a, 2023b). Prime adult mortality is documented also in other natural palimpsests (Haynes, 1991), in some cases probably caused by fights leading also to broken tusks (Villa and d’Errico, 2001).

Ethnoarchaeological observation of modern foragers butchering elephants and hippos shows that they leave very few cut marks after bulk defleshing (Crader, 1983). Modern elephant butchery after culling events (based on a sample of >500 carcasses) also shows that bulk defleshing virtually leaves no traces on bones (Haynes and Krasinski, 2021). Only butchery aiming at the extraction of meat scraps after bulk defleshing imparts most of the cut marks documented experimentally (Haynes and Krasinski, 2021). This casts doubts on whether cutmarking intensity relates to hunting, scavenging or a maximization strategy feasible in all scenarios. To complicate things more, there is a clear mismatch between the butchery intensity documented at several sites and the resulting tool kit associated with it. Some Middle Pleistocene elephant carcasses display abundant cut marks, but there are very few or no stone tools associated with them (Gaudzinski-Windheuser et al., 2023a); whereas at other sites, elephant carcasses exhibit not a single cut mark but they occur with hundreds or thousands of associated tools (Chavaillon et al., 1987; Delagnes et al., 2006; Yravedra et al., 2010) (Table 1). In the latter case, it is probable that in some cases the bulk of tools may have been used to process other non-megafaunal fauna. In the particular case of the Nadunǵa elephant (Kenya, 1-0.7 Ma), that explanation for the presence of lithics is improbable since the only other mammal carcass remains found are from one bovid and one suid, most likely a background scatter associated with the bird, fish and reptile remains with which they are spatially associated. The question remains as to why such a divergent range of tool sets (both in quantity and typological composition) accompany similar partial megafaunal skeletal remains if they are truly functionally related to them.

**Table 1.**
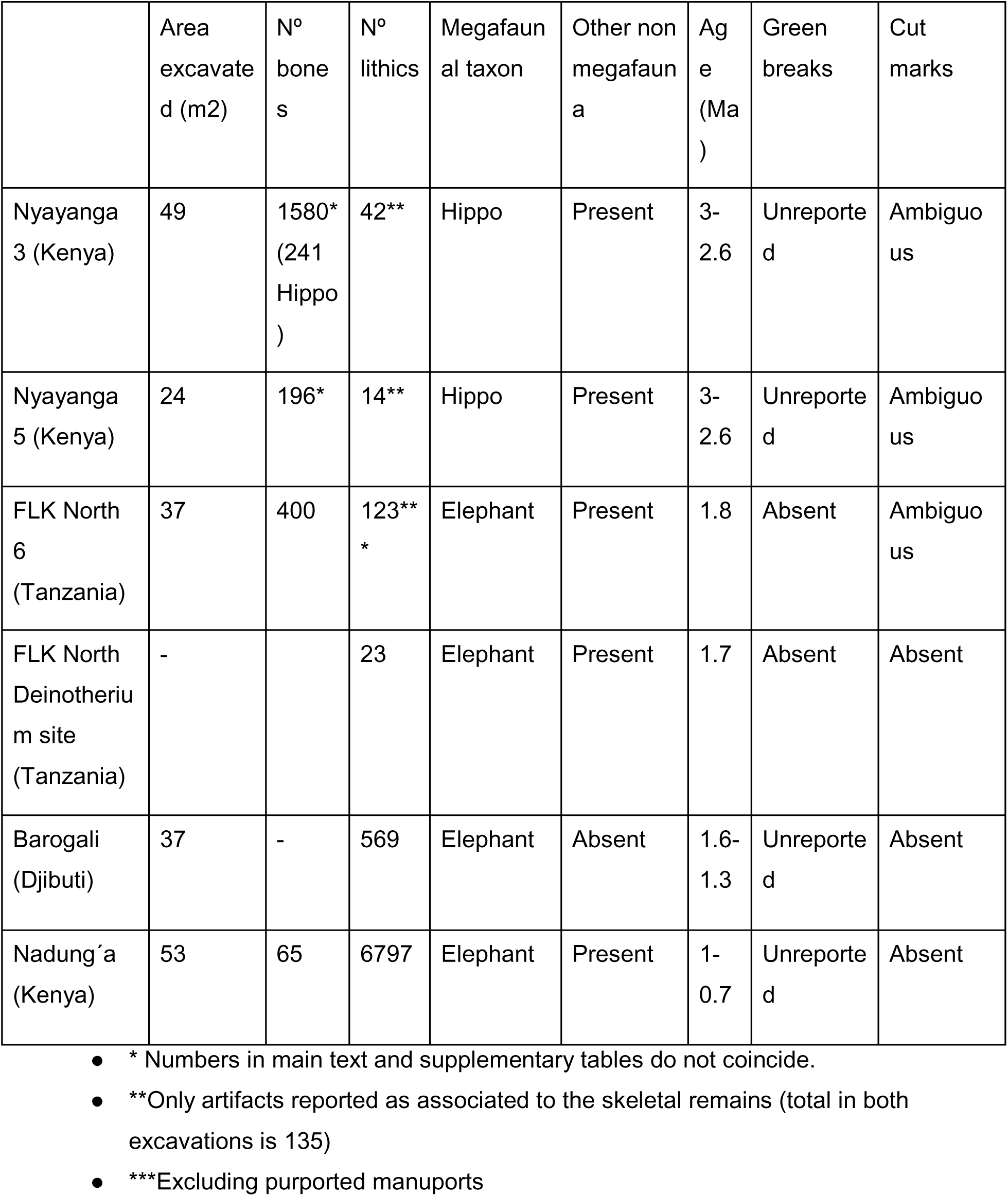
African Early Pleistocene archaeological sites with proboscidean carcasses containing potential evidence of hominin involvement.

To make interpretations even more challenging, many butchery sites exhibit fairly complete and (semi)articulated carcasses with limb bones intact. In some of these instances, elements were dismembered but not demarrowed or degreased (Gaudzinski-Windheuser et al., 2023a). Defleshing or demarrowing/degreasing do not require disarticulating bones, and the energy costs involved in cutting through tendons and ligaments of a proboscidean carcass are better explained as a maximizing strategy aiming at reducing the costs of element transportation (neither defleshing or demarrowing/degreasing requires disarticulation), and/or further exploitation of within-bone fat resources. None of this is clearly documented in those sites. Fat was a crucial resource during human evolution (Ben-Dor and Barkai, 2024 ; Ben-Dor et al., 2011; Reshef and Barkai, 2015; Solodenko et al., 2015), and the large fat deposits stored in proboscidean carcasses rationally would have been a target of prehistoric butchery, especially in strategies of carcass maximization (as documented in the abundant cut mark record at some megafaunal butchery sites). The question of why our ancestors chose to maximize flesh extraction from proboscideans, and yet frequently abandoned their fat deposits untouched remains unanswered. Modern Bisa people (Zambia) butcher elephants and do not exploit their fat, but they follow a “satisficing” strategy (Haynes and Krasinski, 2021), through bulk defleshing (barely impacting bone surfaces) (Crader, 1983), and not a “maximizing” strategy, as suggested for some highly anthropogenically-impacted archaeological proboscidean sites (Gaudzinski-Windheuser et al., 2023a). It could be argued that the meat of healthy elephants contains already a high amount of fat and that other anatomical areas (like the podal pads) contain extremely nutritious fat, or that the relevance of fat exploitation varies according to latitude and climate, but if the strategy was to maximize food resources, long bone medullary contents should also have been consumed (Ben-Dor et al., 2011; Haynes et al., 2021), as documented in sites like Castel di Guido (Italy), Preresa (Spain) (Boschian and Saccà, 2015; Yravedra et al., 2012), or BK (see below). Megafaunal long bone green breakage (linked to continuous spiral fractures on thick cortical bone accompanied by modification [in some cases, intensive] of cortical and medullary surfaces of breakage planes) is probably a less ambiguous trace of butchery than “cut marks”, since many of the latter could be equifinal and harder to identify in contexts of moderate to poor preservation, especially in contexts of high abrasion and trampling (Haynes et al., 2021, 2020). For example, the purported cut-marked pelvis from Fuentenueva 3 (Yravedra el al. 2024) exhibits intense biostratinomic -in addition to diagenetic-modifications impacted by abrasive processes, which renders the pristine experimental reference collection inadequate to confidently classify it as human-made. This is further supported by the fact that the experimental trampling collection used depicts trampling marks as extremely shallow and with wide divergent walls. This does not include the large diversity of morphologies generated by trampling marks, whose cross-sections frequently are indistinguishable from stone-tool cut marks (Domínguez-Rodrigo et al., 2010b). Given the abrasive sedimentary context, the abundance of abrasion marks on the affected specimens, and the lack of description of microscopic features that could unambiguously be used to identify those traces as human-made, we remain skeptical of their anthropogenic nature.

Ideally, cut marks should complement green breakage patterns to identify butchery episodes, but the virtual lack of cut marks in “satisfying” proboscidean exploitation renders their presence/absence non-indicative of the type of anthropic action. Despite the occurrence of green fractures on naturally-broken bones, such as those trampled by elephants (Haynes et al., 2020), and those occurring through traumatic fracturing or gnawed by carnivores (Haynes and Hutson, 2020), these fail to reproduce the elongated, extensive, or helicoidal spiral fractures (uninterrupted by stepped sections), accompanied by the overlapping conchoidal scars (both cortical and medullary), the reflected scarring, the inflection points, or the impact hackled break surfaces and flakes typical of dynamic percussive breakage. Evidence of this type of green breakage had not been documented earlier for the Early Pleistocene proboscidean or hippopotamid carcasses, beyond the documentation of flaked bone with the purpose of elaboration of bone tools (Backwell and d’Errico, 2004; Pante et al., 2020; Sano et al., 2020). Such type of dynamic breaking, in contrast, has been documented in *Pelorovis* and *Sivatherium* carcasses at the 1.3 Ma site of BK (Olduvai Gorge) (Domínguez-Rodrigo et al., 2014a; Labrado, 2017; Organista et al., 2016). One reviewer suggested that such variables have also been documented in non-anthropogenic contexts, including helicoidal spiral fractures attributed to trampling or carnivore activity (Haynes 1983), adjacent or flake-like scars produced by carnivore gnawing (Villa and Bartram 1996), hackled fracture surfaces allegedly resulting from heavy passive breakage such as trampling or sediment pressure (Haynes 1983), and “impact-like” bone flakes documented in carnivore-broken bones. However, this interpretation is epistemologically problematic because it does not satisfy the fundamental criteria for valid analogy as outlined by Bunge (1981), namely substantial, structural, and environmental affinity. Specifically, the cited examples involve agents, materials, and contexts that differ markedly in composition, mechanical properties, and loading regimes from those considered here. Experimental and actualistic studies demonstrate that carnivores—rather than trampling—are also capable of producing spiral fractures and overlapping bone scarring, but these observations are restricted to faunal remains of substantially smaller body size than elephants, which carnivores can gnaw (Haynes 1983; see also Figures S30–S36). To date, no carnivore has been documented as producing comparable fracture morphologies or surface damage on late juvenile and adult elephant bones. Consequently, the proposed analogy is not supported. Moreover, Haynes (1983) provides no empirical evidence that sediment pressure or trampling can generate hackled fracture surfaces. Such features are instead associated with dynamic loading conditions, whereas passive breakage processes have not been shown to produce these types of modifications (Lyman, 2006). This reasoning also applies to impact flakes on elephant limb bones, which can only be produced by the sole modern agent documented to dynamically fracture non-infantile green proboscidean long bones: humans.

Regardless of the type of butchery evidence -and with the taphonomic caveat that no unambiguous evidence exists to confirm that megafaunal carcasses were hunted or scavenged other than hominins accessed them in different taphonomically-defined stages (i.e., early or late)– the principal reasons for exploring megafaunal consumption in early human evolution is its origin, its episodic or temporally-patterned occurrence, its impact on hominin adaptation to certain landscapes, and its reflection on hominin group size and site functionality. If hominins actively sought the exploitation of megafauna, especially if targeting early stages of carcass consumption, the recovery of an apparent surplus of resources reflects a substantially different behavior from the small-group/small-site pattern documented at several earlier Oldowan anthropogenic sites (Domínguez-Rodrigo et al., 2019) -or some modern foragers, like the Hadza, who only exploit megafaunal carcasses very sporadically, mostly upon opportunistic encounters (Marlowe, 2010; O’Connell et al., 1992; Wood, 2010; Wood and Marlowe, 2013). Determining when the process of becoming megafaunal commensal started has major implications for human evolution, since it has clear implications for our understanding of past social group sizes and social dynamics.

The multiple taphonomic biases intervening in the palimpsestic nature of most of these butchery sites often prevent the detection of the causal traces linking megafaunal carcasses and hominins. Functional links have commonly been assumed through the spatial concurrence of tools and carcass remains; however, this perception may be utterly unjustified as we argued above. Functional association of both archaeological elements can more securely be detected through objective spatial statistical methods. This has been argued to be foundational for heuristic interpretations of proboscidean butchery sites (Giusti, 2021). Such an approach removes ambiguity and provides a better argument for spatial functional association, as demonstrated at sites like Marathousa 1 (Konidaris et al., 2018) or TK *Sivatherium (Panera et al., 2019)*. This method will play a major role in the present study.

Here, we present the discovery of a new elephant butchery site (Emiliano Aguirre Korongo, EAK), dated to 1.78 Ma, from the base of Bed II at Olduvai Gorge (Tanzania). It is potentially the oldest unambiguous proboscidean butchery site at Olduvai. In combination with its study, we have carried out extensive survey at a landscape scale on one stratigraphic section of Bed I (Oldowan) and three stratigraphic sections of Bed II: lower Bed II-Lower Augitic Sands (1.8.1.69 Ma), upper middle Bed II (from tuff IIB to IID, 1.5 Ma), and upper Bed II (overlying Tuff IID, 1.3 Ma), which are known to contain Acheulian materials. The goal was to find megafaunal fossils bearing traces of hominin modification across the landscape. Starting at 1.78 Ma, Olduvai Gorge records extensive evidence of hominin involvement with megafaunal carcasses (Domínguez-Rodrigo et al., 2014b, 2009b), namely Proboscidea and Hippopotamidae, followed closely by Giraffidae and big Bovidae. The nature of this interaction is elaborated and analyzed in this paper. We have used a spatial approach, combined with a more general taphonomic analysis, and an analysis of hammerstone-broken bone specimens occurring widespread on the three targeted Bed II stratigraphic intervals. Also provided is a technological study of the lithic artifacts.

## Results

### The Emiliano Aguirre Korongo (EAK) site

EAK is located in the area of the confluence of the two gorges, next to geo-locality 45a, between the FLK-N and FLK-NN sites (Hay, 1976) (Fig. 1). Stratigraphically, the archaeological remains rest on Tuff IF (1.78 M.a), i.e. at the beginning of Bed II (Deino, 2012). As in the whole area of the gorge, the lowermost Bed II is similar to the uppermost Bed I, which is composed of clays deposited in the margins of the lake. The drying period that characterizes Bed II had not yet taken place during this moment. The sedimentary context of the Olduvai basin in this area is dominated by low energy processes, with predominantly clay deposition, controlled by small seasonal rises and falls in lake level. The lake had a very high pH and high salinity in the central zone (Deocampo et al., 2017), but this value is lowered at the margins by the arrival of runoff water. EAK is located roughly coinciding with the FLK fault zone (Fig. 1). The bone and lithic remains lie on a thin layer of clay (<5 cm) on top of tuff IF. They are also covered by the lacustrine clay of the lowermost Bed II. It is therefore the oldest documented archaeological site formed in Bed II in Olduvai.

**Figure 1.**
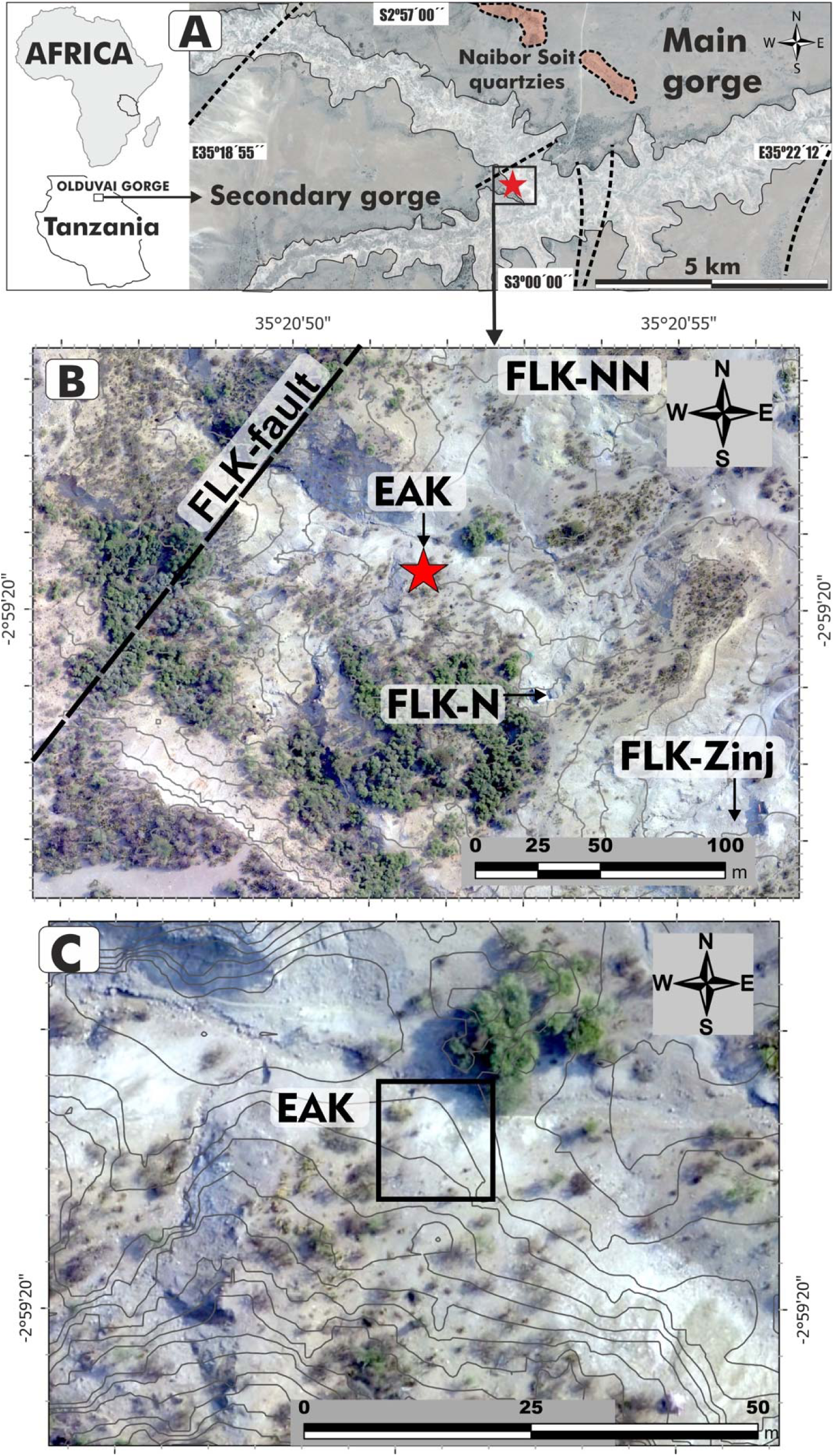
Location of EAK in the junction of the main and secondary branches of Olduvai Gorge, main faults and Naibor Soit quartzites (A), within the area where the Bed I sites cluster at the junction (B), and the specific EAK locus with 1 m contour lines (C).

During the recent Holocene, the EAK site was affected by a small landslide which displaced the site 6.25 m vertically and approximately 12 m downslope. Although the stratigraphic sequence remains clearly recognizable, Tuff I-F, which serves as the paleosurface of the site, is fragmented into numerous blocks up to 1 m in diameter (Fig. 2). The archaeological deposit has moved together with the blocks and their original spatial distribution has been altered by this post-sedimentary deformation.

**Figure 2.**
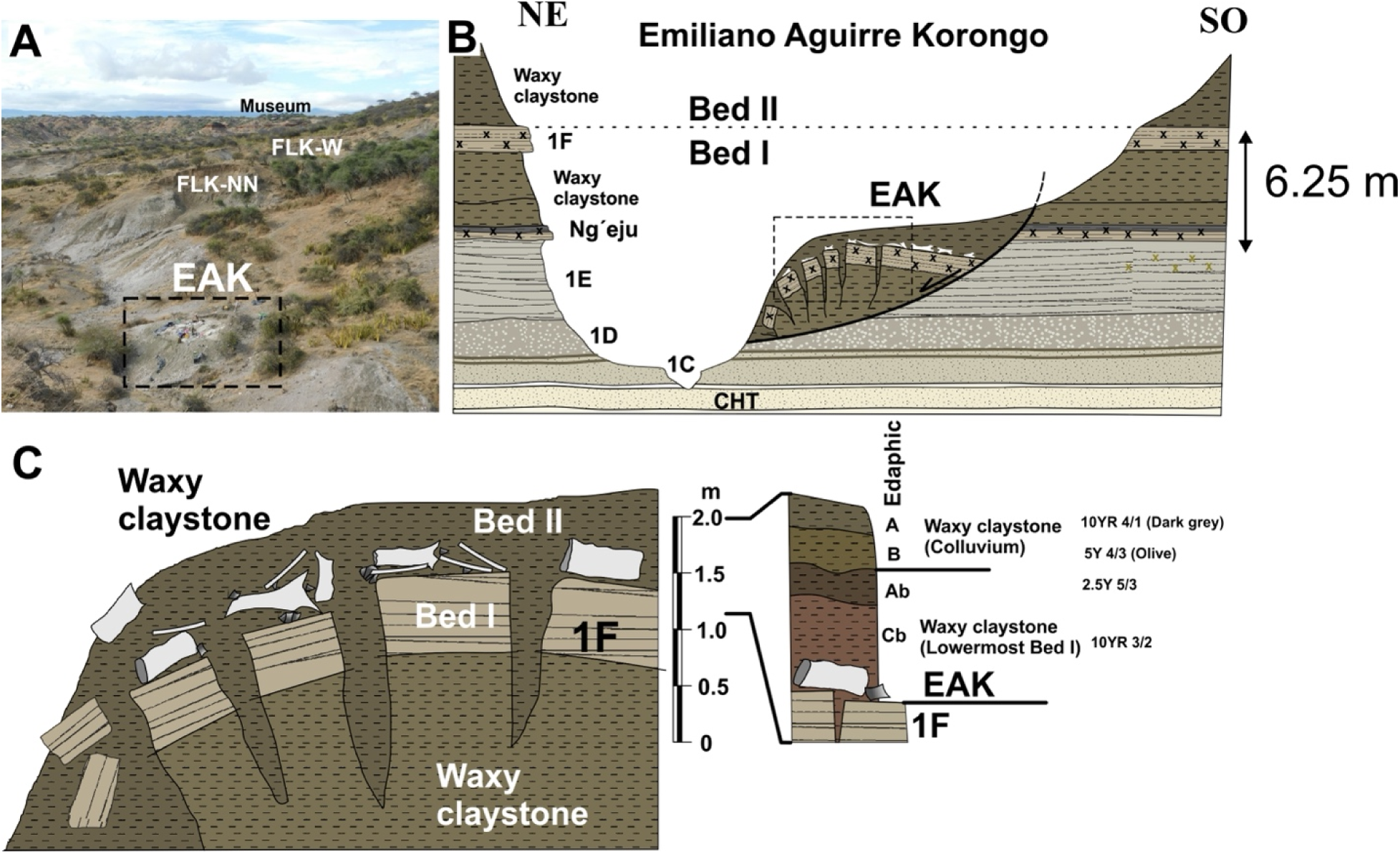
A, Location of EAK on the south side of the Main Gorge (view to the east). B, Stratigraphic section of the gully (Korongo) showing the correlation between the different marker tuffs of Bed I on either side. The landslide displaced the boundary between Bed I and Bed II, including the following sequence: waxy claystone, tuff 1F, waxy claystone. Vertical displacement is 6.25 m. C, detail of the stratigraphy of the site, showing how the archaeological level rests directly on Tuff 1F and was covered by the clay of the lowermost Bed II. The landslide has created vertical cracks that separate both the Tuff 1F and the archaeological level itself. The archaeological remains have moved along with the tuff blocks. The original clay of the lowermost layer II protected the fossils from erosion after the landslide. An incipient soil formed on this surface and was subsequently buried by colluvium. At present, erosion is acting on the part closest to the stream, affecting some of the fossils.

Olduvai Gorge is located in a sedimentary basin bounded to the south by several volcanoes, and to the north by large outcrops of metamorphic rocks. These two groups are the sources of raw material (basalts and quartzites respectively) during practically the entire Pleistocene at Olduvai. Both, the aforementioned basement rocks and the Plio-Quaternary and Quaternary sedimentary fill are affected by these major east-west faults known as 1^st^, 2^nd^, 3^rd^, 4^th^, KK, FLK and 5^th^ faults. It should be noted that the FLK fault can only be identified in the landscape through the deformation of the southern escarpment of the gorge, at the beginning of the gully that gives its name to this deposit (EAK). It is a well-known fault because it defines the FLK-5 fault block, described stratigraphically by Hay (1976). Although it is an escarpment degraded by the current geodynamics, it affects the most recent aeolian deposits of Olduvai (Naisiusiu). This geomorphological feature indicates that the FLK fault remained active at least until the Upper Pleistocene to Holocene. The exact direction of the fault is unknown, although the stratigraphy suggests that it must be SW-NE (Fig. 1), and its vertical drop is several tens of meters. The movement of the fault has probably triggered local earthquakes of varying magnitude. It is well known that earthquakes are one of the main causes of landslides (Keefer, 1984; Ugai et al., 2012; Wasowski et al., 2011), and in this particular case, they may have triggered several landslides affecting deposits from the base of the present gully -i.e. from the upper part of Bed I to the uppermost Bed (Fig. 2). The fluvial erosion of the stream at the base of the slopes also destabilized the base of the slopes in this gully, favoring this type of gravitational process. In the case of EAK, this is a very small landslide, although much larger landslides have been recorded in the surrounding area, affecting Bed II units such as the Lower Augitic Sandstone (LAS), which slides over the waxy clay of the lowermost Bed II. Interestingly, there is a similar archaeological site on the Colorado Plateau in northern New Mexico (USA); in this case, a mammoth presenting a similar post-depositional history (Rowe et al., 2022).

In June 2022, a partial elephant carcass was found at EAK on a fragmented stratigraphic block situated topographically below FLK North, containing most of the sequence from Tuff ID to the carbonated root clays overlying Tuff IF that constitute the base of Bed II. The carcass must have been eroding over the years, since proboscidean bone fragments and lithics had been found over more than 100 m repeatedly along the seasonally-active stream at the bottom of the gulley. In January of the same year, we had not documented anything eroding on the same block, but after the rainy season, several bones emerged, making it obvious that the site contained remains of a proboscidean carcass (Fig. 3). Excavation started right after the discovery, and uncovered remains from a single *E. recki* juvenile individual. We initially set up a 4×2 m trench, followed by an extension consisting of two additional trenches (Figs. 3). The northern side presented an erosive front, probably responsible for several of the scattered elephant bones found along the bottom gully. The remaining excavated partial carcass contained the complete pelvis associated with both hindlimbs (Table 2). These included both femora and tibiae, plus one set of tarsal elements (including calcaneus, astragalus) and a fully articulated foot (Figure S1). In addition, several rib fragments belonging to a minimum of 8 ribs, as well as a portion of an ulna were also discovered (Fig. S2). It is interesting to note that no complete vertebrae were discovered (only one fragment was identified), despite the presence of a portion of the axial skeleton. The skull was positioned upside down and the projecting tusks were partially preserved (Fig. 3). In addition to the elephant carcass, a background scatter of few bones, including a *Parmularius* M^2^, a vertebra, rib, tibia and metatarsal of a small-sized artiodactyl were found.

**Figure 3.**
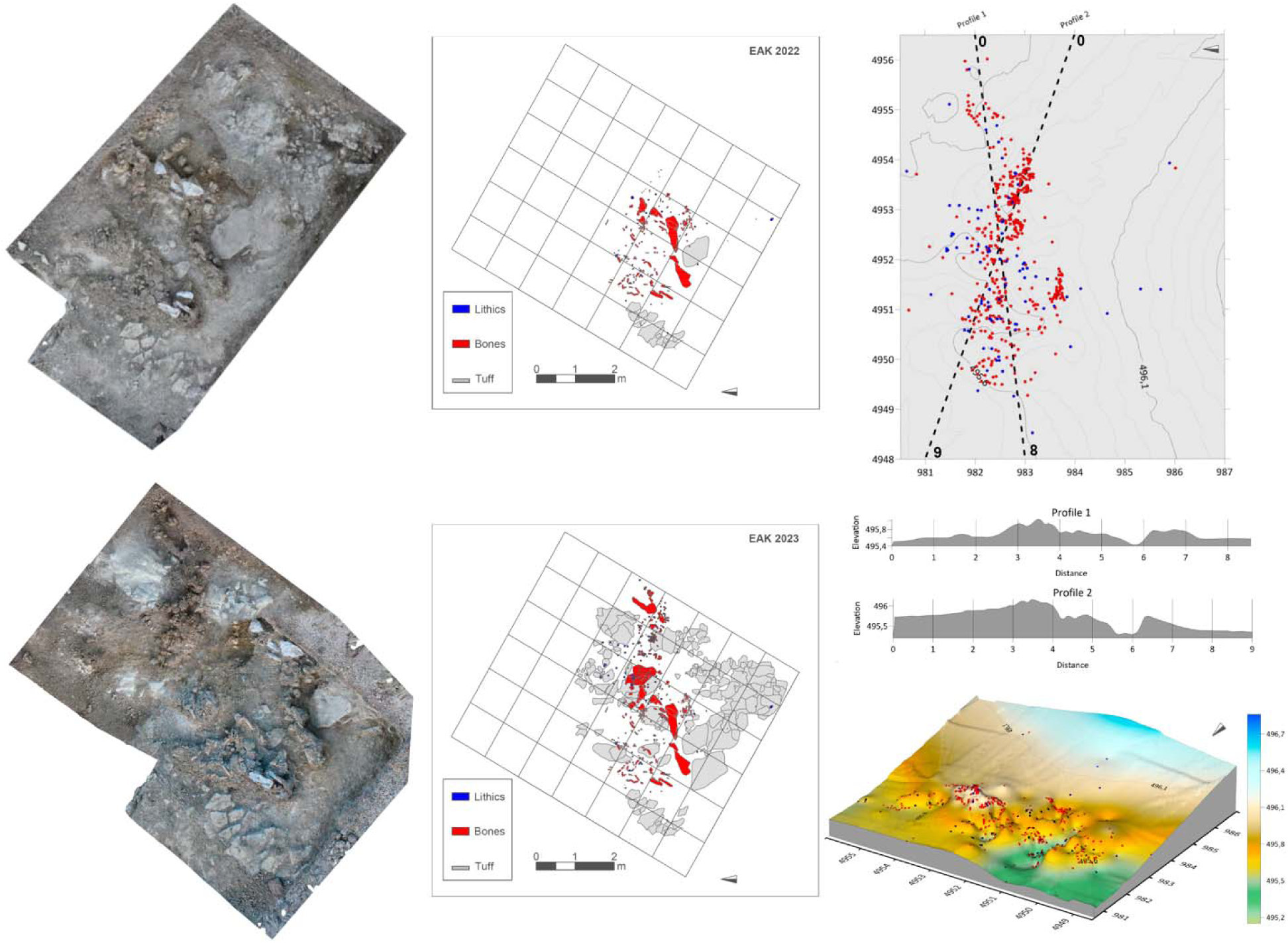
Photogrammetry (left) and planimetry (right) of the EAK site in 2022 and 2023. The site-wide distribution of materials, lithics (blue) and bones (red), topographic profiles, and the 3D terrain model are shown.

A total of 80 stone tools, mostly flakes and flake fragments, were spatially distributed in tight connection with 153 bone remains, pertaining to 46 elements of a proboscidean carcass (Fig. 3; Table S1). Their preservation is exceptional and the edges remain sharp. The close spatial association must be functionally linked. Although the cortical bone in several elements was affected by diagenesis, preventing the preservation of any potential bone surface modifications, the good preservation in the remaining assemblage did not yield any marks created either by hominins or other carnivores. On-going restoration work will provide an accurate estimate of well-preserved and modified fractions of the assemblage. Two elephant bone specimens showed attributes of green breakage, inferred to be of anthropic origin (See Supplementary Information). The tibiae were well preserved, but the epiphyseal portions of the femora were missing, probably removed by carnivores, which would also explain why a large portion of the rib cage and almost all vertebrae are missing. Further support for this likelihood can be found in the deletion of bones from three feet, which are one of the richest sources of fat and affected early in the process of carnivore ravaging (Haynes, 1991, 1988; Haynes and Hutson, 2020). The absence of the forelimbs could be also explained if carnivores had dragged them away from the core axial area; however, the fact that the carcass was detached from its original depositional locus and gravitationally moved (with its overlying sediment) after faulting might also explain that the missing carcass parts could still be in their original depositional spot. Our testing of the area where the block was supposed to have been originally formed yielded no results, mainly because of the thick overburden of Ndutu sediments in the area. The fact that the carcass was moved while encased in its sedimentary context, along with the close association of stone tools with the elephant bones, is in agreement with the inference that the animal was butchered by hominins. A more objective way to assess this association is through spatial statistical analysis.

### The spatial statistical analyses of EAK

The analysis of orientation (azimuth) and slope (plunge) (Fig. S28) has shown that the null hypothesis of spatial isotropy cannot be rejected, whether the entire sample is analysed jointly, or the elephant remains are differentiated from the lithic ones. This result is supported using Rayleigh, Kuiper, and Watson tests, but also by the visualization of the stereograms and rose diagrams, which indeed show the existence of orientations in all directions. Regarding the slope, the stereograms indicate the absence of strong imbrications of remains.

The Spatial Point Pattern (SPP) of the elephant bones has an intensity of 7.74 specimens per square meter. However, this intensity varies significantly across the analyzed surface, with a very high-intensity area in the center of the excavated area and very low intensity surrounding this zone. This pattern is evident both when analyzing the intensity itself and when examining the hot spot maps (Fig. S24 a-c). The use of MAD and DCLF tests has rejected the null hypothesis of spatial randomness with the inhomogeneous *K* and *L* functions, as well as with the *K*scaled and *L*scaled functions (p-value = 0.025), while it could not be rejected with the *F*, *G*, and *J* functions (p-value = 1). These results align with the use of spatial functions (Fig. S24 d-k), as the *F*, *G*, and *J* functions do not reject the random model, whereas the *K*, *L*, *K*scaled, and *L*scaled functions reject randomness and suggest a clustered SPP. Additionally, the inhomogeneous *K* and *L* functions indicate that this clustering occurs only at short distances and that at distances greater than 1 meter, the SPP follows a regular pattern. The pair correlation function also appears to indicate the presence of a clustered pattern at very short distances. According to the nearest-neighbor cleaning test, most of the faunal assemblage is highly likely to be part of the cluster (Fig. S24 l-n).

When performing different regression models, we found that the linear regression had an AIC of −861.53, the quadratic regression −1990.689, the cubic regression −2076.40, and the 4^th^-degree polynomial regression −2100.39. However, we used the quadratic regression for the analysis because the cubic and 4^th^-degree polynomial regressions were found to be unreliable due to issues related to sample size and collinearity. Therefore, we performed modelling using the quadratic regression (Fig. S24 o-r), reinforcing the interpretation of a very high-intensity cluster in the central part of the excavated area. Furthermore, it is proposed that the accumulation should not extend significantly beyond the excavated area. This supports that site erosion may have affected the preserved concentration less than initially suggested.

The analysis of the SPP for lithic remains shows an intensity of 1.67. Although the sample size is much smaller, it exhibits a pattern similar to that of the bone remains. It also presents intensity and hot spot maps (Fig. S25 a-c), with very high intensity in the central area of the spatial window and very low intensity in the surrounding areas. The DCLF and MAD tests yield the same results as those for the elephant remains. However, the functions (Fig. S25 d-k) show some differences. The *F*, *G*, *J*, and pair correlation functions do not reject the null hypothesis, while the *K*, *L*, *K*scaled, and *L*scaled functions do reject it. In this case, the inhomogeneous functions suggest the presence of a regular pattern, while the scaled functions indicate a clustered pattern. Once again, the nearest-neighbor cleaning test (Fig. S25 l-n) suggests that the lithic remains in the high-intensity area are highly likely to be part of a cluster.

The regression analyses in this case show much higher AIC values compared to the faunal remains. However, as before, the 4^th^-degree polynomial regression has the lowest AIC (−87.99), followed by the cubic regression (−83.02), the quadratic regression (−71.73), and finally the linear regression (79.5). However, once again, the cubic and 4^th^-degree polynomial regressions cannot be used due to collinearity and sample size issues, making the quadratic regression the most appropriate choice. Modelling with quadratic regression (Fig. S25 o-r) further reinforces the presence of a large cluster and the possible absence of remains outside the analysed window. Interestingly, the area of maximum intensity does not exactly coincide with that of the elephant remains but is slightly shifted to the left (west) within the spatial window.

The analysis of the SPP with marks (elephant bones vs. lithic tools) shows that, as expected due to the imbalance in sample size, there is always a higher probability of finding elephant remains than lithic remains (Fig. 4 a), with the only exception being at the far left (north) of the spatial window. The analysis of spatial marks reveals a significant segregate correlation between fauna and lithics when using the *K*cross and *L*cross functions (Fig. 4 b-c). The pattern is of regular segregation, meaning that although lithics and bones form part of the same cluster, they tend to occupy different positions, with lithics clustered in the periphery, as would correspond to their use and discard upon the processing of the elephant carcass, which occupies the central part of the cluster.

**Figure 4.**
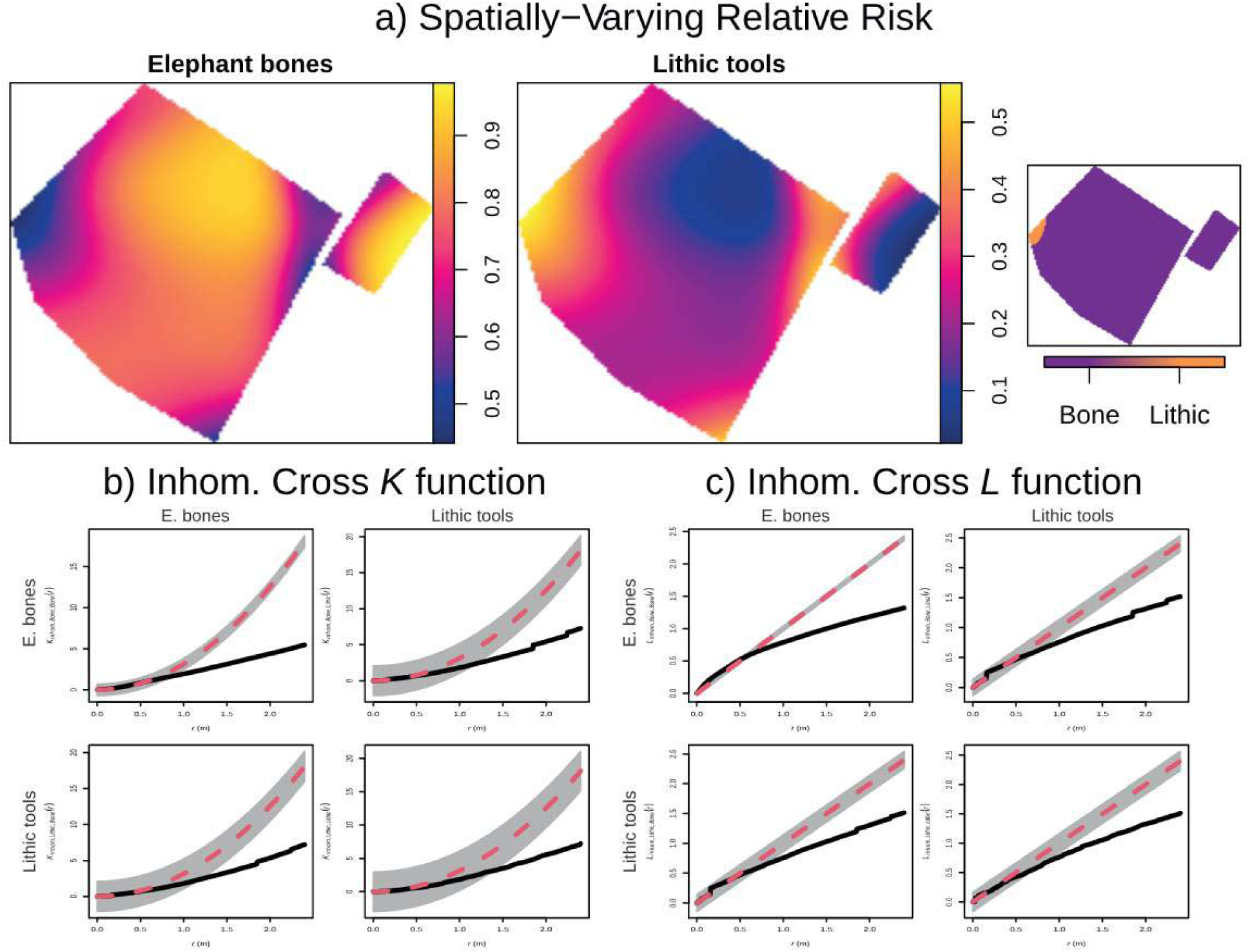
Relative risk (i.e., probability of occurrence) of bones and lithic artifacts at EAK, with second-order functions showing regular correlation between both types of items.

Finally, analyzing the bones as a covariate has further reinforced the presence of high lithic tool intensity in the center of the spatial window and the relation between fauna and lithic artifacts (Fig. 5). First, modelling intensity based on the covariate and the estimated intensity (ρ) shows that intensity is very high when lithic remains are close to elephant remains, but it drops sharply when moving just 50 cm away. This finding is further supported by the ANOVA test on the model (p-value = <2.2×10LJ¹C) and the results of the Berman Z_1_ (p-value = <8.456×10LJC) and Z_2_ (p-value = <1.585×10LJ³C) tests. This also shows that the functional connection between the elephant bones and the tools has been maintained despite the block post-sedimentary movement. This is further supported by the technological analysis (Supplementary Information).

**Figure 5.**
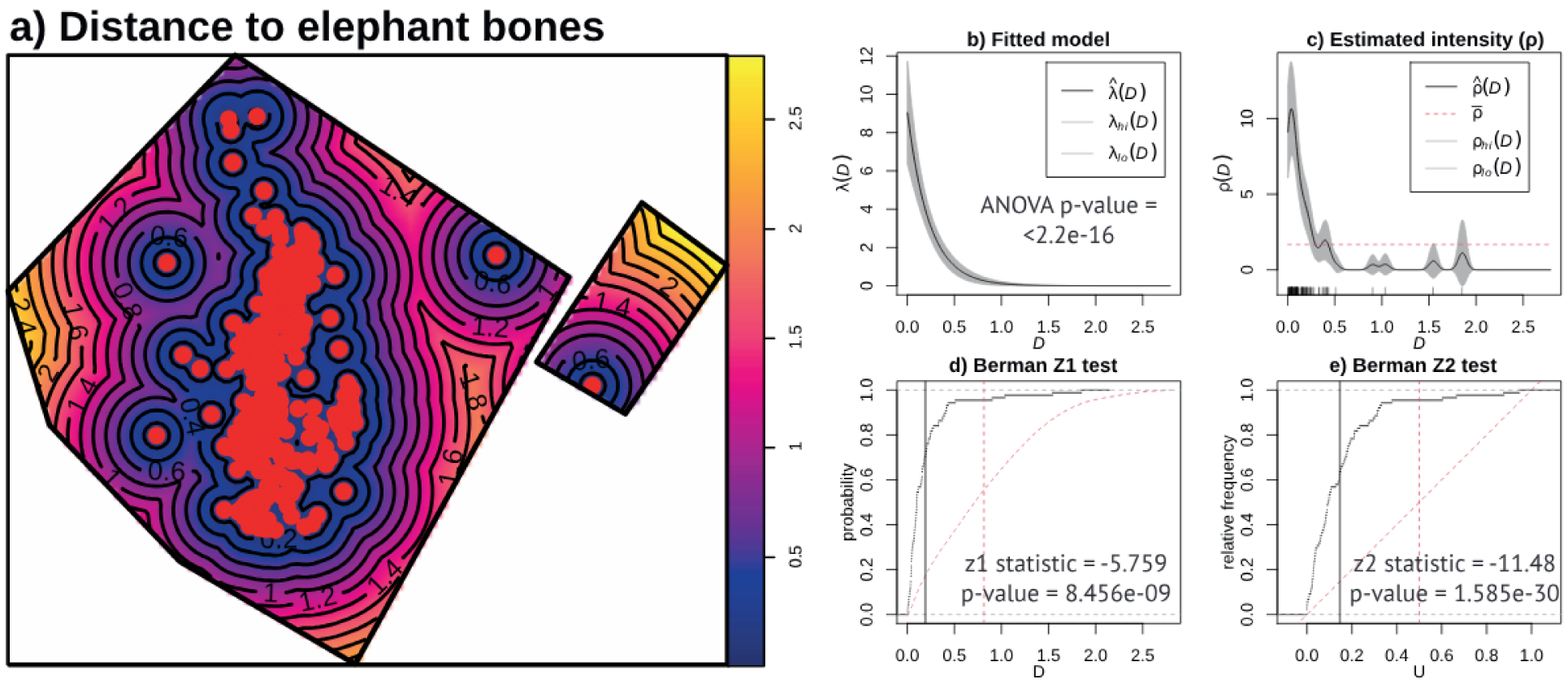
Correlation between lithics and bones at EAK (using the elephant bones as a covariate) with the ρ (‘rhohat’) function and a Z 1 and Z 2Berman–Lawson–Waller.

### Traces of megafaunal exploitation along the paleolandscape at Olduvai Gorge

#### Bed I

During our survey of the Bed I targeted strata in 2024, no significant traces of megafaunal green bone breakage were observed, with the exception of one locality (Fig. S3). In previous years, we had observed few megafaunal fossils, mostly restricted to hippopotamus scattered remains. They were not particularly abundant, despite the clearly lacustrine and alluvial depositional settings sampled at the junction.

#### Bed II

Out of the three targeted units (lower, intermediate and upper), the lower unit yielded the highest concentration of elephant and hippopotamus remains, followed by the upper unit (Figure S3). The distribution shows a clustered pattern, despite the virtual continuity of most of the sequence in larger areas. The junction area contains the highest density or megafaunal remains for the lower unit. Figure S3 shows the thorough overlap of those elephant and hippopotamus carcasses with areas containing high density of stone artifacts. All the carcasses documented in the lower unit but one (EAK) occur within the Lower Augitic Sands (LAS). This unit contains a landscape-scale concentration of lithic artifacts spreading from HWKE to FLK West (Domínguez-Rodrigo et al., 2023; Uribelarrea et al., 2019, 2017).

The distribution of carcasses in the intermediate unit is less dense and mostly restricted to a small portion of the junction. This small area also shows concentration of lithic remains, but its extent is still unknown since only two localities have been excavated [one to the west of FLK West (Fujioka et al., 2022) and BBS] and no much surface material is found that could be securely attributed to this unit in between both sites. BBS (Bob Brain Site) is a recent site discovered in middle Bed II on the same stratigraphic unit as the site named by Fujioka et al. (2022) as FLK West, recently renamed T69 Complex. In the side gorge, SHK represents this unit, with well documented presence of megafauna and cut-marked hippopotamus remains (Diez-Martín et al., 2017; Domínguez-Rodrigo et al., 2014b).

The other large cluster of elephant and hippopotamus carcasses is documented in the upper section of the side gorge (Fig S3). There, most of them concentrate around two sites (BK and SC). These sites are also relevant because they contain other smaller megafauna, such as *Sivatherium* and *Pelorovis* in larger numbers (Domínguez-Rodrigo et al., 2014a; Organista et al., 2019, 2017, 2016). It is at these two sites and the landscape surrounding them that the highest densities of lithic artifacts are also found in the upper section of Bed II (Diez-Martín et al., 2009).

The clustering of the elephant (and hippopotamus) carcasses in the areas containing the highest densities of landscape surface artifacts is suggestive of a hominin agency in at least part of their consumption and modification. The presence of green broken elephant long bone elements in the area surveyed is only documented within such clusters, both for lower and upper Bed II. This constitutes inverse negative evidence for natural breaks occurring on those carcasses through natural (i.e., non-hominin) pre- and peri-mortem limb breaking (Haynes et al., 2021, 2020; Haynes and Hutson, 2020). In this latter case, it would be expected for green-broken bones to show a more random landscape distribution, and occur in similar frequencies in areas with intense hominin landscape use (as documented in high density artifact deposition) and those with marginal o non-hominin intervention (mostly devoid of anthropogenic lithic remains).

This indicates that from lower Bed II (1.78 Ma) onwards, there is ample documented evidence of anthropogenic agency in the modification of proboscidean bones across the Olduvai paleolandscapes. The discovery of EAK constitutes in this respect the oldest evidence thereof at the gorge. The taphonomic evidence of dynamic proboscidean bone breaking across time and space supports, therefore, the inferences made by the spatial statistical analyses of bones and lithics at the site.

## Discussion

Evidence shows that the oldest systematic anthropogenic exploitation of proboscidean carcasses is documented (at several paleolandscape scales) in the Late Pleistocene sites of Neumark-Nord (Germany) (Gaudzinski-Windheuser et al., 2023a, 2023b). Evidence of at least episodic access to proboscidean remains goes back in time (see review in Agam and Barkai, 2018; Ben-Dor et al., 2011; Haynes, 2022). Redundant megafaunal exploitation is well documented at some early Pleistocene sites from Olduvai Gorge (Domínguez-Rodrigo et al., 2014a, 2014b; Organista et al., 2019, 2017, 2016). At the very same sites, the stone artifactual assemblages, as well as the site dimensions, are substantially larger than those documented in the Bed I Oldowan sites (Diez-Martín et al., 2024, 2017, 2014, 2009). Whether this reflects larger hominin groups or more prolonged and time-averaged intervention in more productive environments remains to be determined (it will likely vary according to site).

A recent discovery of a couple of hippopotamus partial carcasses at the 3.0-2.6 Ma site of Nyayanga (Kenya), spatially concurrent with stone artifacts, has been argued to be causally linked by the presence of cut marks on some bones (Plummer et al., 2023). The only evidence published thereof is a series of bone surface modifications on a hippo rib and a tibial crest, which we suggest they might also be the result of byproduct of abiotic abrasive processes; the marks contrast noticeably with the well-defined cut marks found on smaller mammal bones in the same localities (Plummer et al.’s 2023: fig. 3C,D) associated with the hippo remains (Plummer et al., 2023). The two trenches from which both hippo remains derive are located in the stratigraphic unit NY1, consisting of a channel containing cobble conglomerates and coarse sands in addition to silts. Gravel and coarse sand can generate the same types of marks seen in the figures of the Nyayanga hippo bones (Domínguez-Rodrigo et al., 2009a; Manuel Domínguez-Rodrigo et al., 2010b). Hippopotamidae are the most abundant taxa in the Nyayanga excavations, indicating a productive taphocoenotic environment, in which hippopotamus remains co-occur with a large diversity of mammal taxa, most of which could have been naturally deposited and mixed. No green fractures have been reported on the hippopotamus bones as found in some other mammal fauna. Although Nyayanga could potentially be one of the earliest examples of megafaunal butchery, the taphonomic evidence remains ambiguous. Butchery experiments show that complete butchery of animals is reflected in a large percentage of remains bearing traces of butchery marks. In T3, these are documented in <0.9% and in T5 in 1%. This is a magnitude of 10-30 time lower than expected using experimental butchery frameworks (even if using only well-preserved bone specimens) (Domínguez-Rodrigo, 1997; Domínguez-Rodrigo et al., 2014; Pizarro-Monzo et al., 2021). The 1.8 Ma *Elephas recki* of FLK North (level 6, Olduvai Bed I) has the potential of being the oldest proboscidean butchery site (Leakey, 1971). It was argued that cut marks existed on its complete bones (Bunn, 1982; Delagnes et al., 2006), but a subsequent taphonomic study of those marks suggested that they might have been made by trampling (Domínguez-Rodrigo et al., 2007). Additionally, it is unclear if the spatial association of stone tools and elephant bones is functional or artificial. Leakey (1971) did not report on the third dimension of her excavations and this is crucial. In our excavations of FLK N6, we documented that the level was a deposit spanning more than half a meter depth and fossils and stone tools were vertically scattered throughout (Domínguez-Rodrigo et al., 2010a). It would have been essential to document that the FLK N6 tools associated with the elephant were either on the same depositional surface as the elephant bones and/or on the same vertical position.

The ambiguity about the FLK N6 elephant renders EAK, potentially, the oldest proboscidean butchery evidence at Olduvai, and also probably one of the oldest in the early Pleistocene elsewhere in Africa. FLK North Deinotherium occupied a position above the EAK stratigraphic level, and evidence of hominin intervention there is missing (Leakey, 1971). A cut-marked bone fragment of a large animal (probably a hippopotamus) was documented at El-Kherba (Algeria) dated to 1.78 Ma (Sahnouni et al., 2013). These findings are suggestive of the beginning of megafaunal exploitation by humans in similar times both in East and North Africa.

Our landscape study of the megafaunal remains at Olduvai show that elephants and hippopotamus overlap on the same clustering areas. In all of them, the geology is indicative of extensive alluvial zones. At the base of Bed II, the gorge junction is characterized by extensive wetlands (Ashley, 2003; Ashley et al., 2009; Deocampo et al., 2002; Hay, 2023; Liutkus and Ashley, 2003). In the overlying LAS, wetlands are crossed by important fluvial systems (Domínguez-Rodrigo et al., 2023; Uribelarrea et al., 2017). In the middle and upper Bed II fluvial and wetland systems intertwined as the lake was reduced and moved away from its former shoreline in the Bed I-Bed II transition (Arráiz et al., 2017; Garrett, 2017; Uribelarrea del Val and Domínguez-Rodrigo, 2017). In all cases, these wet environments must have been preferred places for water-dependent megafauna, like elephants and hippos, and their overlapping ecological niches are reflected in the spatial co-occurrence of their carcasses. Both types of megafauna show traces of hominin use through either cutmarked or percussed bones, green-broken bones or both (Supplementary Information).

Green-broken elephant bones are documented at Olduvai at a landscape scale during the LAS deposition (1.7 Ma) (Diez-Martín et al., 2015; Domínguez-Rodrigo et al., 2023; Uribelarrea et al., 2017). This record is now added to the newly discovered EAK site, showing tight spatial association (both vertically and horizontally) between bones and stone tools. This trend of spatial association of stone artifacts and proboscidean bones continues through the deposition of Bed II. Although no cut marks have been documented on the proboscidean bones that we identified (part of which had cortical surfaces impacted by biostratinomic and diagenetic modification), the presence of green-broken bones at the sites, the butchery of carcasses as indicated by spatial statistical analyses at sites like EAK and TK (Domínguez-Rodrigo et al., 2014a; Organista et al., 2016; Panera et al., 2019), and the presence of hammerstone-broken bones on the surrounding paleo-landscapes of these and other sites, support the occasional exploitation of megafaunal remains by humans. In most green-broken limb bones, we document the presence of a medullary cavity, despite the continuous presence of trabecular bone tissue on its walls. We do not know if the bone breakage was intended to obtain marrow or to create raw material for making artifacts or tools with functional purposes or both. We have been able to identify only one potential bone tool in our sample (Figures 6, S4-S23); the scarcity of bone implements supports the former option. The exploitation of megafauna is not anecdotal, since at sites like BK or TK (upper Bed II), the taphonomic analysis of other megafaunal remains (namely, *Sivatherium* and *Pelorovis*) clearly shows intensive exploitation of complete carcasses in both of these taxa, especially at BK (Domínguez-Rodrigo et al., 2014a; Organista et al., 2016; Panera et al., 2019). The food surplus generated by the exploitation of some of these animals would have been a significant advantage for the survival and adaptation of the hominin groups.

**Figure 6.**
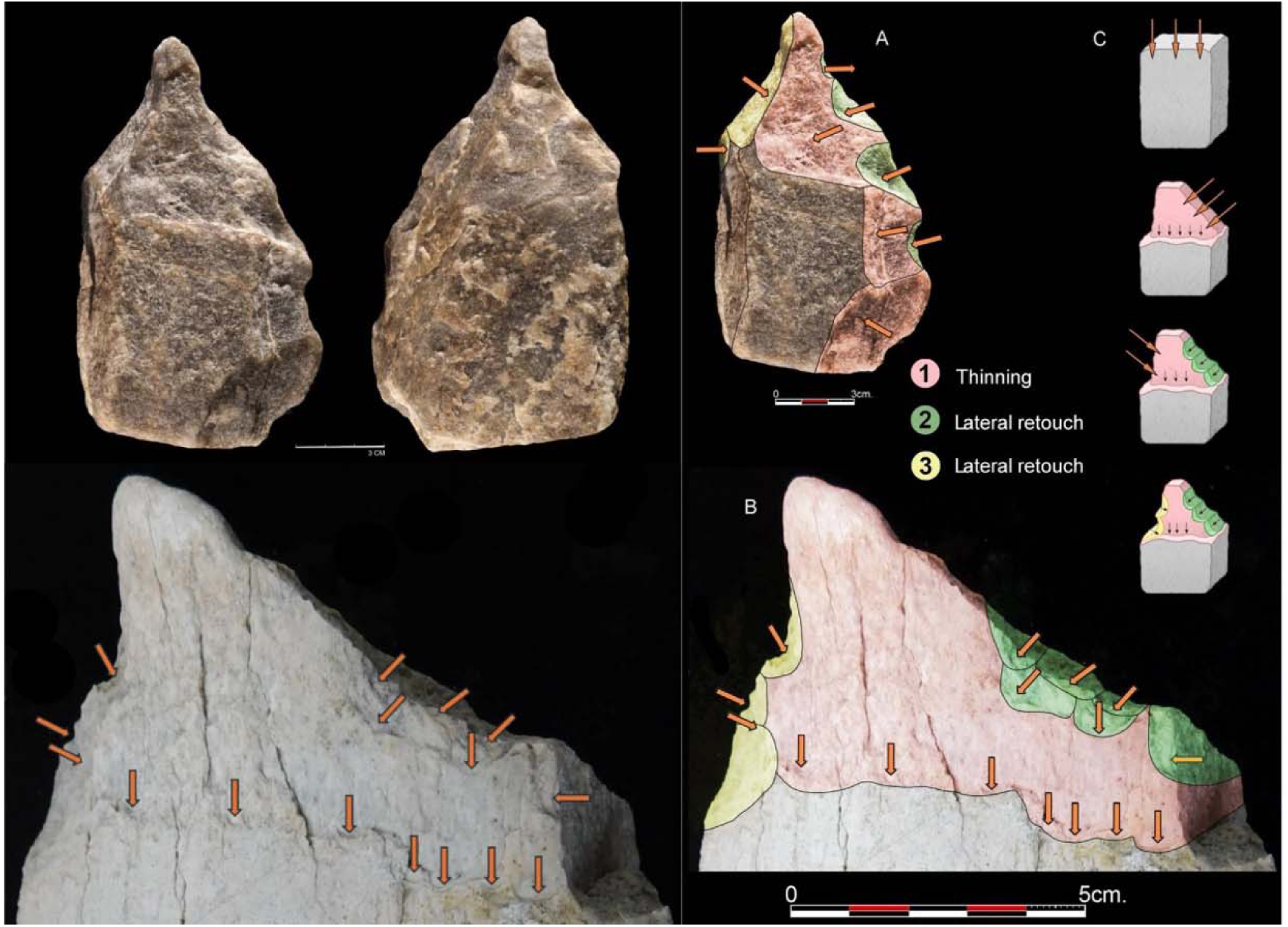
Left: Intentional shaping of points in a quartzite LCT from FLK West, and on a proboscidean femur shaft. Arrows indicate conchoidal scars probably caused by use (given their location away from the edge) and their reflection and stepped morphology by the pointed area. Notice the polishing at the point, which contrasts with the unpolished state of the remainder of the artifact. More complete cortical and medullary views of this artifact can be seen in Figures S6-S8. Right: Comparison of the configuration strategies identified on the quartz LCT and the LAS proboscidean femur shaft. A. LCT on a quartz slab from FLK West, Level 6; B. The proboscidean femoral shaft from LAS in the vicinity of EAK; C. Diacritical diagram showing point conceptualization sequencing identified in the previous specimens: 1. First thinning work (oblique to the distal end in the quartz LCT and vertical in the bone shaft); 2. Right-sided retouch; 3. Left-sided retouch. Series 2 and 3 determine the final shaping of the distal tip.

These results of widespread conspicuous evidence of hammerstone-breaking behaviors during Bed II partially confirm the recent report on bone artefacts discovered in Bed II by another research team (de la Torre et al., 2025). Excavations at T69 Complex area unearthed a wealth of megafaunal (proboscidean and hippopotamus) remains displaying hammerstone green breakage and hominin modifications. These remains seem to be spatially connected with a highly productive paleolandscape, since at its northern end it connects with BBS, a recently discovered site with megafaunal (Hippopotamus) remains. Green breakage at BBS also reflects hominin intervention in the palustrine environment congregating so many megafaunal carcasses. This unit would be equivalent to our second unit in Bed II.

We are not confident that the artefacts reported by dela Torre et al. at the T69 Complex are indeed tools. They are similar to the green-broken specimens that we have reported here. They are characterized by repeated number of flake scars along the edges; however, this is not a diagnostic feature, since multiple scars along edges also indirectly result from the repeated battering until the final breakage of the bone during butchery (Haynes et al., 2021, 2020; work in progress). In large bovids, it can also be derived from the gnawing of durophagous carnivores (Villa and Bartram, 1996) (Fig. S32). Carnivores capable of breaking elephant bones and producing the similar break shapes go back to at least the early Middle Miocene (Fig S33), although no carnivore today can effectively break the dense long bone mid-shafts of adult elephant bones. Another argument used by the authors is that the purported tools are longer and bear more flake scars compared to other faunal fragments (approximate average 13 versus 4). Again, this is documented among durophagous carnivores (Villa and Bartram, 1996; see examples in the Supplementary Information). A higher scar number in the T69 artifact collection could result from a correlation of different variables (i.e., longer dimension of the compared specimens, or different carcass sizes) and be, therefore, unrelated to anthropic intentionality. A third argument provided is that the items share consistent technological features, such as the production of morphologically similar, elongated, pointed, and notched bone tools, suggesting a patterned behavior. Again, this is equifinal with unintentional breaking of megafaunal remains during butchery (Haynes et al., 2021, 2020; Figs.S 6-8-10-11-16-19; experimental work in progress). There are several arguments that could be used to support this alternative:

1. The scars in the T69 artifacts exhibit an asymmetric distribution, characterized by stepped or reflected, incomplete fractures predominantly on the medullary surface, with minimal invasiveness. This contrasts with the expected conchoidal and complete scarring distributed equally across both cortical (i.e., periosteal) and medullary surfaces, as would be anticipated in cases of intentional flaking. Given that bone is more easily flaked than metamorphic rock, and that bifacial knapping of stone typically produces symmetrical, conchoidal scars on both surfaces, this would also be expected in the T69 bone specimens.
2. This bilateral symmetry principle is observed in Middle Pleistocene bone handaxes, where intentional modification is inferred based on the equal treatment of both sides and evidence of artificial shaping (Zutovski and Barkai, 2016). It is also documented on the similarly aged 1.4 Ma bone handaxe from Konso (Sano et al., 2020), and a purported handaxe from FC (Bed II, Olduvai) (Backwell and d’Errico, 2004), but it is absent from the reported items in T69, as well as those of the present study. Why, otherwise, would scars on the medullary surface be mostly limited to the wall of the breakage plane?
3. The repeated presence of incomplete, stepped scars on the medullary surfaces of the artifacts found at T69 aligns more closely with damage resulting from deliberate battering associated with breakage rather than subsequent edge retouching. The latter process would be expected to generate more controlled and complete conchoidal scars indicative of intentional shaping (Backwell and d’Errico, 2004; Haynes et al., 2021, 2020). This reflected scarring is also documented in Bos/Bison bones broken by hyenas, caused by redundancy in gnawing and the resulting overlapping micro-cracking leading to final collapse(Villa and Bartram, 1996) (Fig. S31).
4. Scar frequency is not a taphonomically valid criterion to attribute agency, since elephant bones can be broken in multiple ways, thus impacting experimentally-derived scar patterns. Sequential battering of initial crack lines with handheld hammerstones can generate overlapping notches and scars (work in progress). This is not necessarily reproduced when other experimental studies use alternative methods (such as throwing or bashing against immobile anvils) (Backwell and d’Errico, 2004; Haynes et al., 2021).
5. In several of the artifacts from T69, most of the overlapping scarring occurs in association to notches (and, in some cases, associated percussion marks) derived from dynamic loading. This reinforces that they may have been resulted from repeated battering during bone breaking.
6. The T69 Complex bone artifacts and those reported here are large-sized. If their potential function was for heavy-duty activities, those should have left any visible traces (at the macro and microscales) on the scarred edges. These are absent. A faunal assemblage that has good preservation of cut and percussion marks should certainly have good preservation conditions for microwear use and macro-damage scars.
7. If applying the same criteria defining bone tools at T69 Complex to Bos/Bison modified-bones, hyenas would also create similar artifacts (Fig. S32).

A final argument used by the authors to justify the intentional artifactual nature of their bone implements is that the bone tools were found *in situ* within a single stratigraphic horizon securely dated to 1.5 million years ago, indicating systematic production rather than episodic use. This is taphonomically unjustified. In a productive paleosurface where megafauna is abundant and hominin intervention is intense, the presence of this type of highly-scarred green-broken bones can be fairly common, since they result from butchery and carnivore post-depositional ravaging on non-proboscidean megafaunal bones (Domínguez-Rodrigo et al., 2014a, 2014b, 2009b; Villa and Bartram, 1996). Here, we have shown that most green-broken elephant bones cluster in three main areas of the Olduvai gorge.

Leakey collected more than a hundred purported bone tools from the same locations of Bed II where we collected our hammerstone-broken bones (Leakey, 1971) (Fig S33). All of them derive from Bed II (Backwell and d’Errico, 2004). A recent taphonomic review cautiously identified six potential bone tools in that collection (Pante et al., 2020). Some of these authors, using the much smaller T69 Complex artifact sample, now argue about systematic bone tool production. Bone artifacts (i.e., hominin-made items) that are used as tools should display evidence thereof through the macroscopic scarring, the polishing of the actively used surface, and the presence of microabrasive modifications on it (Backwell and d’Errico, 2008, 2001; d’Errico and Backwell, 2009; LeMoine, 1994). None of that has been documented in the T69 assemblage. We do not deny that the items reported by de la Torre et al. (2025) have the potential of having been used as tools, but proof is currently missing. The same applies to our reported sample (Fig 6; S4-S25). Only one artifact shows modifications that are potentially derived from its use as a tool. If confirmed, it is interesting to note the convergence in the point shape targeted in such an implement and those intentionally shaped in the earliest Acheulian implements documented at Olduvai Gorge (Figs 6, S6-S8). If such a bone implement was a tool, it would be the oldest bone tool documented to date (>1.7 Ma).

## Conclusions

Here, we have reported a significant change in hominin foraging behaviors during Bed I and Bed II times, roughly coinciding with the replacement of Oldowan industries by Acheulian tool kits -although during Bed II, both industries co-existed for a substantial amount of time (Domínguez-Rodrigo et al., 2023; Uribelarrea et al., 2019, 2017). This correlation does not necessarily imply causation, given that most tools documented around some of these butchery sites (e.g. EAK) are typologically Oldowan (mostly flakes and fragments). The traditional Acheulian large cutting tools do not seem to have played any role in megafaunal butchery, since they are most commonly undocumented or under-represented at early butchery sites, and where they are predominant or very common (e.g., TK), they are not functionally associated with faunal exploitation (see the examples of typological changes at TK, when *Elephas* and *Sivatherium* carcasses are exploited) (Panera et al., 2019). In contrast, the ocurrence of these megafaunal exploitation areas in three Bed II paleolandscape sections containing extremely large sites (e.g., FLK West in the LAS strata; SHK and BK in middle and upper Bed II respectively in the secondary gorge) attests to larger group sizes, which may be in connection with the intensity of exploitation of megafaunal resources. This was argued to be the main trigger in the thorough exploitation of a complete *Sivatherium* carcass at BK (Domínguez-Rodrigo et al., 2014), another one at TK (Panera et al.,. 2019), and several *Pelorovis* individuals at BK (Organista et al., 2016). The documentation of *H. erectus* around this time, with a much bigger anatomy than previous and sympatric hominins, suggests that a meat-based diet was behind these changes (Domínguez-Rodrigo et al., 2021). Having repeated access to high-yielding megafaunal taxa would have enabled access to a large supply of food for a large group for a substantial amount of time. The same as modern Hadza can fend off predators like lions from large kills, those early hominins might have sustained a similar eco-trophic position, especially given their bigger size. An additional important resource, fat, might have also pushed hominins to target proboscideans, since their long bones contain large amounts of fat, rich in nutritional EPA. This resource is currently under no competition, since modern hyenas cannot break open elephant long bone shafts from adult individuals.

The evidence presented here, together with that documented by de la Torre et al. (2025), represents the most geographically extensive documentation of repeated access to proboscidean and other megafaunal remains at a single fossil locality. The transition from Oldowan sites, where lithic and archaeofaunal assemblages are typically concentrated within 30–40 m² clusters, to Acheulean sites that span hundreds or even over 1000 m² (as in BK), with distinct internal spatial organization and redundancy in space use across multiple archaeological layers spanning meters of stratigraphic sequence (Domínguez-Rodrigo et al., 2014a, 2009b; Organista et al., 2017), reflects significant behavioral and technological shifts. This pattern likely signifies critical innovations in human evolution, coinciding with major anatomical and physiological transformations in early hominins (Dembitzer et al., 2022; Domínguez-Rodrigo et al., 2021, 2012). The causal relationships of these events that appear correlated in the archaeological record remain to be established.

## Methods

### Excavation of EAK

The systematic excavation of the stratigraphic layers involved a small crew. Materials exceeding 2 cm in size were meticulously plotted using laser theodolites (TOPCON total stations). Each trench was subjected to stereo-photography upon exposure, enabling the creation of photogrammetric 3D reconstructions of the paleosurface and deposited materials prior to recovery. The orientation and inclination of the artifacts were recorded using a compass and an inclinometer, respectively. Sediments removed from the trenches were sieved using 5 mm and 3 mm meshes.

The photogrammetric reconstructions of each trench were subsequently georeferenced and the resulting orthophotos were imported into AutoCAD, facilitating the detailed mapping of archaeological materials. The decision to conduct photogrammetry by individual trenches was made to minimize the exposure time of the excavated materials. Prolonged exposure could accelerate desiccation, potentially causing the most fragile specimens to become brittle and fragmentary. To mitigate this risk, excavation was limited to small, manageable areas, with small trenches proving ideal for efficient excavation due to the limited overburden. Each trench was excavated until the entire archaeological horizon was exposed, followed by comprehensive photogrammetric documentation prior to 3D plotting. Throughout the excavation, professional restoration experts provided necessary treatment for the bones. This was crucial for the recovery and identification of many bone fragments.

### Restoration of the EAK elephant bones

The main interventions were carried out in two locations: in situ, and in the field laboratory at the Emiliano Aguirre Research Station. For the consolidation and extraction of remains at the site, interventions were performed using Paraloid B-72 diluted in acetone at different concentrations, applied exclusively to fossils exhibiting powdering. This methodology strengthened the bones prior to extraction, ensuring better handling and reducing the risk of fractures. The use of Paraloid B-72 is a standard practice in the conservation of osseous remains due to its stability and compatibility with the material, though its limited penetration in some bone types has been noted(Daura et al., 2017). During this process, bandaging with sterile gauze impregnated with Paraloid B-72 was selectively applied to necessary areas to further protect fragile sections.

Additionally, superficial bandaging was conducted on larger fossils that visibly presented potential fractures and retained their cortical surface, aiming to prevent deterioration during the extraction process.

The greatest challenge was extracting long bones due to their size, weight, and fragility. These bones exhibited a greater presence of cortical surface, although they were not completely preserved, as they were fragmented but still retained significant areas suitable for study. Another significant challenge was the extraction of tusk remains, as the ivory exhibited a high degree of degradation and fragmentation. To recover these, multiple applications of consolidant were performed, allowing certain tusk blocks to be extracted in a controlled manner.

Larger fossils were extracted in block to minimize the risk of structural damage. The extraction process involved:

1. Initial consolidation by applying Paraloid B-72 dropwise to the surface, along with bandaging with sterile gauze in necessary areas to provide additional protection.
2. Surface protection with a sacrificial layer, avoiding direct contact with other reinforcing materials.
3. Creation of a counter-mold using plaster bandages, reinforced with interwoven steel rods to provide rigidity and stability.
4. Insertion of metal spatulas beneath the remains to lift and safely transport them.

In the field laboratory, cleaning of the bone remains was carried out, along with adhesion of fragments and their consolidation when necessary. The primary priority was preparing the remains for study by the rest of the team, opting for a purely conservative intervention, prioritizing detailed cleaning of some fragments for identification. Additionally, reassembly of larger fragments was performed, using Paraloid B-72 diluted in acetone at a higher concentration to ensure effective adhesion. For bone remains, mechanical and chemical cleaning procedures were applied depending on the nature of the adhered deposits.

This restoration procedure enabled the preservation of many bone fragments. Cleaning them properly will require much more time. In their current status, several specimens still have some adhering sedimentary matrix. This has prevented us from thoroughly examining all bone surfaces looking for BSM or conducting comprehensive identification. Some elements and fragments remain unidentified given also the lack of a proper comparative proboscidean collection. Despite this, most of the assemblage has been successfully studied.

### Taphonomic analysis of EAK

Skeletal part profiles were determined using the Number of Identifiable Specimens (NISP) and the Minimum Number of Elements (MNE) (Table S1). Given that with the exception of a few faunal specimens, the majority of the assemblage belongs to a single elephant individual, we did not use hierarchical units like MNI or MAU. To determine the MNE, ribs have been divided according to Rodríguez-Hidalgo’s (2017) system: portion 1 belongs to the epiphysis which includes the head, neck and costal tubercle; portion 2 belongs to the coastal angle; portion 3 belongs to the proximal body; portion 4 is composed by the distal body and portion 5 belongs to the sternal zone. The high frequency of fragmentation documented in the EAK’s ribs and the lack of landmarks made us determine the rib’s MNE through portions 1 and 2, in addition to a side-by-side comparison of rib shaft fragment widths (portions 3 and 4).

In the present study, presence of abrasion and polishing due to water dynamics (i.e., resedimentation, transport, lag) was taken into account. The first issue we analysed regarding the spatial distribution of archaeological remains was the orientation and slope of the items. First, we created stereograms and rose diagrams for the sample of elephant bone remains and lithic artifacts, as well as for both sets combined. These graphs were generated using OpenStereo software. Next, we conducted Rayleigh, Watson, and Kuiper tests to determine whether the isotropy/anisotropy tendence could be rejected (Fisher 1995).

An evaluation of the cortical surfaces was made, followed by an analysis of bone surface modifications; namely, cut marks, tooth marks, percussion marks and natural marks (i.e., biochemical and abrasion marks). Marks were identified by using hand lenses under strong direct light (60W) following the methodological and mark diagnostic criteria specified in Blumenschine (1995) and Blumenschine & Selvaggio (1989) for tooth and percussion marks, Dominguez-Rodrigo et al. (2009a, 2010b) for cut marks, and Domínguez-Rodrigo and Barba (2006) for biochemical marks. The analysis of bone breakage at EAK is based on the identification of green and dry breakage planes following Villa and Mahieu’s (1991) criteria.

### Lithic analysis of EAK

A total of 80 lithic specimens have been recovered from EAK. The technological analysis of these materials follows established classification criteria and methodological frameworks(Diez-Martín et al., 2009), categorizing artifacts into the following groups:

(a) Unmodified Specimens – Naturally occurring cobblestones lacking evidence of anthropogenic modification.
(b) Percussion Tools – Includes hammerstones or cobbles exhibiting diagnostic battering, pitting, and/or impact scars consistent with percussive activities.
(c) Cores – Handheld cores are classified based on key technological attributes: (i) the number of reduction surfaces (unifacial, bifacial, multifacial), (ii) the number of striking platforms (unipolar, bipolar, multipolar), and (iii) the spatial organization of knapping sequences (linear, opposed, orthogonal, centripetal). Bipolar cores are identified based on diagnostic features associated with anvil-supported percussion.
(d) Detached Products – This category encompasses both complete and fragmented flakes, derived from either bipolar or freehand reduction. Technological attributes recorded for detached products include core reduction stage (assessed through the presence and extent of cortical coverage on dorsal surfaces and striking platforms), platform morphology, and dorsal scar patterning.
(e) Knapping By-Products – Includes all non-diagnostic lithic debris resulting from core reduction and flake detachment. This category incorporates non-diagnostic flake fragments with evidence of direct handheld percussion, shatter, blocky/angular undetermined detached fragments, core-like fragments, and debris ≤20 mm in size.

### The spatial statistical analyses of EAK

The spatial point patterns (SPP) of faunal and lithic remains have been analyzed using the ‘spatstat’ library(Baddeley et al., 2015) in R (www.r-project.org). This methodological approach is of great interest for paleontological and archaeological contexts, as it allows for the analysis of aspects such as the degree of randomness of the SPP, potential relationships between different types of materials, or enabling predictions about unexcavated areas. We conducted the analysis in three different ways after selecting the spatial window, i.e. the analysed excavated area (52.56 m^2^).

First, we analysed the SPPs without marks (i.e., raw SPPs, without considering any specific spatial variable) for the elephant remains and lithic remains independently. In these initial cases, we created intensity maps, hot spot maps, and hot spot maps with a 99% Confidence Interval (C.I.). Next, we applied a chi-squared test with four different grid configurations (5×5, 8×8, 10×10, 12×12)(Moclán et al., 2023) to determine whether the SPPs are homogeneous or inhomogeneous. These analyses showed that, in all cases, the SPPs can be considered inhomogeneous (p-value <0.05). Therefore, we proceeded to use tests to analyse the type of SPP. We employed the Pair Correlation, *K*, *L*, *F*, *G*, and *J* functions in their inhomogeneous versions, as well as the *Kscaled* and *Lscaled* functions to analyse the typology of the SPPs (i.e., random, cluster, or regular). Both versions of the functions were used because the sample size does not allow us to analyse the type of inhomogeneity through a studentized permutation test. Similarly, we used the DCLF (Diggle-Cressie-Loosmore-Ford) and MAD (Maximum Absolute Deviation) tests with the *K*, *L*, *F*, *G*, *J*, *Kscaled*, and *Lscaled* functions to check the typology of the point patterns (i.e., random vs. non-random). Finally, we used the ‘nnclean’ function(Baddeley et al., 2015) to detect possible clustering of materials through nearest neighbour clutter removal. Lastly, we performed spatial modelling using linear, quadratic, cubic, and 4^th^-degree polynomial regressions to delve deeper into the characteristics of the SPPs. The models were evaluated using the Akaike Information Criterion (AIC) index, and subsequently, we compared their validity through ANOVA. Finally, we selected the best-performing model (i.e., the one with the lowest AIC values and that does not show issues of collinearity or suffer from the small sample size) and generated four maps: 1) estimated intensity of the point process based on the fitted model, 2) simulation of a new point pattern using the fitted model with the Metropolis-Hastings algorithm, 3) a difference map subtracting the intensity of the simulated new point pattern from the “real” intensity of the spatial point pattern (Domínguez-Rodrigo et al., 2024), and 4) an extrapolated estimation of potential points within a larger spatial window (308.23 m^2^) than the original one.

The second analysis consists of examining the degree of statistical correlation between two marks, i.e., spatial variables, in this case, fauna vs. lithic tools. To do this, we first calculated the relative risk to assess the likelihood of each type of material appearing in the analysed window. Then, we applied a chi-square test to verify once again, using the same grid configuration, whether the samples are homogeneous or inhomogeneous. The chi-square test again indicates that the spatial distribution is inhomogeneous. Therefore, we used the *K*·, *L*·, *K*cross, and *L*cross inhomogeneous functions to analyse whether there is spatial correlation between the two material types and, if so, what kind it is.

Finally, given the size disparity between the elephant remains and the lithics, we analysed the relationship between both types of materials by considering the location of the elephant remains as a covariate. First, a distance map was generated to the centroids of the bone remains. ANOVA was then conducted to assess the fitted effect of the covariate in the SPPs. This was followed by the application of the ρ function (‘rhohat’ function) to investigate the influence of the distance to the bones. This test is similar to the previous one but provides a more detailed view of the potential variations in the intensity of remains with respect to distance from the bones. Finally, a Z_1_ and Z_2_ Berman–Lawson–Waller test was performed.

### Megafaunal exploitation along the paleolandscape sequence of Olduvai Gorge

An intensive survey seeking stratigraphically-associated megafaunal bones was carried out in the months of June 2023 and 2024. We targeted the Bed I and Bed II exposures of the main Gorge (From FLK NNN to the KK fault), and the side gorge, from the junction until BK. Linearly, this comprises over 6 km of outcrops. We targeted this area because it contains the highest density of fossils of all the gorge. We focused on proboscidean bones and used hippopotamus bones, some of the most abundant in the megafaunal fossils, as a spatial control. Bones were counted as separate carcasses when enough distance existed in between findings (i.e., localities). Stratigraphic association was carried out by direct observation of the geological context and with the presence of a Quaternary geologist during the whole survey. When fossils found were ambiguously associated with specific strata, these were excluded from the present analysis.

For the sake of analysis, and given that we were not focusing on time-resolution landscapes, we used four time intervals: Bed I, Bed II lower (from Tuff IF to LAS), Bed II intermediate (from Tuff IIB to Tuff IIC, and Bed II upper (from Tuff IID to the interface with Bed III). Only in one case, Bed II lower, were most of the remains circumscribed to a single stratigraphic unit (Lower Augitic Sands [LAS], which mark the beginning of Haýs middle Bed II). The goals of this survey were: a) collect a spatial sample of proboscidean and megafaunal bones enabling us to understand if carcasses on the Olduvai paleolandscapes were randomly deposited or associated to specific habitats; b) given the water dependence of elephants, we wanted to understand if they showed the same type of ecological association to water-dependent megafauna, like hippopotamus, as their modern counterparts (using taphocoenoses as indicators of past megafaunal biocoenosis); c) approach potential anthropogenic agency in proboscidean exploitation by comparing carcass dispersal and accumulation and productivity of anthropogenic landscapes (i.e., concentrations and dispersals of lithic artifacts); d) seek anthropogenic traces of proboscidean exploitation in the form of cut and percussion marks and green bone breakage; e) associate this latter point with the hominin use of space, by determining if these taphonomic evidences occurred randomly or clustered and spatially associated with stone tools.

## Acknowledgements

We thank the Spanish Ministry of Science and Innovation for funding this research (PID2023-146260NB-C2), and the Spanish Ministry of Culture for their funding through the program of Archaeology Abroad. We thank the Commission for Science and Technology (COSTECH), the Ngorongoro Conservation Area Authorities (NCAA), the Division of Antiquities, and the Tanzanian Ministry of Natural Resources and Tourism for their permission to conduct research in Tanzania. AM is funded by a postdoctoral contract grant from the Fyssen Foundation. We are grateful to the Tanzanian co-workers at Olduvai Gorge. We are deeply indebted to the constructive comments made by G. Haynes, R. Barkai and one anonymous reviewer to an earlier version of this manuscript. We also thank a second anonymous reviewer, despite our disagreement with some of his/her comments. We also appreciate the professional editorship of Yonatan Sahle and Detlef Weigel.

## Footnotes

1 In this study, *megafauna* refers to animals with a body mass exceeding 800 kg

## References

Adrien Hannus L. 2018. Clovis Mammoth Butchery: The Lange/Ferguson Site and Associated Bone Tool Technology. Texas A&M University Press.

Agam A, Barkai R. 2018. Elephant and Mammoth Hunting during the Paleolithic: A Review of the Relevant Archaeological, Ethnographic and Ethno-Historical Records. Quaternary 1:3.

Agam A, Barkai R. 2016. Not the brain alone: The nutritional potential of elephant heads in Paleolithic sites. Quat Int 406:218–226.

Altamura F, Gaudzinski-Windheuser S, Melis RT, Mussi M. 2020. Reassessing Hominin Skills at an Early Middle Pleistocene Hippo Butchery Site: Gombore II-2 (Melka Kunture, Upper Awash valley, Ethiopia). Journal of Paleolithic Archaeology 3:1–32.

A matter of fat: Hunting preferences affected Pleistocene megafaunal extinctions and human evolution. n.d.

Arráiz H, Barboni D, Uribelarrea D, Mabulla A, Baquedano E, Domínguez-Rodrigo M. 2017. Paleovegetation changes accompanying the evolution of a riverine system at the BK paleoanthropological site (Upper Bed II, Olduvai Gorge, Tanzania). Palaeogeogr Palaeoclimatol Palaeoecol 488:84–92.

Ashley G. 2003. Tracks, Trails and Trampling by Large Vertebrates in a Rift Valley Paleo-Wetland, Lowermost Bed II, Olduvai Gorge, Tanzania. Ichnos 9:23–32.

Ashley GM, Tactikos JC, Owen RB. 2009. Hominin use of springs and wetlands: Paleoclimate and archaeological records from Olduvai Gorge (∼1.79–1.74 Ma). Palaeogeogr Palaeoclimatol Palaeoecol 272:1–16.

Backwell L, d’Errico F. 2008. Early hominid bone tools from Drimolen, South Africa. J Archaeol Sci 35:2880–2894.

Backwell L, d’Errico F. 2004. The first use of bone tools: a reappraisal of the evidence from Olduvai Gorge, Tanzania. Palaeontologia africana 40:95–158.

Backwell LR, d’Errico F. 2001. Evidence of termite foraging by Swartkrans early hominids. Proc Natl Acad Sci U S A 98:1358–1363.

Baddeley A, Rubak E, Turner R. 2015. Spatial Point Patterns, Chapman & Hall/CRC Interdisciplinary Statistics. Oakville, MO: Apple Academic Press.

Barkai R. 2019. When Elephants Roamed Asia: The Significance of Proboscideans in Diet, Culture and Cosmology in Paleolithic Asia In: Kowner R, Bar-Oz G, Biran M, Shahar M, Shelach-Lavi G, editors. Animals and Human Society in Asia: Historical, Cultural and Ethical Perspectives. Cham: Springer International Publishing. pp. 33–62.

Ben-Dor, M., Barkai, R. 2024. A matter of fact: hunting preferences affected Pleistocene megafaunal extinctions and human evolution. Quaternary Science Reviews, 331. 10.1016/j.quascirev.2024.108660.

Ben-Dor M, Barkai R. 2020. The importance of large prey animals during the Pleistocene and the implications of their extinction on the use of dietary ethnographic analogies. Journal of Anthropological Archaeology 59:101192.

Ben-Dor M, Gopher A, Hershkovitz I, Barkai R. 2011. Man the fat hunter: the demise of Homo erectus and the emergence of a new hominin lineage in the Middle Pleistocene (ca. 400 kyr) Levant. PLoS One 6:e28689.

Berthelet A, Chavaillon J. 2001. The early Palaeolithic butchery site of Barogali (Republic of Djibouti)The World of Elephants. International Congress Proceedings, Consiglio Nazionale Delle Ricerche, Rome. pp. 176–179.

Boschian G, Saccà D. 2015. In the elephant, everything is good: Carcass use and re-use at Castel di Guido (Italy). Quat Int 361:288–296.

Blumenschine, R.J., 1995. Percussion marks, tooth marks, and experimental determinations of the timing of hominid and carnivore access to long bones at FLK Zinjanthropus, Olduvai Gorge, Tanzania. Journal of human Evolution, 29(1), pp.21–51.

Blumenschine, R.J. and Selvaggio, M.M., 1988. Percussion marks on bone surfaces as a new diagnostic of hominid behaviour. Nature, 333(6175), pp.763–765.

Bunge, M. 1981. Analogy between systems. International Journal og General Systems 7: 221–223.

Bunn HT. 1982. Meat-eating and Human Evolution: Studies on the Diet and Subsistence Patterns of Plio-Pleistocene Hominids in East Africa. University of California, Berkeley.

Chavaillon, J., Berthelet, A., 2004. The archaeological sites of Melka Kunture. In: Chavaillon, J., Piperno, M. (Eds.), Studies on the Early Paleolithic Site of Melka Kunture, Ethiopia. Istituto Italiano di Preistoria e Protostoria, Roma, pp. 25–80.

Chavaillon J, Boisaubert J-L, Faure M, Guerin C, Ma J-L, Nickel B. 1987. Le site de dépeçage pléistocène à Elephas recki de Barogali (République de Djibouti): nouveaux résultats et datation. C R Acad Sci II 305:1259–1266.

Crader DC. 1983. Recent single-carcass bone scatters and the problem of “butchery” sites in the archaeological record. Animal and Archaeology 1 Hunters and their Prey, BAR International Series 163:107–141.

Daura J, Sanz M, Arsuaga JL, Hoffmann DL, Quam RM, Ortega MC, Santos E, Gómez S, Rubio A, Villaescusa L, Souto P, Mauricio J, Rodrigues F, Ferreira A, Godinho P, Trinkaus E, Zilhão J. 2017. New Middle Pleistocene hominin cranium from Gruta da Aroeira (Portugal). Proc Natl Acad Sci U S A 114:3397–3402.

Deino AL. 2012. (40)Ar/(39)Ar dating of Bed I, Olduvai Gorge, Tanzania, and the chronology of early Pleistocene climate change. J Hum Evol 63:251–273.

Delagnes A, Lenoble A, Harmand S, Brugal J-P, Prat S, Tiercelin J-J, Roche H. 2006. Interpreting pachyderm single carcass sites in the African Lower and Early Middle Pleistocene record: A multidisciplinary approach to the site of Nadung’a 4 (Kenya). J Anthropol Archaeol 25:448–465.

de la Torre I, Doyon L, Benito-Calvo A, Mora R, Mwakyoma I, Njau JK, Peters RF, Theodoropoulou A, d’Errico F. 2025. Systematic bone tool production at 1.5 million years ago. Nature. doi:10.1038/s41586-025-08652-5

Dembitzer J, Barkai R, Ben-Dor M, Meiri S. 2022. Levantine overkill: 1.5 million years of hunting down the body size distribution. Quat Sci Rev 276:107316.

Deocampo DM, Berry PA, Beverly EJ, Ashley GM, Jarrett RE. 2017. Whole-rock geochemistry tracks precessional control of Pleistocene lake salinity at Olduvai Gorge, Tanzania: A record of authigenic clays. Geology 45:683–686.

Deocampo DM, Blumenschine RJ, Ashley GM. 2002. Wetland Diagenesis and Traces of Early Hominids, Olduvai Gorge, Tanzania. Quat Res 57:271–281.

d’Errico F, Backwell L. 2009. Assessing the function of early hominin bone tools. J Archaeol Sci 36:1764–1773.

Diez-Martín F, Fraile C, Uribelarrea D, Sánchez-Yustos P, Domínguez-Rodrigo M, Duque J, Díaz I, De Francisco S, Yravedra J, Mabulla A, Baquedano E. 2017.SHKExtension: a new archaeological window in theSHKfluvial landscape of Middle BedII(Olduvai Gorge, Tanzania). Boreas 46:831–859.

Diez-Martín F, Fraile-Márquez C, Duque-Martínez J, Sánchez-Yustos P, de Francisco S, Baquedano E, Mabulla A, Domínguez-Rodrigo M. 2024. On time scales and “synchronic” variability in the archaeology of human origins: short-term technological variations at SHK (Olduvai Gorge, Tanzania). Archaeol Anthropol Sci 16. doi:10.1007/s12520-024-02092-4

Diez-Martín F, Sánchez P, Domínguez-Rodrigo M, Mabulla A, Barba R. 2009. Were Olduvai Hominins making butchering tools or battering tools? Analysis of a recently excavated lithic assemblage from BK (Bed II, Olduvai Gorge, Tanzania). Journal of Anthropological Archaeology 28:274–289.

Diez-Martín F, Sánchez Yustos P, Uribelarrea D, Baquedano E, Mark DF, Mabulla A, Fraile C, Duque J, Díaz I, Pérez-González A, Yravedra J, Egeland CP, Organista E, Domínguez-Rodrigo M. 2015. The origin of the Acheulean: The 1.7 million-year-old site of FLK West, Olduvai Gorge (Tanzania). Sci Rep 5:17839.

Diez-Martín F, Sánchez-Yustos P, Uribelarrea D, Domínguez-Rodrigo M, Fraile-Márquez C, Obregón R-A, Díaz-Muñoz I, Mabulla A, Baquedano E, Pérez-González A, Bunn HT. 2014. New archaeological and geological research at SHK main site (Bed II, Olduvai Gorge, Tanzania). Quat Int 322-323:107–128.

Domıinguez-Rodrigo, M., 1997. Meat-eating by early hominids at the FLK 22Zinjanthropussite, Olduvai Gorge (Tanzania): an experimental approach using cut-mark data. Journal of human Evolution, 33(6), pp.669–690.

Domínguez-Rodrigo, M & Barba, R. (2006). New estimates of tooth marks and percussion marks from FLK Zinj, Olduvai Gorge (Tanzania): the carnivore-hominid-carnivore hypothesis falsified. Journal of Human Evolution 50:170.194.

Domínguez-Rodrigo M, Baquedano E, Organista E, Cobo-Sánchez L, Mabulla A, Maskara V, Gidna A, Pizarro-Monzo M, Aramendi J, Galán AB, Cifuentes-Alcobendas G, Vegara-Riquelme M, Jiménez-García B, Abellán N, Barba R, Uribelarrea D, Martín-Perea D, Diez-Martin F, Maíllo-Fernández JM, Rodríguez-Hidalgo A, Courtenay L, Mora R, Maté-González MA, González-Aguilera D. 2021. Early Pleistocene faunivorous hominins were not kleptoparasitic, and this impacted the evolution of human anatomy and socio-ecology. Sci Rep 11:16135.

Domínguez-Rodrigo M, Barba R, Egeland CP. 2007.Deconstructing Olduvai: A Taphonomic Study of the Bed I Sites. Springer Science & Business Media.

Domínguez-Rodrigo M, Bunn HT, Mabulla AZP, Baquedano E, Uribelarrea D, Pérez-González A, Gidna A, Yravedra J, Díez-Martín F, Egeland CP, Others. 2014a. On meat eating and human evolution: A taphonomic analysis of BK4b (Upper Bed II, Olduvai Gorge, Tanzania), and its bearing on hominin megafaunal consumption. Quat Int 322:129–152.

Domínguez-Rodrigo M, Cobo-Sánchez L, Aramendi J, Gidna A. 2019. The meta-group social network of early humans: A temporal–spatial assessment of group size at FLK Zinj (Olduvai Gorge, Tanzania). J Hum Evol 127:54–66.

Domínguez-Rodrigo M, Cobo-Sánchez L, Baquedano E, Mabulla A, Gidna A, Diez-Martin F. 2024. Reconstructing Olduvai: The Behavior of Early Humans at David’s Site. Elsevier Science.

Domínguez-Rodrigo M, de Juana S, Galán AB, Rodríguez M. 2009a. A new protocol to differentiate trampling marks from butchery cut marks. J Archaeol Sci 36:2643–2654.

Domínguez-Rodrigo M, Diez-Martín F, Yravedra J, Barba R, Mabulla A, Baquedano E, Uribelarrea D, Sánchez P, Eren MI. 2014b. Study of the SHK Main Site faunal assemblage, Olduvai Gorge, Tanzania: Implications for Bed II taphonomy, paleoecology, and hominin utilization of megafauna. Quat Int 322-323:153–166.

Domínguez-Rodrigo M, Mabulla A, Bunn HT, Barba R, Diez-Martín F, Egeland CP, Espílez E, Egeland A, Yravedra J, Sánchez P. 2009b. Unraveling hominin behavior at another anthropogenic site from Olduvai Gorge (Tanzania): new archaeological and taphonomic research at BK, Upper Bed II. J Hum Evol 57:260–283.

Domínguez-Rodrigo M, Mabulla AZP, Bunn HT, Diez-Martin F, Baquedano E, Barboni D, Barba R, Domínguez-Solera S, Sánchez P, Ashley GM, Yravedra J. 2010. Disentangling hominin and carnivore activities near a spring at FLK North (Olduvai Gorge, Tanzania). Quat Res 74:363–375.

Domínguez-Rodrigo M, Pickering TR, Bunn HT. 2010b. Configurational approach to identifying the earliest hominin butchers. Proceedings of the National Academy of Sciences 107:20929–20934.

Domínguez-Rodrigo M, Pickering TR, Diez-Martín F, Mabulla A, Musiba C, Trancho G, Baquedano E, Bunn HT, Barboni D, Santonja M, Uribelarrea D, Ashley GM, Martínez-Ávila M del S, Barba R, Gidna A, Yravedra J, Arriaza C. 2012. Earliest porotic hyperostosis on a 1.5-million-year-old hominin, olduvai gorge, Tanzania. PLoS One 7:e46414.

Domínguez-Rodrigo M, Uribelarrea D, Diez-Martín F, A, Mabulla, Gidna A, Cobo-Sánchez L, Martín-Perea DM, Organista E, Barba R, Baquedano E. 2023. Earliest Acheulian paleolandscape reveals a 1.7 million-year-old megasite at Olduvai Gorge (Tanzania). Quat Sci Rev 316:108262.

Dominguez-Rodrigo, M., Bunn, H.T., Yravedra, J. 2014. A critical re-evaluation of bone surface modification models for inferring fossil hominin and carnivore interactions through a multivariate approach: application to the FLK Zinj archaeofaunal assemblage (Olduvai Gorge, Tanzania). Quaternary International 322-323: 32–43.

Fisher NI (1995) Statistical Analysis of Circular Data. Cambridge University Press, Cambridge.

Fujioka T, Benito-Calvo A, Mora R, McHenry L, Njau JK, de la Torre I. 2022. Direct cosmogenic nuclide isochron burial dating of early Acheulian stone tools at the T69 Complex (FLK West, Olduvai Bed II, Tanzania). J Hum Evol 165:103155.

Garrett K. 2017. Multi-proxy reconstuction of a ∼1.3 Ma freshwater wetland, Olduvai Gorge, Tanzania. doi:10.7282/T3T43WJ0

Gaudzinski S, Turner E, Anzidei AP, Àlvarez-Fernández E, Arroyo-Cabrales J, Cinq-Mars J, Dobosi VT, Hannus A, Johnson E, Münzel SC, Scheer A, Villa P. 2005. The use of Proboscidean remains in every-day Palaeolithic life. Quat Int 126-128:179–194.

Gaudzinski-Windheuser S, Kindler L, MacDonald K, Roebroeks W. 2023a. Hunting and processing of straight-tusked elephants 125.000 years ago: Implications for Neanderthal behavior. Sci Adv 9:eadd8186.

Gaudzinski-Windheuser S, Kindler L, Roebroeks W. 2023b. Widespread evidence for elephant exploitation by Last Interglacial Neanderthals on the North European plain. Proc Natl Acad Sci U S A 120:e2309427120.

Giusti D. 2021. Investigating the spatio-temporal dimension of past human-elephant interactions: a spatial taphonomic approach.

Gomez MS, Lopez N, Gonzalez AP. 1978. Acheulean occupation sites in the Jarama valley (Madrid, Spain). Curr Anthropol 19:394–395.

Goren-Inbar N, Lister A, Werker E, Chech M. 1994. A Butchered elephant skull and associated artifacts from the Acheulian site of Gesher Benot Ya’Aqov, Israel. Paléorient 20:99–112.

Haynes, G. (1983). A guide for differentiating mammalian carnivore taxa responsible for gnaw damage to herbivore limb bones. Paleobiology, 9(2), 164–172.

Haynes G. 2022. Late Quaternary Proboscidean Sites in Africa and Eurasia with Possible or Probable Evidence for Hominin Involvement. Quaternary 5:18.

Haynes G. 1991. Mammoths, Mastodonts, and Elephants: Biology, Behavior and the Fossil Record. Cambridge University Press.

Haynes G. 1988. Longitudinal studies of african elephant death and bone deposits. J Archaeol Sci 15:131–157.

Haynes G, Hutson J. 2020. African elephant bones modified by carnivores: Implications for interpreting fossil proboscidean assemblages. Journal of Archaeological Science: Reports 34:102596.

Haynes G, Krasinski K. 2021. Butchering marks on bones of Loxodonta africana (African savanna elephant): Implications for interpreting marks on fossil proboscidean bones. Journal of Archaeological Science: Reports 37:102957.

Haynes G, Krasinski K, Wojtal P. 2021. A Study of Fractured Proboscidean Bones in Recent and Fossil Assemblages. Journal of Archaeological Method and Theory 28:956–1025.

Haynes G, Krasinski K, Wojtal P. 2020. Elephant bone breakage and surface marks made by trampling elephants: Implications for interpretations of marked and broken Mammuthus spp. bones. Journal of Archaeological Science: Reports 33:102491.

Hay R.1976. Geology of the Olduvai Gorge. Univ of California Press.

Isaac GL, Isaac B. 2023. Koobi Fora Research Project: Plio-Pleistocene archaeology. Clarendon.

Keefer DK. 1984. Landslides caused by earthquakes. GSA Bulletin 95:406–421.

Konidaris GE, Athanassiou A, Tourloukis V, Thompson N, Giusti D, Panagopoulou E, Harvati K. 2018. The skeleton of a straight-tusked elephant (Palaeoloxodon antiquus) and other large mammals from the Middle Pleistocene butchering locality Marathousa 1 (Megalopolis Basin, Greece): preliminary results. Quat Int 497:65–84.

Konidaris GE, Barkai R, Turlukēs B, Harvati K. 2021. Human-elephant Interactions: From Past to Present. Tübingen University Press.

Labrado EO. 2017. Estudio tafonómico de los niveles arqueológicos de BK (Bell Korongo), garganta de Olduvai, Tanzania.

Leakey MD. 1971. Olduvai Gorge: Volume 3, Excavations in Beds I and II, 1960-1963. Cambridge University Press.

LeMoine GM. 1994. Use wear on bone and antler tools from the Mackenzie Delta, Northwest Territories. Am Antiq 59:316–334.

Lemorini C, Santucci E, Caricola I, Nucara A, Nunziante-Cesaro S. 2023. Life around the elephant in space and time: An integrated approach to study the human-elephant interactions at the Late Lower Paleolithic site of La Polledrara di Cecanibbio (Rome, Italy). J Archaeol Method Theory 30:1233–1281.

Lev M ‘ayan, Barkai R. 2016. Elephants are people, people are elephants: Human–proboscideans similarities as a case for cross cultural animal humanization in recent and Paleolithic times. Quat Int 406:239–245.

Liutkus CM, Ashley GM. 2003. Facies Model of a Semiarid Freshwater Wetland, Olduvai Gorge, Tanzania. J Sediment Res 73:691–705.

Lyman, R.L. 2006. Vertebrate Taphonomy. Cambridge University Prress, Cambridge.

Marlowe F. 2010. The Hadza: Hunter-gatherers of Tanzania. University of California Press.

Moclán A, Cobo-Sánchez L, Domínguez-Rodrigo M, Méndez-Quintas E, Rubio-Jara S, Panera J, Pérez-González A, Santonja M. 2023. Spatial analysis of an Early Middle Palaeolithic kill/butchering site: the case of the Cuesta de la Bajada (Teruel, Spain). Archaeol Anthropol Sci 15. doi:10.1007/s12520-023-01792-7

Mussi M, Villa P. 2008. Single carcass of Mammuthus primigenius with lithic artifacts in the Upper Pleistocene of northern Italy. J Archaeol Sci 35:2606–2613.

O’Connell JF, Hawkes K, Blurton-Jones NG. 1992. Patterns in the distribution, site structure and assemblage composition of Hadza kill-butchering sites. J Archaeol Sci 19:319–345.

Organista E, Arriaza MC, Barba R, Gidna A, Ortega MC, Uribelarrea D, Mabulla A, Baquedano E, Domínguez-Rodrigo M. 2019. Taphonomic analysis of the level 3b fauna at BK, Olduvai Gorge. Quat Int 526:116–128.

Organista E, Domínguez-Rodrigo M, Egeland CP, Uribelarrea D, Mabulla A, Baquedano E. 2016. Did Homo erectus kill a Pelorovis herd at BK (Olduvai Gorge)? A taphonomic study of BK5. Archaeol Anthropol Sci 8:601–624.

Organista E, Domínguez-Rodrigo M, Yravedra J, Uribelarrea D, Arriaza MC, Ortega MC, Mabulla A, Gidna A, Baquedano E. 2017. Biotic and abiotic processes affecting the formation of BK Level 4c (Bed II, Olduvai Gorge) and their bearing on hominin behavior at the site. Palaeogeogr Palaeoclimatol Palaeoecol 488:59–75.

Panera J, Rubio-Jara S, Domínguez-Rodrigo M, Yravedra J, Méndez-Quintas E, Pérez-González A, Bello-Alonso P, Moclán A, Baquedano E, Santonja M. 2019. Assessing functionality during the early Acheulean in level TKSF at Thiongo Korongo site (Olduvai Gorge, Tanzania). Quat Int 526:77–98.

Pante M, Torre I de la, d’Errico F, Njau J, Blumenschine R. 2020. Bone tools from Beds II–IV, Olduvai Gorge, Tanzania, and implications for the origins and evolution of bone technology. J Hum Evol 148:102885.

Pizarro-Monzo, M., Prendergast, M., Gidna, A., Baquedano, E, Mora, R., González-Aguilera, D., Maté-Gonzalez, MA, Domínguez-Rodrigo, M. (2021). Do human butchery patterns exist? A study of the interaction of randomness and channelling in the distribution of cut marks on long bonesJ. R. Soc. Interface.18 2020095820200958.

Plummer TW, Oliver JS, Finestone EM, Ditchfield PW, Bishop LC, Blumenthal SA, Lemorini C, Caricola I, Bailey SE, Herries AIR, Parkinson JA, Whitfield E, Hertel F, Kinyanjui RN, Vincent TH, Li Y, Louys J, Frost SR, Braun DR, Reeves JS, Early EDG, Onyango B, Lamela-Lopez R, Forrest FL, He H, Lane TP, Frouin M, Nomade S, Wilson EP, Bartilol SK, Rotich NK, Potts R. 2023. Expanded geographic distribution and dietary strategies of the earliest Oldowan hominins and Paranthropus. Science 379:561–566.

Rabinovich, R., Ackermann, O., Aladjem, E., Barkai, R., Biton, R., Milevski, I., Solodenko, N., and Marder, O., 2012. Elephants at the middle Pleistocene Acheulian open-air site of Revadim Quarry, Israel. Quaternary International, 276, pp.183–197.

Reshef H, Barkai R. 2015. A taste of an elephant: The probable role of elephant meat in Paleolithic diet preferences. Quat Int 379:28–34.

Rocca R, Boschin F, Aureli D. 2021. Around an elephant carcass: Cimitero di Atella and Ficoncella in the behavioural variability during the Early Middle Pleistocene in italy.

Rowe TB, Stafford TW, Fisher DC, Enghild JJ, Quigg JM, Ketcham RA, Sagebiel JC, Hanna R, Colbert MW. 2022. Human Occupation of the North American Colorado Plateau C37,000 Years Ago. Frontiers in Ecology and Evolution 10. doi:10.3389/fevo.2022.903795

Saccà D. 2012.Taphonomy of Palaeloxodon antiquus at Castel di Guido (Rome, Italy): Proboscidean carcass exploitation in the Lower Palaeolithic. Quat Int 276-277:27–41.

Sahnouni M, Rosell J, van der Made J, Vergès JM, Ollé A, Kandi N, Harichane Z, Derradji A, Medig M. 2013. The first evidence of cut marks and usewear traces from the Plio-Pleistocene locality of El-Kherba (Ain Hanech), Algeria: implications for early hominin subsistence activities circa 1.8 Ma. J Hum Evol 64:137–150.

Sano K, Beyene Y, Katoh S, Koyabu D, Endo H, Sasaki T, Asfaw B, Suwa G. 2020. A 1.4-million-year-old bone handaxe from Konso, Ethiopia, shows advanced tool technology in the early Acheulean. Proc Natl Acad Sci U S A 117:18393–18400.

Santucci E, Marano F, Cerilli E, Fiore I, Lemorini C, Palombo MR, Anzidei AP, Bulgarelli GM. 2016. Palaeoloxodon exploitation at the Middle Pleistocene site of La Polledrara di Cecanibbio (Rome, Italy). Quat Int 406:169–182.

Solodenko N, Zupancich A, Cesaro SN, Marder O, Lemorini C, Barkai R. 2015. Fat residue and use-wear found on Acheulian biface and scraper associated with butchered elephant remains at the site of Revadim, Israel. PLoS One 10:e0118572.

Tappen M, Bukhsianidze M, Ferring R, Coil R, Lordkipanidze D. 2022. Life and death at Dmanisi, Georgia: Taphonomic signals from the fossil mammals. J Hum Evol 171:103249.

Ugai K, Yagi H, Wakai A. 2012. Earthquake-Induced Landslides: Proceedings of the International Symposium on Earthquake-Induced Landslides, Kiryu, Japan, 2012. Springer Science & Business Media.

Uribelarrea del Val D, Domínguez-Rodrigo M. 2017. Geoarchaeology in a meandering river: A study of the BK site (1.35 Ma), Upper Bed II, Olduvai Gorge (Tanzania). Palaeogeogr Palaeoclimatol Palaeoecol 488:76–83.

Uribelarrea D, Martín-Perea D, Díez-Martín F, Sánchez-Yustos P, Dominguez-Rodrigo M, Baquedano E, Mabulla A. 2017. A reconstruction of the paleolandscape during the earliest Acheulian of FLK West: The co-existence of Oldowan and Acheulian industries during lowermost Bed II (Olduvai Gorge, Tanzania). Palaeogeogr Palaeoclimatol Palaeoecol 488:50–58.

Uribelarrea D, Perea DM, Díez-Martín F. 2019. A geoarchaeological reassessment of the co-occurrence of the oldest Acheulean and Oldowan in a fluvial ecotone from lower middle Bed II (1.7 ma) at Olduvai Gorge (Tanzania). Quaternary International 526: 39–48.

Villa P. 1990. Torralba and Aridos: elephant exploitation in Middle Pleistocene Spain. J Hum Evol 19:299–309.

Villa P, Bartram L. 1996. Flaked bone from a hyena den. Paléo 8:143–159.

Villa, P. and Mahieu, E., 1991. Breakage patterns of human long bones. Journal of human evolution, 21(1), pp.27–48.

Villa P, d’Errico F. 2001. Bone and ivory points in the Lower and Middle Paleolithic of Europe. J Hum Evol 41:69–112.

Villa P, Soto E, Santonja M, Pérez-González A, Mora R, Parcerisas J, Sesé C. 2005. New data from Ambrona: closing the hunting versus scavenging debate. Quat Int 126-128:223–250.

Wasowski J, Keefer DK, Lee C-T. 2011. Toward the next generation of research on earthquake-induced landslides: Current issues and future challenges. Eng Geol 122:1–8.

Wood BM. 2010. Household and Kin Provisioning by Hadza Males. Harvard University.

Wood BM, Marlowe FW. 2013. Household and kin provisioning by Hadza men. Hum Nat 24:280–317.

Yravedra J, Domínguez-Rodrigo M, Santonja M, Pérez-González A, Panera J, Rubio-Jara S, Baquedano E. 2010. Cut marks on the Middle Pleistocene elephant carcass of Áridos 2 (Madrid, Spain). J Archaeol Sci 37:2469–2476.

Yravedra J, Rubio-Jara S, Panera J, Martos JA. 2019. Hominins and Proboscideans in the Lower and Middle Palaeolithic in the Central Iberian Peninsula. Quat Int 520:140–156.

Yravedra J, Rubio-Jara S, Panera J, Uribelarrea D, Pérez-González A. 2012. Elephants and subsistence. Evidence of the human exploitation of extremely large mammal bones from the Middle Palaeolithic site of PRERESA (Madrid, Spain). J Archaeol Sci 39:1063–1071.

Yravedra, J., Courtenay, L., Gutiérrez-Rodríguez, M., Reinoso-Gordo, M., Saarinen, J., Égüez, N., Luzón, C., Rodríguez-Alba, J., Solano, J., Titton, S., Montilla-Jiménez, E., Cámara-Donoso, E., Herranz-Rodrigo, D., Estaca, V., Serrano-Ramos, A., Amorós, G., Azanza, B., Bocherens,H., DeMiguel,D., Fagoaga, A., García-Alix,A.,González-Quiñones,J., Jiménez-Espejo, F., Kaakinen, H., Munuera,M., Ochando, J., Piñero, P., Sánchez-Bandera, C., Viranta, S., Fortelius, M.,Agustí,J., Hugues-Alexandre Blain, Carrión, J., Barsky, D., Oms, O., Mallol,C., Jiménez-Arenas,J.M. 2024. Not seen before. Unveiling depositional context and Mammuthus meridionalis exploitation at Fuente Nueva 3 (Orce, southern Iberia) through taphonomy and microstratigraphy. Quaternary Science Reviews,Volume 329, 10.1016/j.quascirev.2024.108561.

Zutovski K, Barkai R. 2016. The use of elephant bones for making Acheulian handaxes: A fresh look at old bones. Quat Int 406:227–238.

