## Supplementary Information for "Earliest Evidence of Elephant Butchery at Olduvai Gorge (Tanzania) Reveals the Evolutionary Impact of Early Human Megafaunal Exploitation"

**Index**

EAK EXCAVATION AND LANDSCAPE DISTRIBUTION
OF MEGAFAUNAL GREEN-BROKEN BONES........................................................ 3

STRATIGRAPHICALLY ASSOCIATED GREEN-BROKEN
PROBOSCIDEAN BONES......................................................................................... 5

GREEN-BROKEN SPECIMENS WITH UNCERTAIN
STRATIGRAPHIC ATTRIBUTION................................................................................16

GREEN-BROKEN BONES AT EAK............................................................................ 17

THE SPATIAL STATISTICAL ANALYSIS OF EAK...................................................... 19

THE TECHNOLOGICAL ANALYSIS OF EAK..............................................................21

ORIENTATION OF MATERIALS AT EAK...................................................................29

CARNIVORE DAMAGE ON AN ADULT GIRAFFE
(CHOBE NATIONAL PARK, BOTSWANA)...............................................................30

THE ISSUE OF BONE TOOLS..................................................................................33

**EAK EXCAVATION AND LANDSCAPE DISTRIBUTION OF MEGAFAUNAL GREEN-BROKEN BONES**

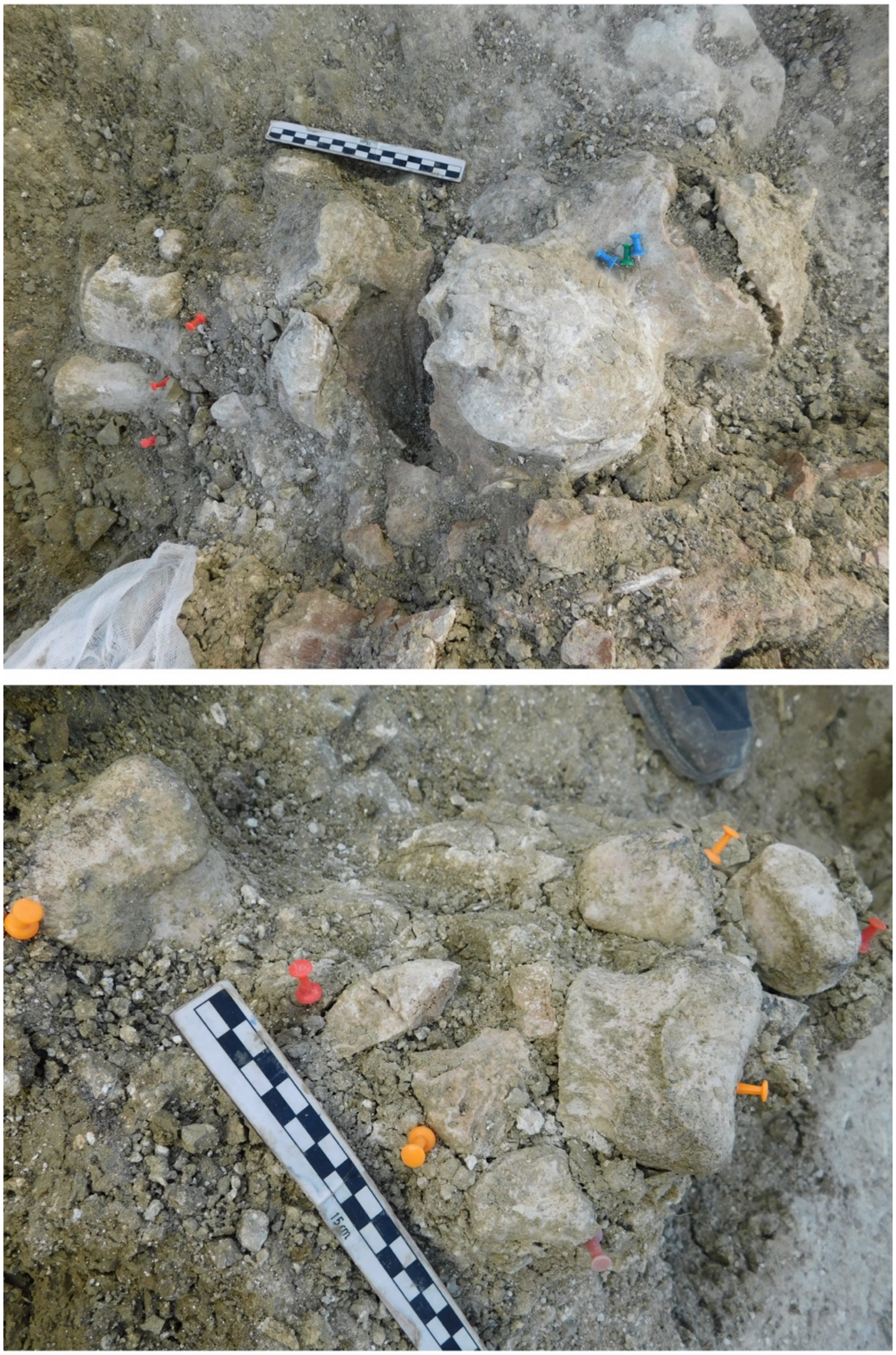

*Fig S1. Excavation at EAK of the partial articulated rear foot of the juvenile elephant, displaying the calcaneum with unfused epiphysis (upper) and articulated phalanges (lower).*

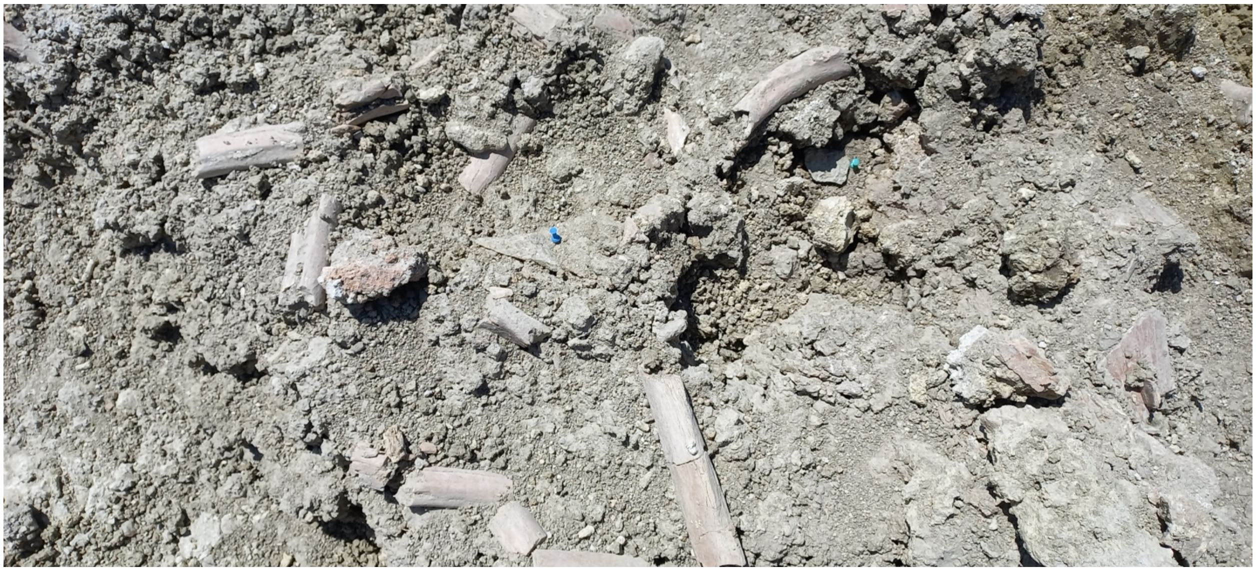

*Fig S2. Fragmented and anatomically connected ribs from the proboscidean individual at EAK.*

| | NISP | MNE |
|---|---|---|
| cranium | 12 | 1 |
| mandible | | |
| teeth | 44 | 5 |
| vertebrae | 1 | 1 |
| ribs | 51 | 8 |
| pelvis | 11 | 1 |
| scapula | | |
| humerus | | |
| radius | | |
| ulna | 1 | 1 |
| carpals | | |
| metacarpals | | |
| femur | 5 | 2 |
| tibia | 2 | 2 |
| patella | 1 | 1 |
| calcaneum | 2 | 1 |
| astragalus | 1 | 1 |
| tarsals | 5 | 5 |
| metatarsals | 5 | 5 |
| phalanges | 12 | 12 |

*Table S1. Skeletal representation of the elephant carcass at EAK.*

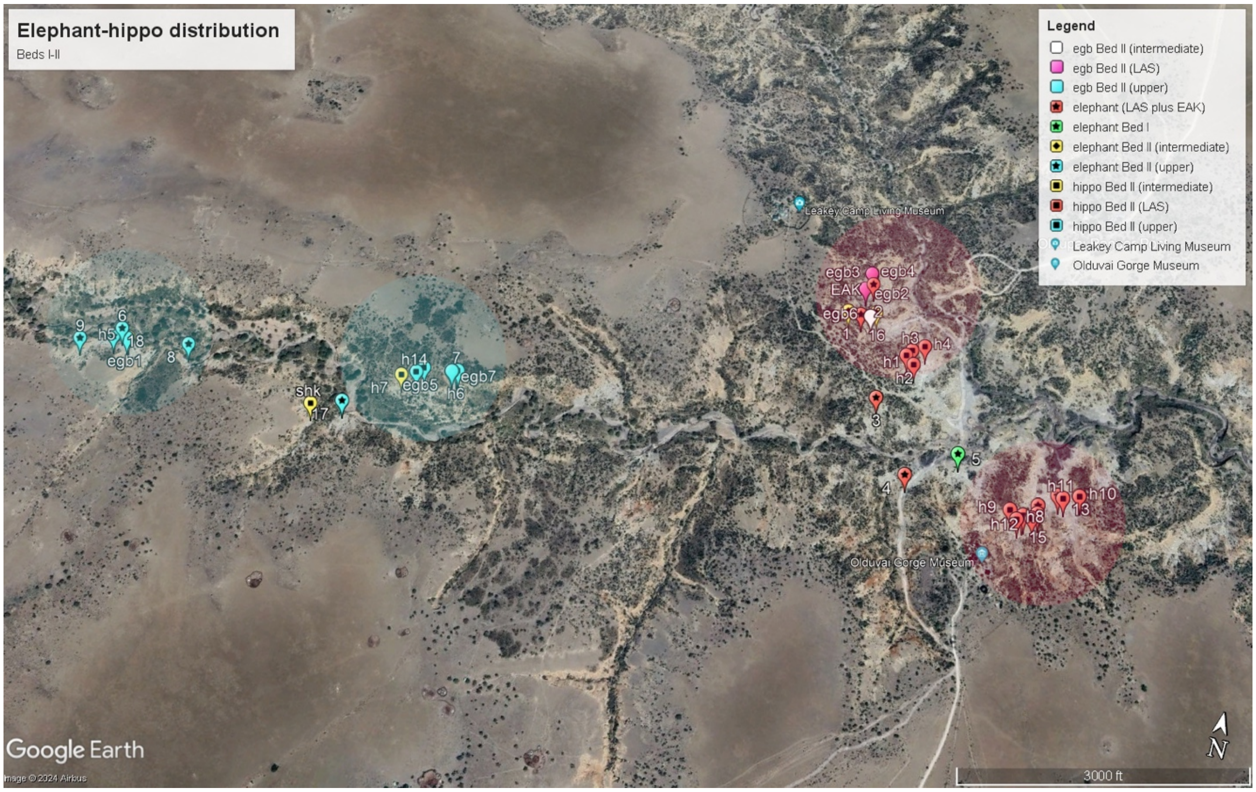

*Fig S3. Distribution of elephant and hippopotamus carcasses at the main section of Olduvai Gorge, documented in the survey conducted in June 2024. The lower (red), intermediate (yellow) and upper (blue) stratigraphic units display a clustered distribution of carcasses. Notice the spatial overlap of those megafaunal clusters with the dense landscape-scale concentrations of stone artifacts (in faded red for the LAS lower unit, and faded blue for the upper unit). Elephant carcasses appear numbered and hippopotamus carcasses start with "h" before the number. EGB, elephant green-broken bone. The megafaunal site of TK* Sivatherium *is outside the map.*

**STRATIGRAPHICALLY ASSOCIATED GREEN-BROKEN PROBOSCIDEAN BONES**

**(see Figure S3)**

A small collection of green-broken elephant bones was retrieved from the lower and upper Bed II units. They appear here as example of green-broken megafaunal bones found in the three paleolandscape units targeted. One of them, a proximal femoral shaft found within the LAS unit, has all the traces of having been used as a tool (Figure 6). It displays green breaks on all its sides, with conchoidal fractures observed both on the cortical and medullary surfaces (Figure S5-S7). This indicates multiple dynamic loading events, reflected in abundant step fractures, ripple marks and multiple, same-direction curved outlined step micro-conchoidal scars on the same front of its proximal end. This, together with their association to a polished tip bearing more abrasion than the rest of the bone, is suggestive of the bone specimen having been used as a tool (Fig 6). An equally compelling case of anthropogenic agency was found on another specimen found a few meters away (probably suggesting butchery of the same individual elephant), with a complete and smooth set of two spiral overlapping green fractures (Fig S4 middle), and a series of step fractures on the breakage plane of the alternative break (Fig S4 lower), finalized with a percussion mark on the cortical surface of the spiral break (Fig S4 upper).

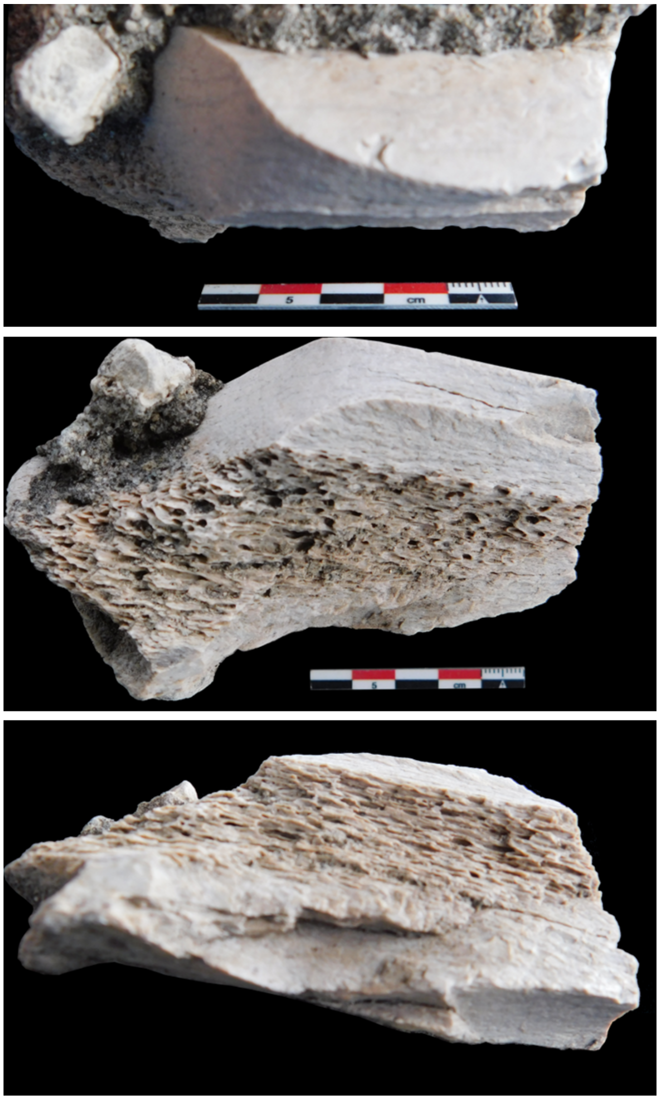

*Figure S4. Example of elephant green-broken limb bone shaft displaying percussion mark (upper) and overlapping reflected/stepped scars on its medullary surface resulting from hammerstone breaking.*

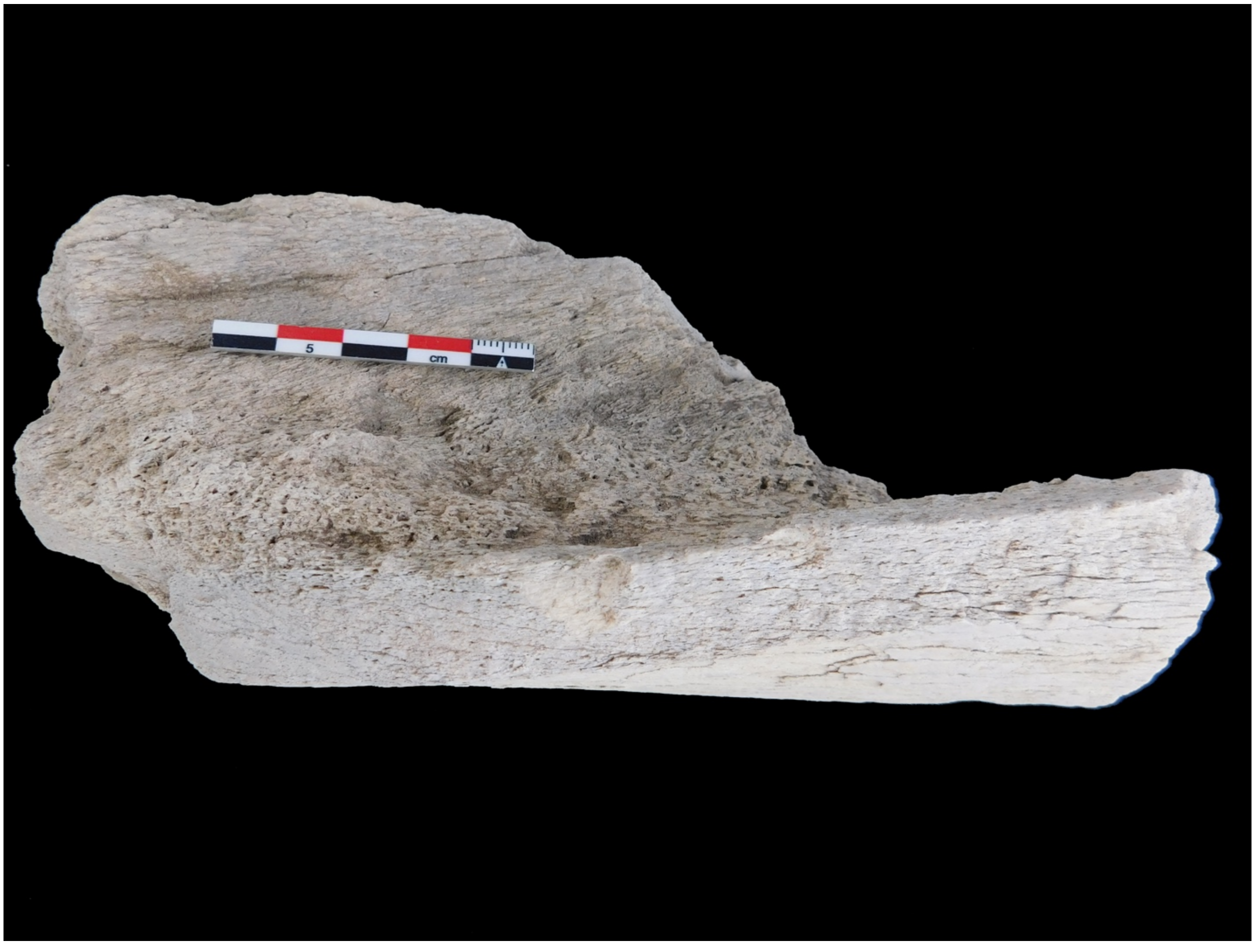

*Figure S5. EGB1: Proboscidean limb bone shaft with green breaks found at BK (upper unit).*

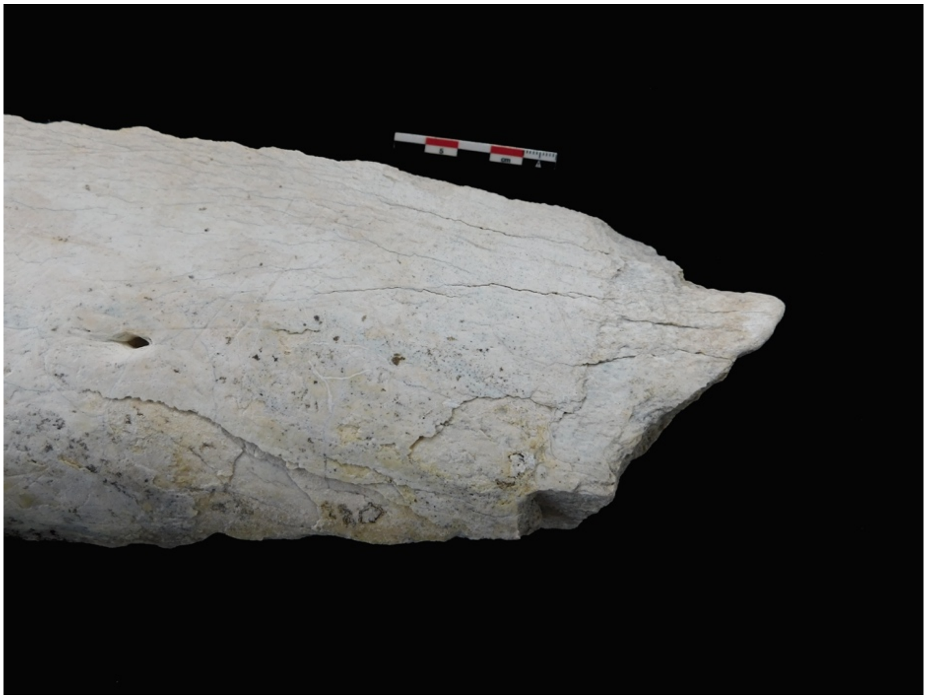

*Figure S6. EGB 2: Proboscidean femoral shaft with green breaks and a distal portion showing a combination of curved scars and a polished tip, most likely polished anthropogenically.*

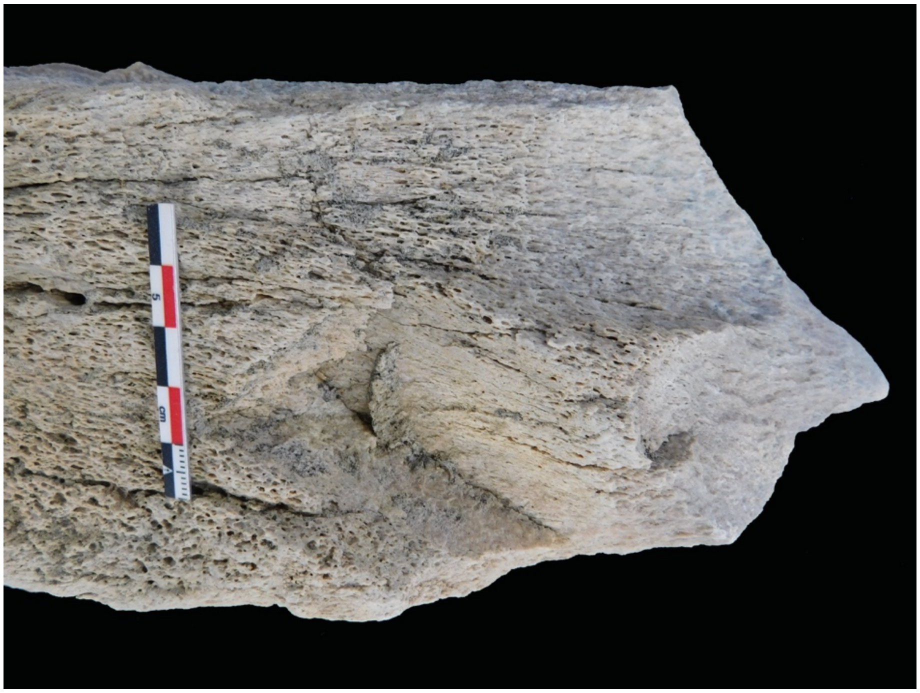

*Figure S7. EGB 2: Medullary view of the same proboscidean specimen displaying green breaks and a step fractures on its proximal portion.*

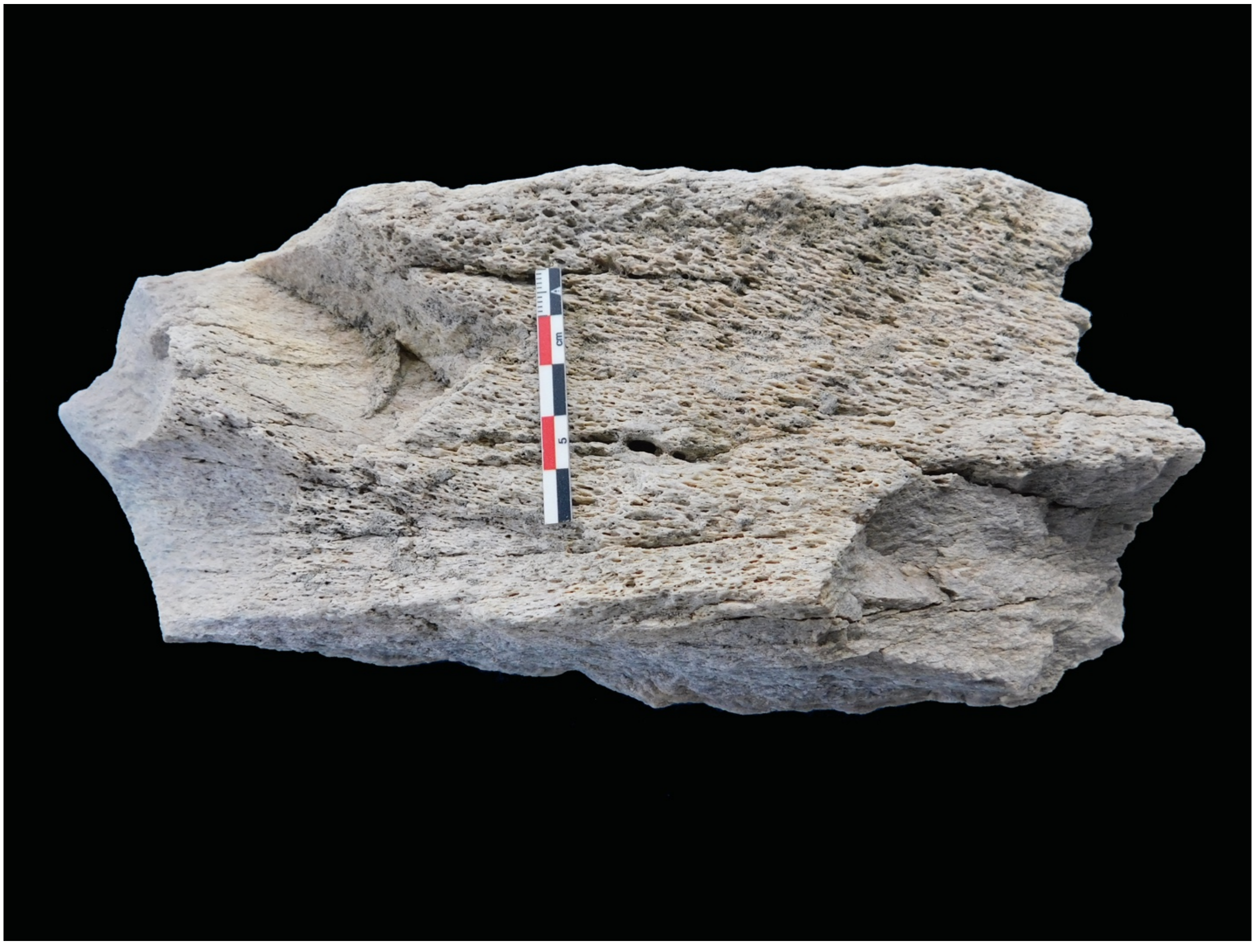

*Figure S8. EGB 2: Same specimen showing green fractures on all its sections, as seen from a medullary perspective. All breaks show sharp edges in contrast with the polished tip.*

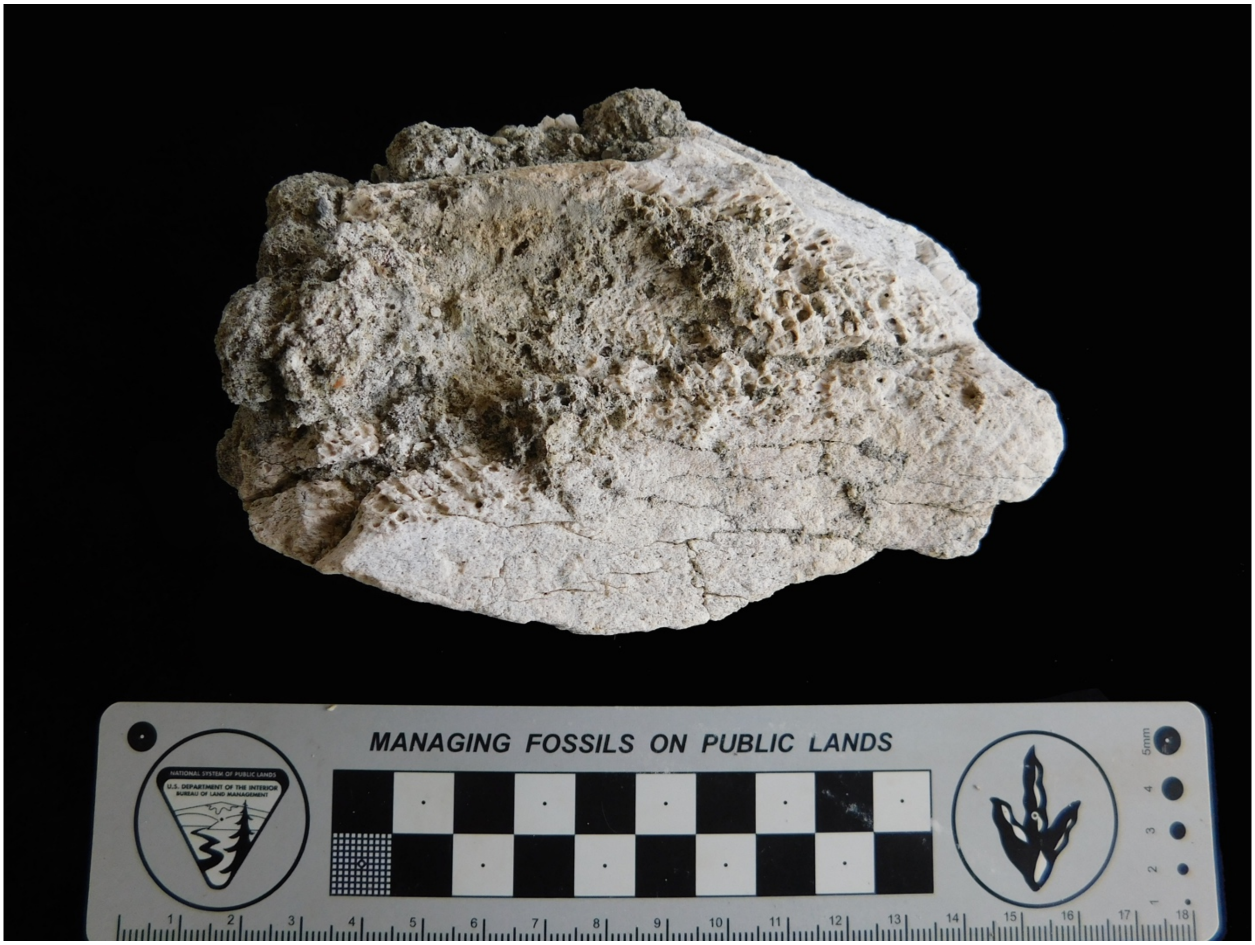

*Figure S9. EGB3: Proboscidean limb shaft with smooth green breaks. It was found in the LAS lower unit.*

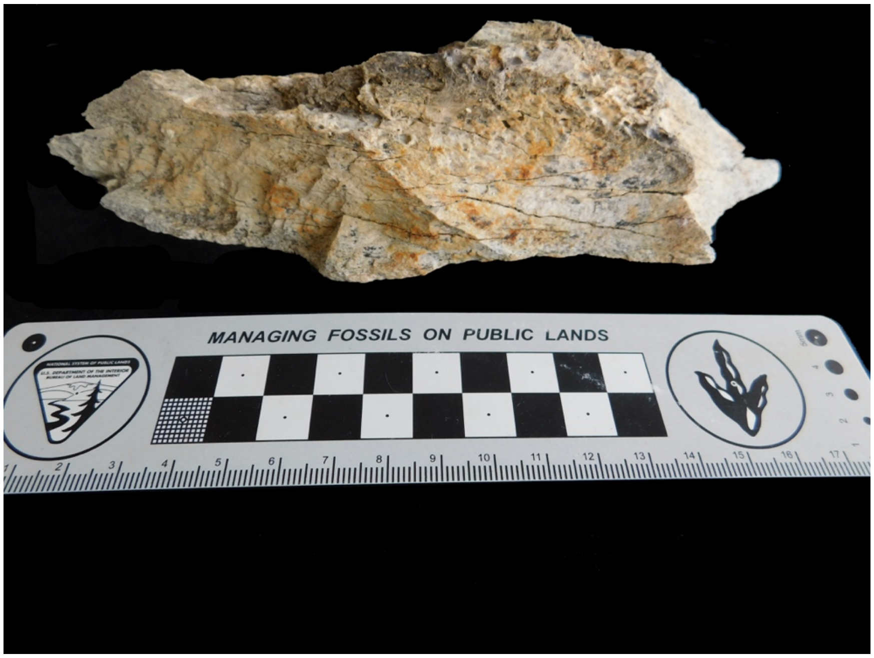

*Figure S10. EGB4: Proboscidean limb shaft fragment with green breaks showing multiple overlapping conchoidal scars and hackle marks typical of dynamic loading. It was found in the LAS lower unit.*

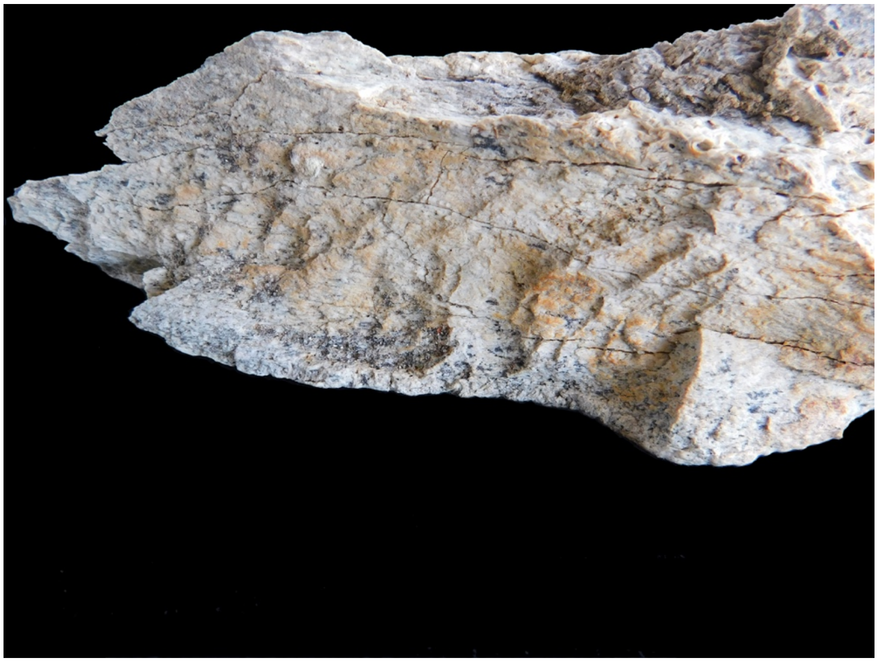

*Figure S11. EGB4: Detail of the hackle marks of the same specimen, typical of dynamic loading.*

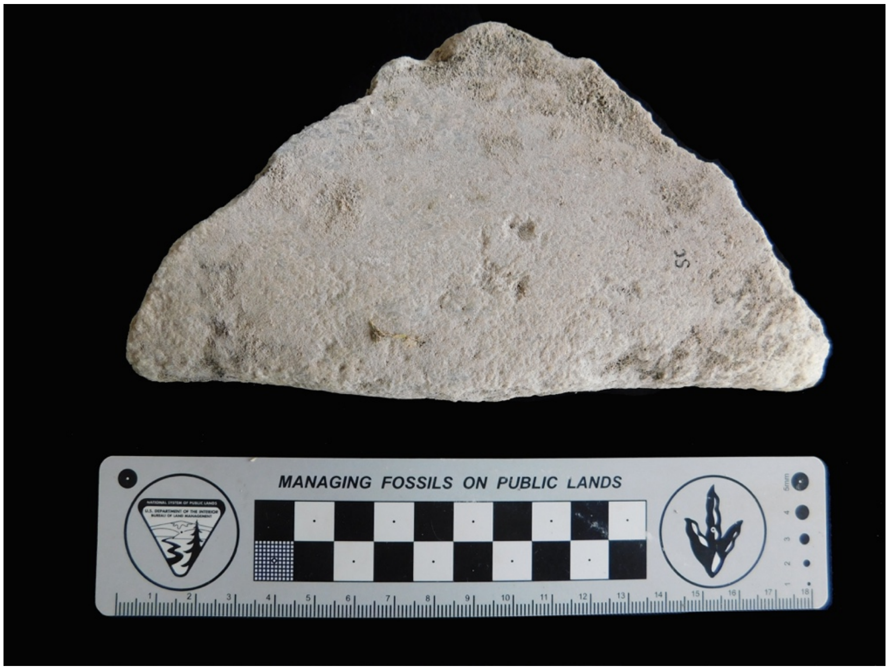

*Figure S12. EGB5: Probable proboscidean flat bone found at SC (upper unit). It has attributes of green breakage and some pitting on its cortical surface, possibly resulting from percussion.*

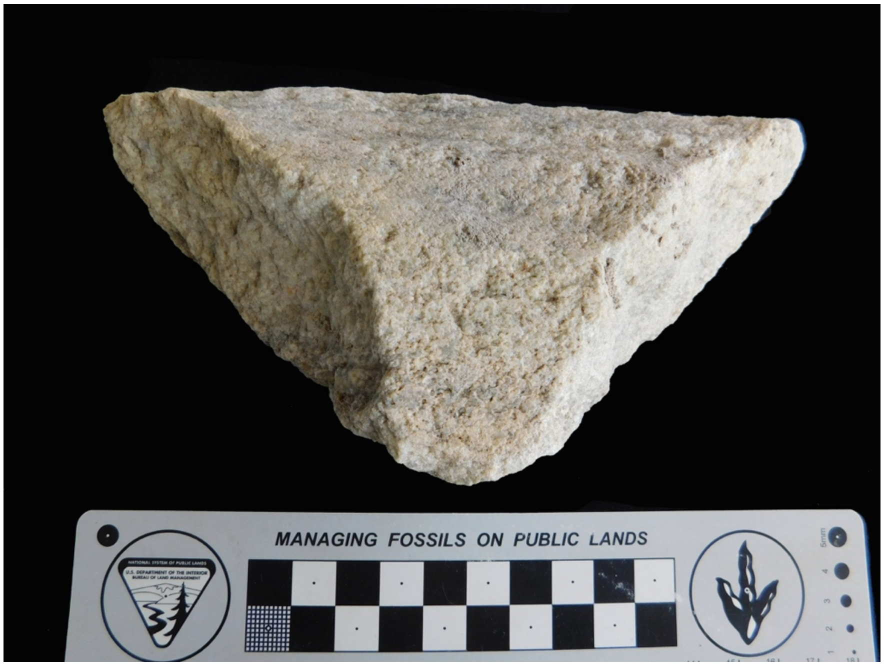

*Figure S13. EGB 5: same specimen seen from the other side.*

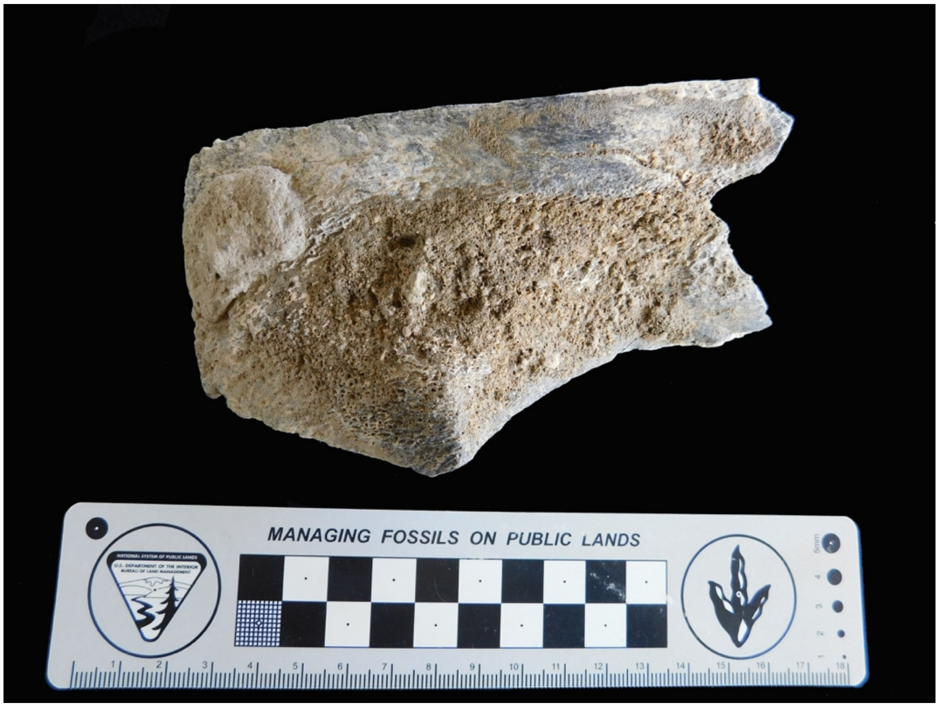

*Figure S14. EGB6: Medullary view of possible proboscidean limb shaft fragment with green break morphology, found in the intermediate unit in the vicinity of FLK west.*

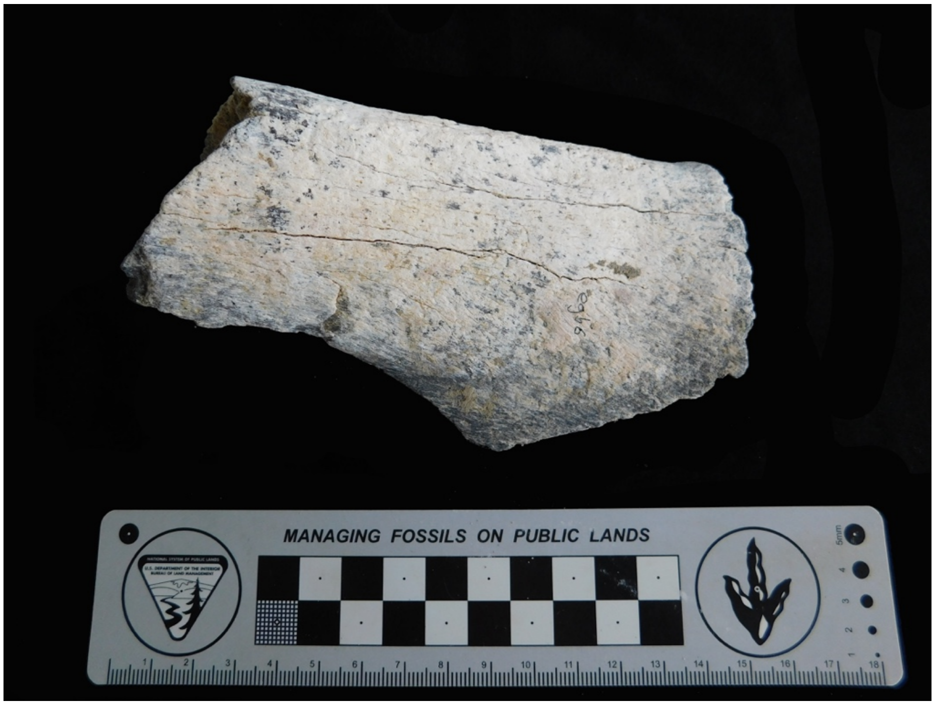

*Figure S15. EGB 6: same specimen seen from its cortical side. Notice the pitting on its surface near the notch.*

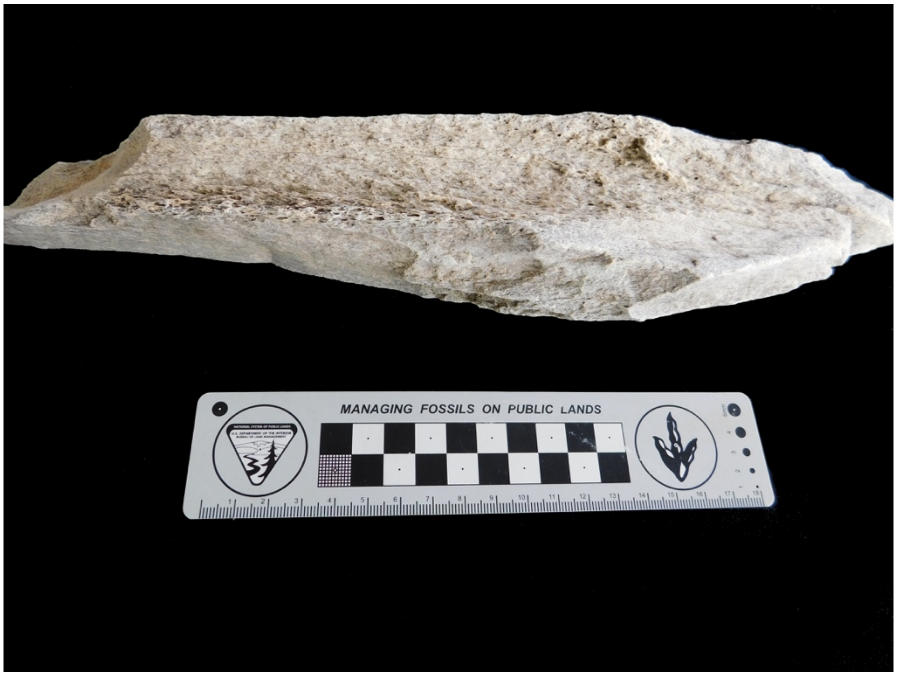

*Figure S16. EGB7: Proboscidean limb shaft fragment (probably from a femur) with green break attributes, showing multiple overlapping conchoidal scars (center) found at SC (upper unit).*

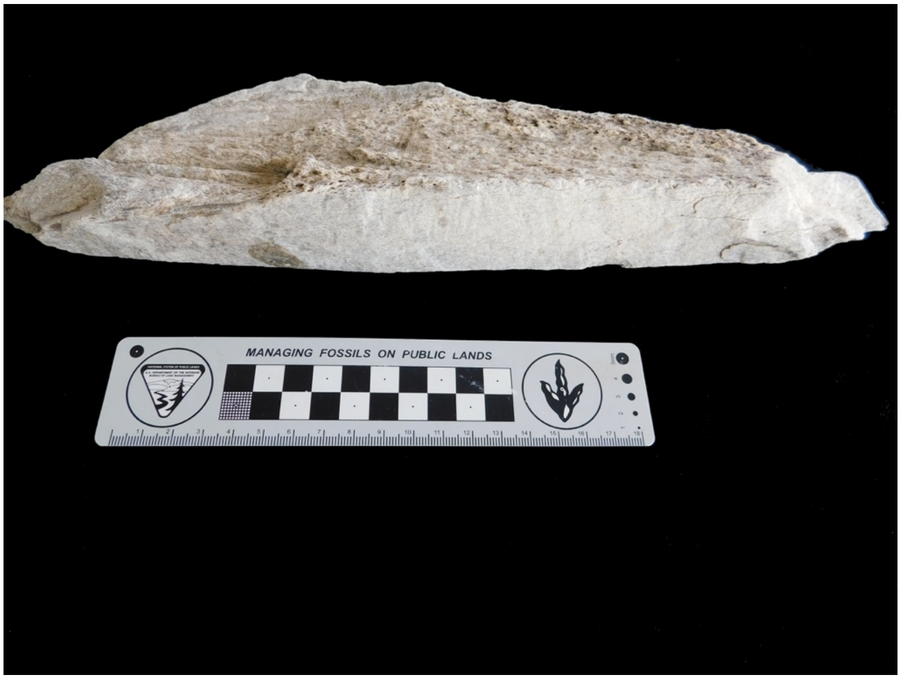

*Figure S17. EGB7: Other side of the same proboscidean limb shaft fragment with attributes of green-bone breakage, such as smooth fracture surface  and conchoidal scars, found at SC (upper unit).*

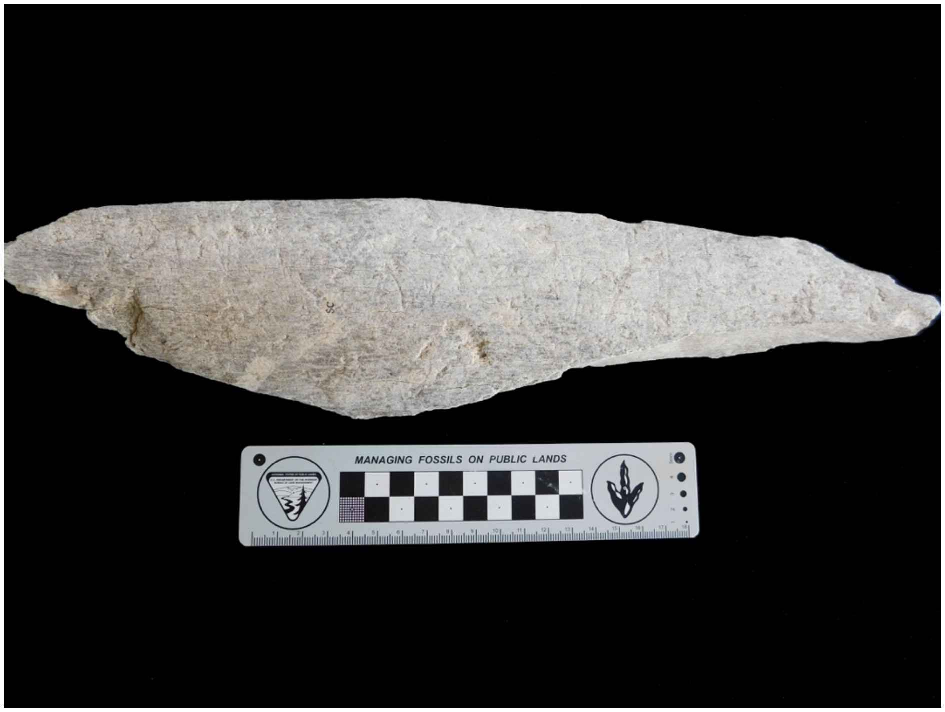

*Figure S18. EGB7: Same proboscidean limb shaft fragment, cortical view. Note the marks from abrasive agencies, plus some pitting by both lower edges of breakage planes suggestive of the potential dynamic loading effector.*

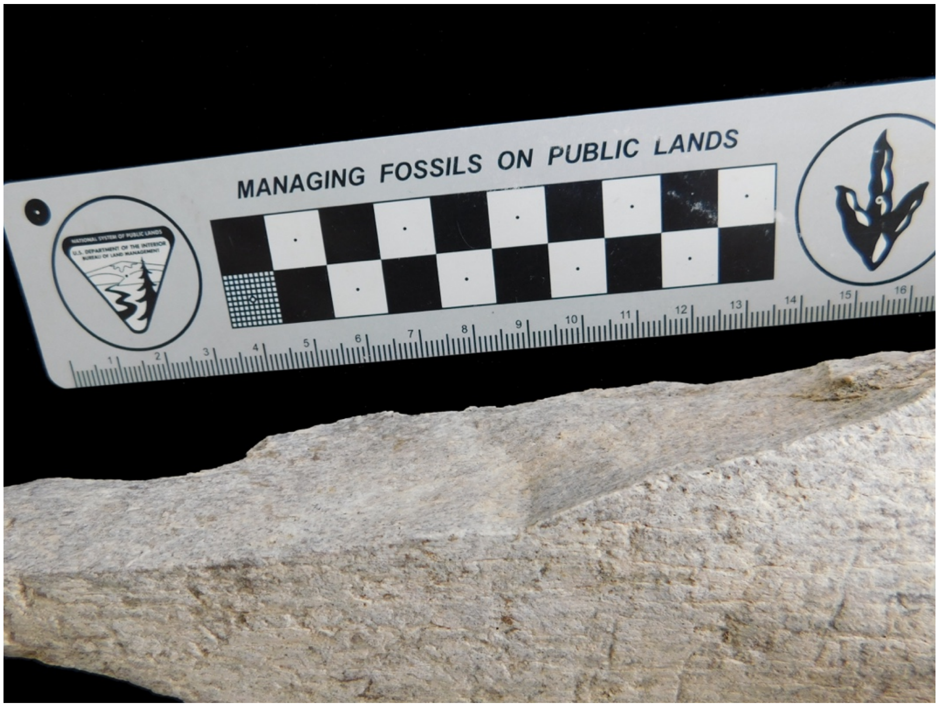

*Figure S19. EGB7: Detail of the same limb shaft fragment showing the longest smooth fracture surface typical of a green break with inflection point in the middle.*

### GREEN-BROKEN SPECIMENS WITH UNCERTAIN STRATIGRAPHIC ATTRIBUTION

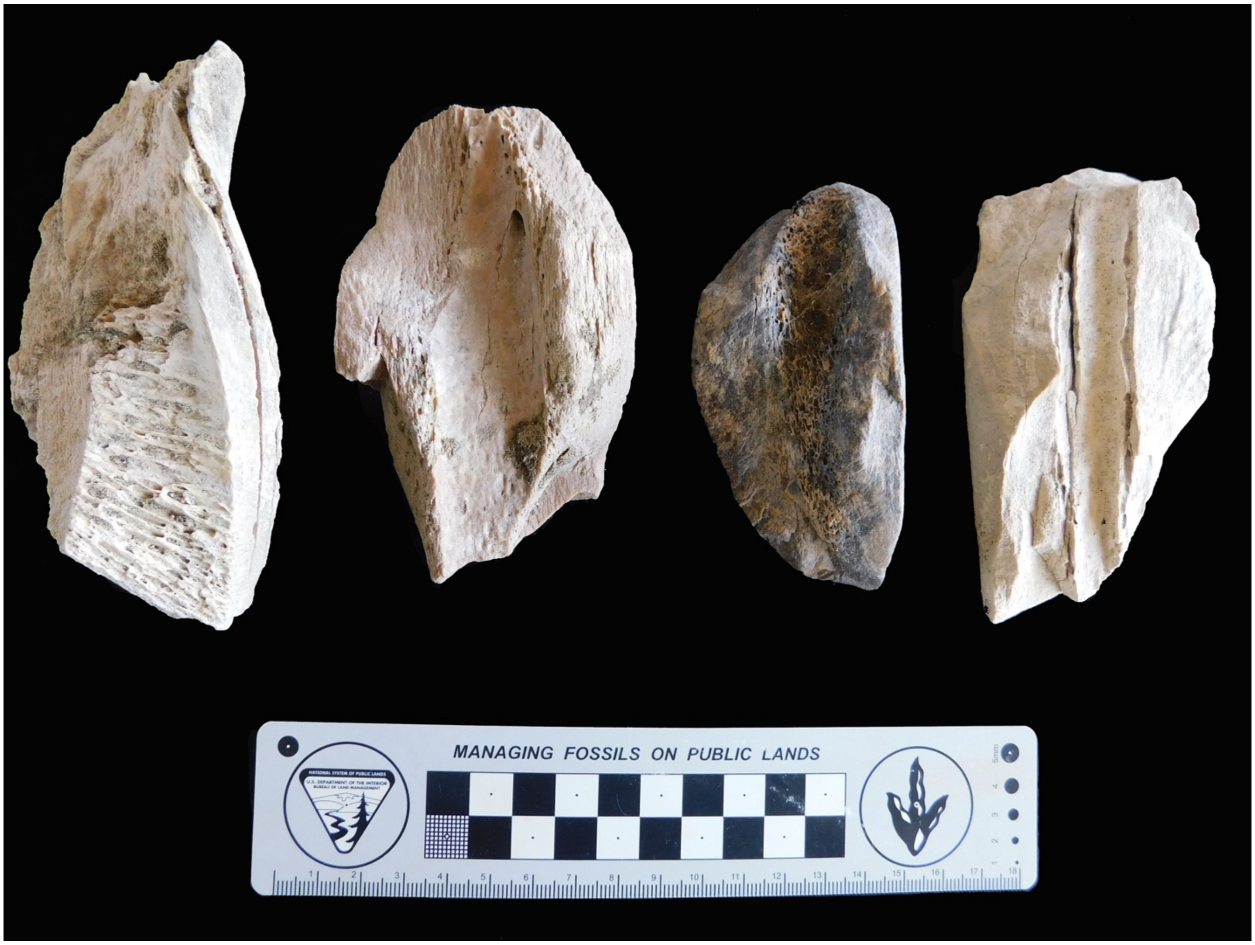

*Figure S20. Example of other megafaunal long bone shafts showing green breaks.*

### GREEN-BROKEN BONES AT EAK

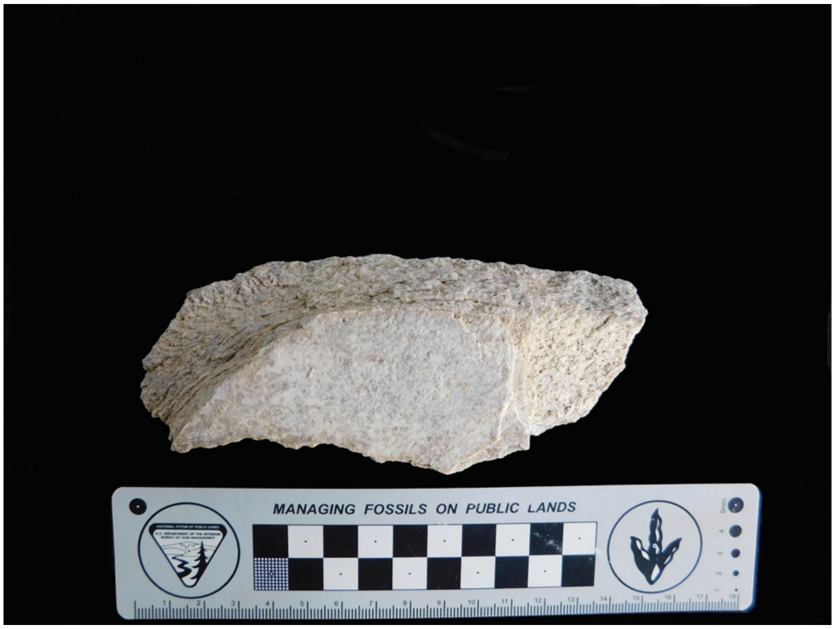

*Figure S21. Elephant-sized flat bone fragment with green-bone curvilinear break*

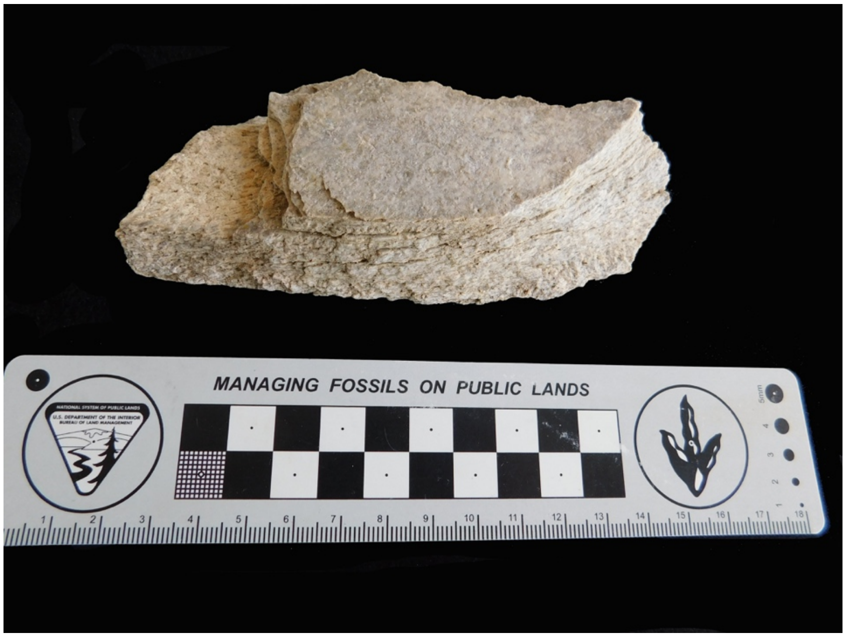

*Figure S22. Same specimen with a different view.*

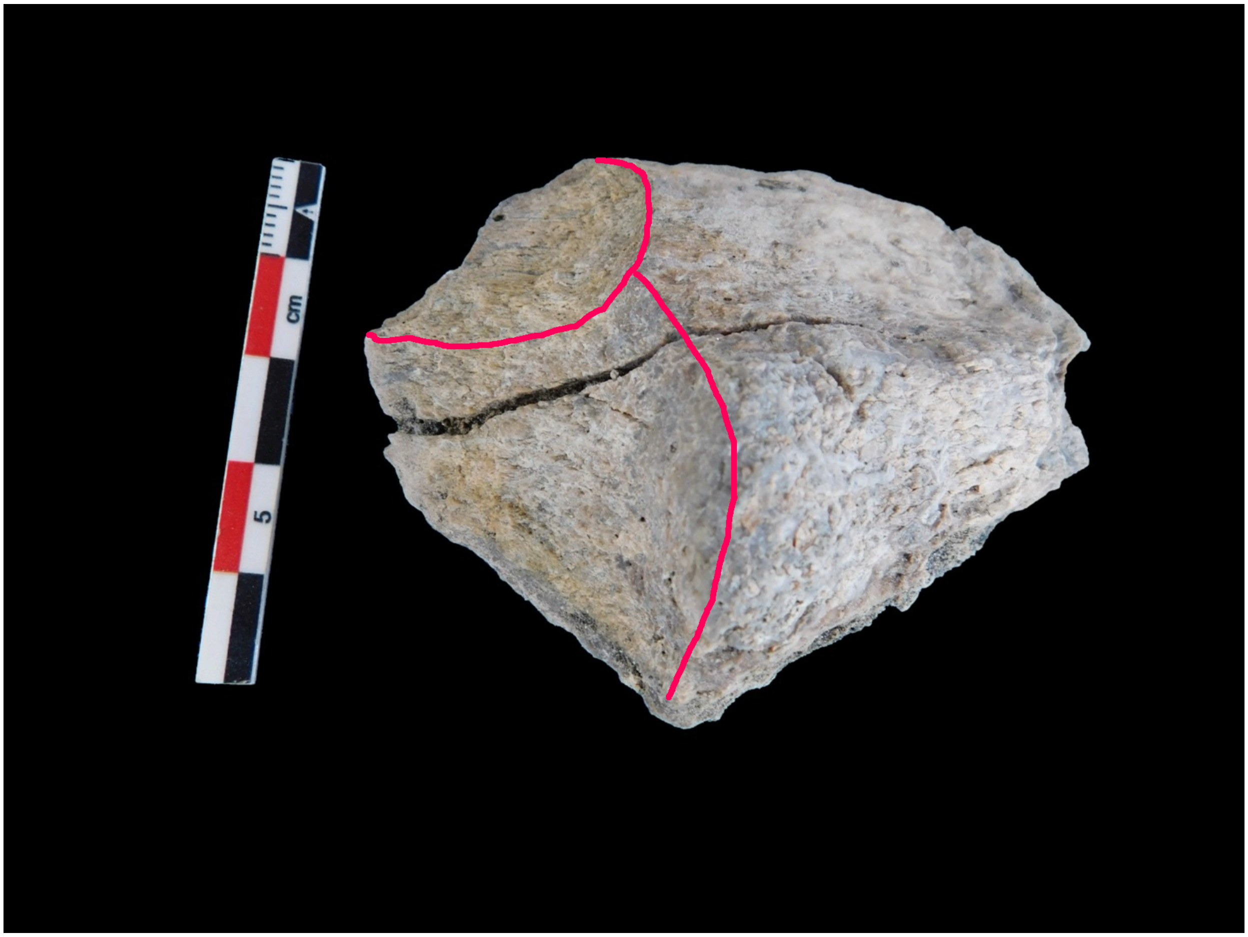

*Figure S23. Proboscidean limb bone fragment found at EAK, with morphology of a green-bone break. Red lines show the outline of the overlapping scars.*

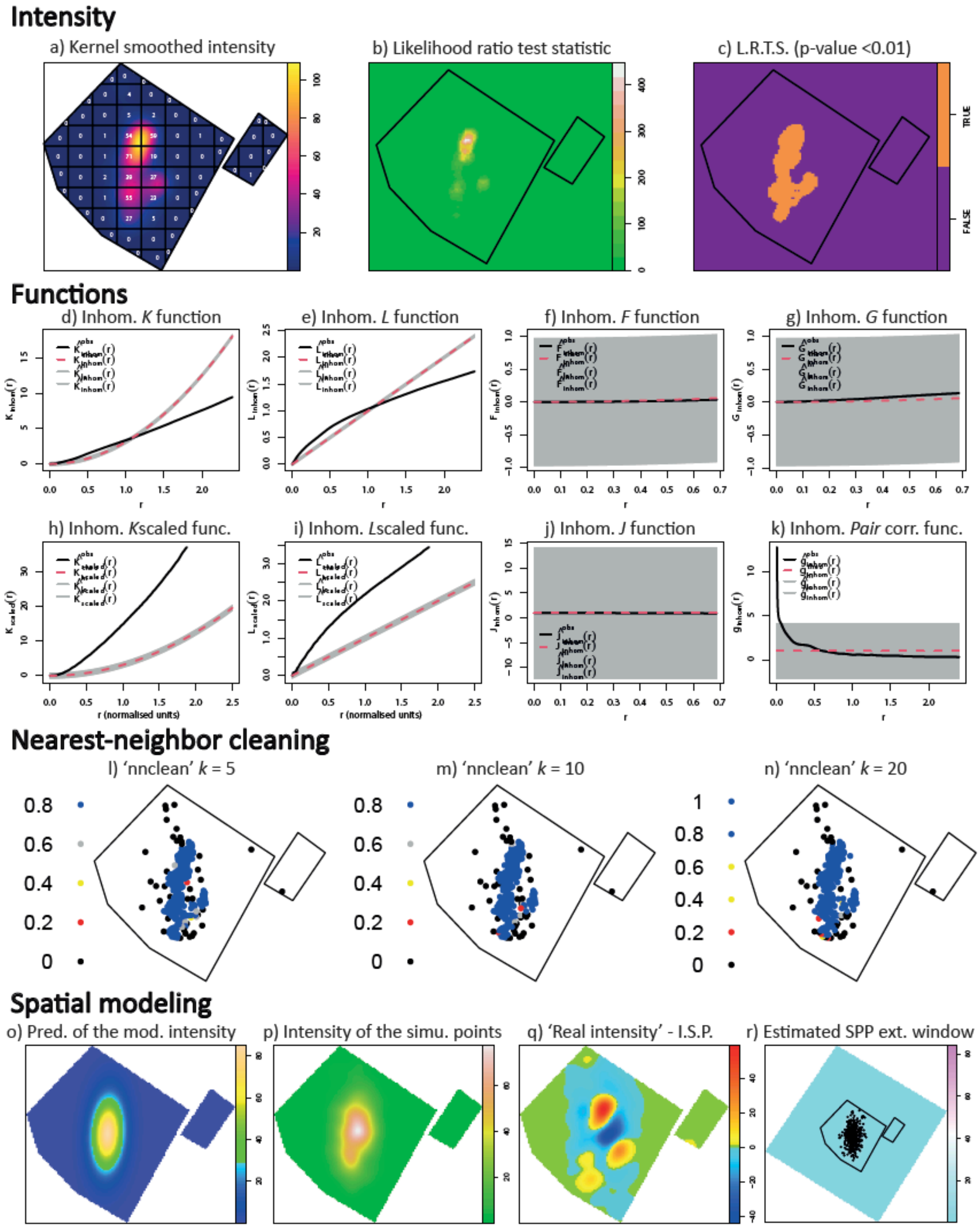

*Figure S24. Mapping of intensity, nearest-neighbor cleaning, and spatial modeling of the SPP of bones at EAK, including graphs of the inhomogeneous second-order functions.*

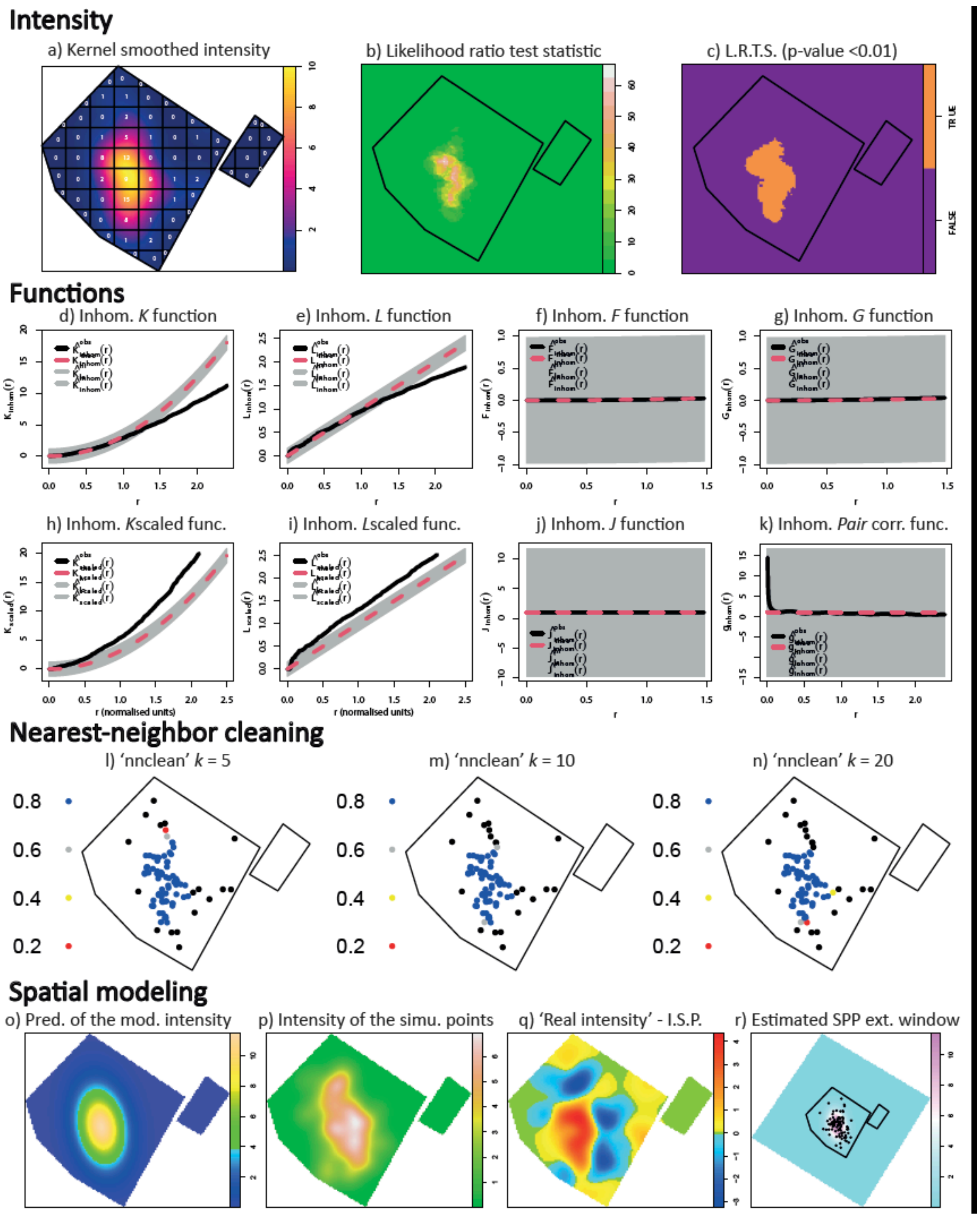

*Figure S25. Mapping of intensity, nearest-neighbor cleaning, and spatial modeling of the SPP of lithics at EAK, including graphs of the inhomogeneous second-order functions.*

### The technological analysis of EAK

*Materials and methods of the lithic analysis*

A collection of 80 lithic specimens has been retrieved from EAK. For the study of these materials we have taken in consideration the following technological categories and procedures implied in the analysis (Diez-Martín et al. 2009a, 2009b, 2011, 2017, 2024): (a) Unmodified specimens are cobblestones devoid of any sign of anthropogenic manipulation; (b) percussion materials, include Hammerstones or cobbles showing battering and/or scarring on their surface; (c) Cores. Handheld cores have been classified according to the combination of the following traits: number of reduction surfaces (unifacial, bifacial and multifacial); number of striking platforms (unipolar, bipolar, multipolar); arrangement of the identified knapping series (lineal, opposed, orthogonal, centripetal). Bipolar cores have been defined through the technical traits related to anvil percussion; (d) Detached products include both diagnostic bipolar and handheld complete and broken flakes. Technical traits recorded in flakes include reduction stages (according to the variable retention of cortex on dorsal areas and striking platforms), striking platform type, and dorsal scar arrangements;(e) Waste material includes the by-products of the knapping processes: non-diagnostic flake fragments with signs of direct handheld percussion, shatter or blocky/angular undetermined detached fragments, core-like fragments and ≤20 mm debris.

*Results*

All the specimens but one have been preserved in mint fresh conditions, lacking any signs of post-sedimentary alteration (R0). An indeterminate shatter fragment shows severe abrasion on surfaces and edges (R3). The bulk of the sample is constituted by quartz specimens (96%), while the sum is formed by few lavas (two basalt cobbles and a phonolite flake). **Table S2** shows the distribution of lithics sorted by category and raw materials. In general, the EAK collection is a meager sample with a limited expression of knapping behaviors, as all the categories with technological significant are represented by few specimens. Non-diagnostic waste is the most significant category, mostly ≤20 mm debris and flake fragments.
In the following description, lithic categories are grouped by operational sequences (techno-economic actions sharing common goals):

1) Unmodified material. Natural and non-modified materials are represented by a low-quality vesicular basalt amorphous cobble (87x59x52 mm, 296 g) showing no sign of modification on its surfaces and, thus, devoid of anthropogenic use.

2) Percussion activities. Percussive actions are represented by a single high-quality basalt hammerstone. The specimen is oval and shows significant signs of battering on its whole perimeter (proximal and distal ends, and lateral areas) (**Figure S26.2**).

3) Production of small and medium-sized flakes. Fractions of the flake production sequences have been documented on quartz and, very partially, on phonolite:

a) Cores represent 5% of the total sample. Two knapping methods have been identified on quartz blanks. Direct handheld percussion is represented by an exhausted multifacial specimen showing at least 10 negative scars detached from multiple planes. Considering its advanced stage of exploitation, the last flakes produced from this core are significantly small (mean length and breadth 24x28 mm) (**Figure S26.4**). The bipolar on anvil technique is represented by three cubic quartz specimens in an advanced phase of reduction (mean 66x52x41 mm, 225 g), most probably tabular slab fragments, showing opposed bipolar scars detached from platform and base that, in two cases, show circular or semi-circular core rotation (**Figure S26.3 and S26.5**). Two specimens show signs of frosting on their platform areas. When properly identified in negative scars (n=4), flakes detached from bipolar (mean 34x40 mm) cores are larger (significantly wider) than those observed in the freehand multipolar specimen (mean 24x28 mm) and one of the flakes removed from a bipolar core can be included within the medium-sized flake group (breath 63 mm) (**Figure S26.1**).

b) Detached material. Most of the complete and broken diagnostic flakes retrieved have been assigned to direct handheld percussion (83%), while the sum is represented by specimens showing clear traits of bipolar knapping. As for the former method, the few broken specimens show longitudinal axial snaps (a very common fracture accident observed in the Naibor Soit raw material), although a side-proximal snap has also been identified. The mean size of broken flakes is 32x23x10 mm and 8 g. Most complete flakes retrieved fall within the small size group (≤50 mm of maximum length), as mean flake size is 33x29x11 mm. With a length of 58 mm, only one retrieved flake exceeds this range and, thus, can be included in the medium-sized flake range (**Figure S27.2**). Size and mass parameters of complete flakes are detailed in **Table S3**. The phonolite flake included in this category fits well within the quartz size pattern and with the most representative technical traits described for quartz specimens (**Figure S27.1**). Regarding their position in the reduction sequence, 90% of the flake sample is included in the full production stage, this is to say, flakes with no cortex retention on both dorsal surface and butt (Type 6). The remaining flakes partially retain cortex on the dorsal surface (Type 5, n=1) or striking platform (Type 3, n=1). Striking platforms are clearly dominated by unifaceted butts (73.7%), although other non-cortical butts are also represented (lineal and point striking platforms). A single cortical butt has been identified in a specimen with no cortex on the dorsal area (Type 3). Mean breadth and thickness of striking platforms is 19x8 mm (range 11-31 mm and 6-12 mm, respectively). Flake morphology in the frontal plane is preferentially rectangular (33%), quadrangular (28%) and triangular (22%) (**Figure S27.3**). With respect to dorsal patterns, mean negative scars observed in dorsal areas is 2.17 (interval 1-5). The directional arrangements of flakes, excluding those showing a single dorsal scar (n=4), follow a variety patterns, although the most numerous are bidirectional opposed (longitudinal and lateral) (**Figure S27.4 and S27.6**) and orthogonal (partial or full) (**Figure S27.5**), both patterns accounting for 44% of the identified cases, respectively. These dorsal models fit well within the reduction strategies observed in the core sample. Natural and potentially usable cutting edges on flakes tend to be located on the side (60%), being specifically lateral (45%) or bilateral. Distal (35%) and transversal (5%) locations are also documented. Mean cutting edge length per flake is 32 mm (30 mm in lateral specimens, 58 mm in bilateral, and 27 mm in distal). The case of the single phonolite flake deserves a comment. As no volcanic cores have been retrieved from the site, this specimen might well have arrived to the site already produced from other area. This impression is reinforced by the fact that, in agreement with the dimensional pattern observed among the quartz flake collection, this small product shows a potentially usable cutting edge of 92 mm (summing bilateral and distal edges), far beyond mean edge length in quartz or even the maximum length recorded in specimens detached from this raw material (56 mm). A small collection of flakes clearly detached by means of anvil technique has been retrieved (with identification of platform and base areas, opposed detachments, and irregular ventral surfaces). Mean size of these flakes is 36x30x15 mm and 17 g, which shows that they tend to be particularly longer and thicker than freehand flakes. One specimen shows frosting on the basal area.

c) Waste constitutes the most abundant group of lithics. Most likely these pieces are the by-products of both freehand and bipolar reduction. All these specimens have been obtained from quartz blanks. Most waste (47%) is constituted by small debris (mean size 16.6x11x6 mm and 1.5 g). Flake fragments represent 26% of this category and can be tentatively ascribed to handheld knapping, although no clear technical traits can be identified due to the relevance of snapping accidents in these pieces. Mean size of flake fragments is 25x20x11 mm and 4.5 g of mass. Shatter includes larger indeterminate positives with a blocky morphology whose mean size is 27x20x18 mm and 7 g. Finally, a small group of unspecific core fragments include three angular, cubic and non-diagnostic specimens that might be related to the exploitation of tabular slabs. Mean size of these specimens is 51x27x26 mm and a mass of 46 g.

*Conclusions*

A small lithic collection has been retrieved from EAK. Although percussion activities carried out at the site have been documented, they are residual. A single hammerstone, with intense signs of battering, evidences this behavior linked either to the knapping processes or to other unspecific percussive tasks. The size and characteristics of the sample shows that lithic behaviors at this spot are mostly related to the casual exploitation of quartz tabular slabs to produce small usable flakes. The preferred selection of this blank type is confirmed by the fact that all identified cores and purported core fragments show a cubic morphology, in agreement with the shape of natural slabs as found at the quartz outcrops. Considering their contribution to the core sample (n=3), it seems that most of these blanks were exploited following the anvil method. Although clear bipolar products are significantly less numerous than freehand flakes, it is important to note that a significant fraction of waste (some debris, shatter and fragments) is compatible with the bipolar reduction of quartz slabs. A single exhausted multifacial-multipolar core is indicative of the presence of handheld percussion. As might be the case of bipolar knapping, usable freehand flakes outnumber negative scar count in this exhausted core. However, considering that this specimen could have gone through an intense process of blank reduction, the imbalance between on-site detached and retrieved flakes could be considered moderate. By means of bipolar and freehand knapping small-sized usable flakes with a mean cutting edge length of 32 mm were obtained. Considering that mean maximum length of these small flakes is 33 mm, an efficient balance between detached flakes and usable natural cutting edges is observed. None of these natural edges shows signs of transformation or resharpening via retouch or trimming. Among the flake sample, a small-sized phonolite specimen has been retrieved. Considering both the lack of volcanic cores in the sample and the particularly efficient ratio between size and cutting-edge length in this specimen, this might be a good example of inward fluxes of high-quality flakes produced elsewhere and consumed on-site. This movement of products along different fractions of the ancient landscape might have affected some quartz cores and/or flakes, although it seems more plausible that quartz exploitation and consumption took place preferentially on-site. In sum, at EAK hominins were undertaking casual quartz knapping activities and probably transporting some products obtained with the main goal of obtaining small usable natural cutting edges.

**References**

Diez-Martín, F.; Sánchez Yustos, P.; Mabulla, A.; Domínguez-Rodrigo, M.; Barba, R. 2009a. Were Olduvai hominins making butchering tools or battering tools? Analysis of a recently excavated lithic assemblage from BK (Bed II, Olduvai Gorge, Tanzania). Journal of Anthropological Archaeology 28: 274-289.

Diez-Martín, F.; Domínguez-Rodrigo, M., Barba, R., Tarriño, A., Sánchez, P., Luque, L., Mabulla, A., Prendergast, M. 2009b. The MSA/LSA technological transition in East Africa. New data from Mumba rockshelter Bed V and their implications in the origin of modern human behaviour. Journal of African Archaeology 7: 147-173.

Diez-Martín, F.,Sánchez, P., Domínguez-Rodrigo, M., Prendergast, M.E. 2011. An experimental study of bipolar and freehand knapping of Naibor Soit quartz from Olduvai Gorge (Tanzania). American Antiquity 112: 690-708.

Diez-Martin, F., Fraile, C., Uribelarrea, D., Sanchez-Yustos, P., Dominguez-Rodrigo, M., Duque, J., Diaz, I., de Francisco, S., Yravedra, J., Mabulla, A. & Baquedano, E. 2017. SHK Extension: A new archaeological window in the SHK fluvial landscape of Middle Bed II (Olduvai Gorge, Tanzania). Boreas 46, 4: 831-859.

Diez-Martín F., Duque Martínez, J., Fraile Márquez, C., Cobo Sánchez, L., Baquedano, E., Mabulla, A., Domínguez Rodrigo, M. 2024. Reconstructing early human behavior through the technological analysis of DS. In Domínguez-Rodrigo, Cobo-Sánchez, L., M., Mabulla, A., Baquedano, E., Gidna, A., Diez-Martín, F., eds: Reconstructing Olduvai. The behavior of early humans at David's site. Academic Press, London, pp. 241-282.

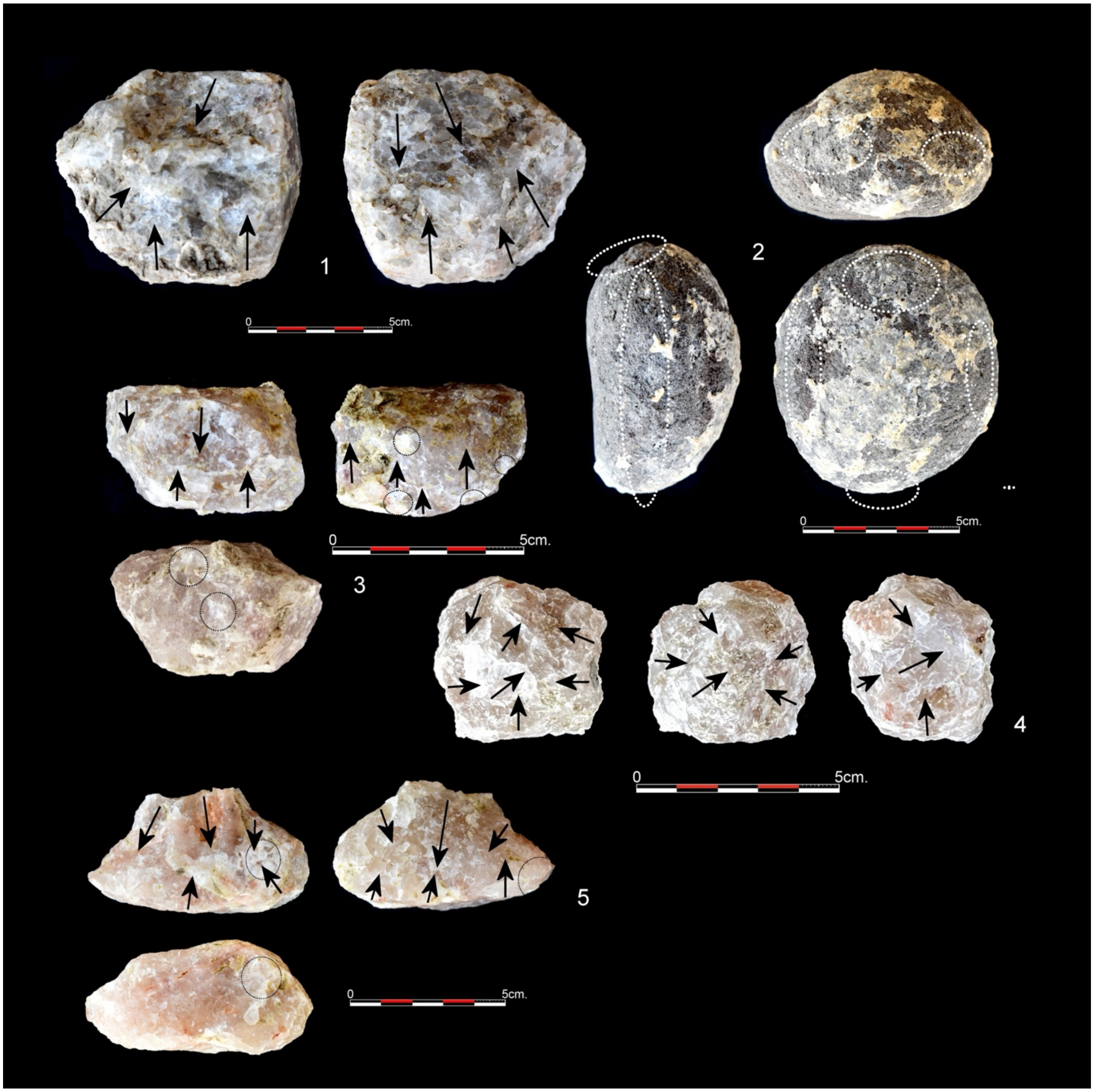

*Figure S26. Lithics from EAK. Bipolar cores: 1 Cubic specimen (77x70x58 mm, 455 g) from which a medium-sized flake (47x63 mm) has been detached; 3. A cubic blank has been exploited in the thickness plane around the perimeter of the piece with a circular tendency (51x47x32 mm, 194 g). Frosting is observed on the basal area and on some ridges; 5. Triangular morphology and opposed detachments, showing clear frosting on the base (71x40x35 mm, 118g). Hammerstone: 2. Oval basalt cobble (78x70x48 mm, 323 g) showing battering around the perimeter of the piece. Handheld cores: 4. Exhausted multifacial-multipolar core with a cubic shape (41x40x30 mm, 75 g), showing a minimum number of 10 negative scars crossing in multiple planes.*

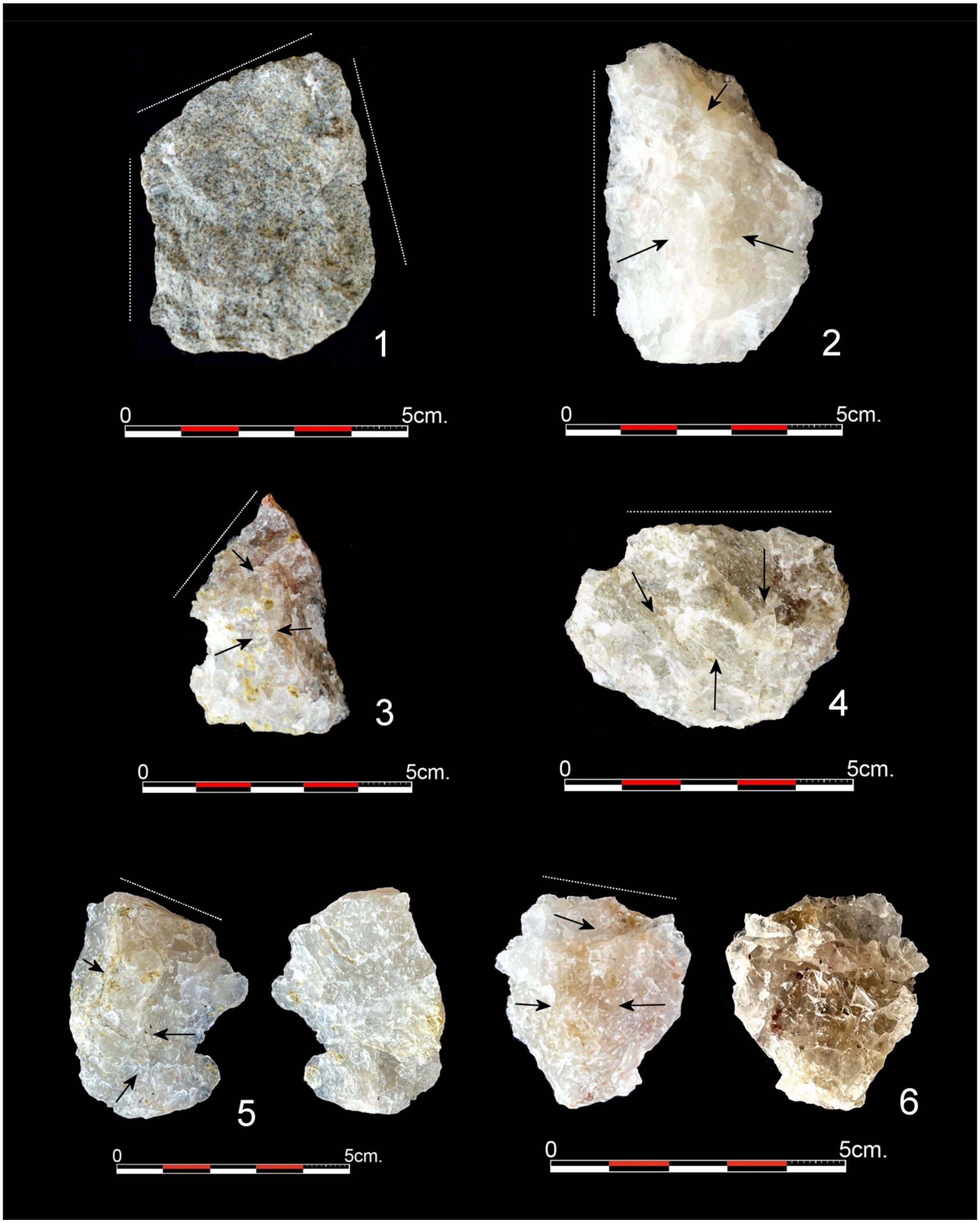

*Figure S27. Flakes from EAK. Phonolite: 1. Rectangular, Type 6 specimen, unifaceted butt, one dorsal negative scar, with 92 mm of bilateral and distal cutting edge (48x38x13 mm, 29g). Quartz: 2. Rectangular, Type 6, unifaceted platform, simple orthogonal dorsal pattern and 50 mm lateral cutting*

*edge (58x39x19 mm, 42 g); 3. Triangular, Type 6, unifaceted platform, simple orthogonal dorsal pattern and 30 mm of latero-distal edge (recent notch on the left side) (43x28x9 mm, 13 g); 4. Oval, Type 6, lineal platform, opposed dorsal pattern and 40 mm of transversal cutting edge (35x50x12 mm, 19g); 5. Quadrangular, Type 6, unifaceted butt, orthogonal pattern, diffuse bulb on ventral face and 28 mm of distal cutting edge (48x37x14 mm, 24 g); 6. Oval, Type 6, unifaceted butt, lateral opposed dorsal pattern, and 20 mm of distal cutting edge (36x32x13 mm, 13 g).*

**Tables**

| Lithic category | Raw material | | | | Total | |
|---|---|---|---|---|---|---|
| | V | B | P | Q | n | % |
| Unmodified | | | | | 1 | 1.25 |
| Cobbles | 1 | | | | | |
| Percussion | | | | | 1 | 1.25 |
| Hemmerstones | | 1 | | | | |
| Cores | | | | | 4 | 5 |
| Handheld | | | | 1 | | |
| Bipolar | | | | 3 | | |
| Detached | | | | | 23 | 28.75 |
| Flakes | | | 1 | 14 | | |
| Broken flakes | | | | 4 | | |
| Bipolar flakes | | | | 4 | | |
| Waste | | | | | 51 | 63.75 |
| Flake fragments | | | | 14 | | |
| Debris | | | | 24 | | |
| Shatter | | | | 10 | | |
| Fragments | | | | 3 | | |
| Total n | 1 | 1 | 1 | 77 | | |
| Total % | 1.25 | 1.25 | 1.25 | 96.25 | | |

*Table S2. Distribution of the lithic specimens retrieved from EAK sorted by lithic category and raw material. V, vesicular; B, basalt; P, phonolite; Q, quartz.*

| | Min | Max | Mean | SD |
|---|---|---|---|---|
| L | 20 | 58 | 32.76 | 10.66 |
| B | 19 | 50 | 29.41 | 7.97 |
| T | 6 | 19 | 11.17 | 3.66 |
| M | 4 | 42 | 13.35 | 10.56 |

*Table S3. Size (L= length, B= breadth; T= thickness) and mass (M) of complete flakes retrieved from EAK.*

### Orientation of materials at EAK

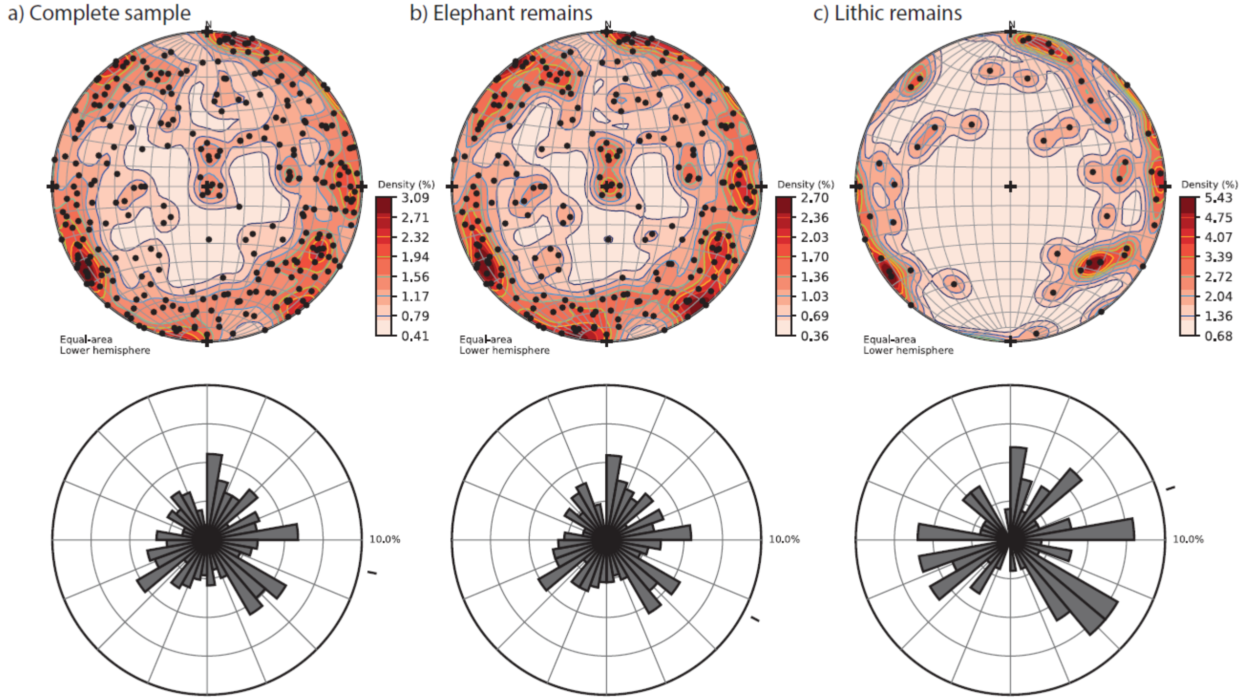

| | Statistic | p-value |
|---|---|---|
| Rayleigh test of uniformity | 0.0557 | 0.3868 |
| Kuiper's one-sample test | 1.3794 | >1.15 |
| Watson's godness of fit test | 0.0764 | >0.10 |

| | Statistic | p-value |
|---|---|---|
| Rayleigh test of uniformity | 0.0397 | 0.6671 |
| Kuiper's one-sample test | 1.0078 | >0.15 |
| Watson's godness of fit test | 0.0396 | >0.10 |

| | Statistic | p-value |
|---|---|---|
| Rayleigh test of uniformity | 0.1564 | 0.2944 |
| Kuiper's one-sample test | 1.5327 | >0.15 |
| Watson's godness of fit test | 0.0966 | >0.10 |

*Figure S28. Stereograms, rose diagrams, and results obtained from the application of the Rayleigh, Kuiper, and Watson tests.*

**Carnivore damage on an adult giraffe (Chobe National Park, Botswana)**

In late May 2024, two of us (MDR & EB) carried out a taphonomic study of proboscidean carcasses in a National Park and a Reserve in Botswana (analysis in progress). Here we will advance some green-broken specimens (most likely caused by hyena ravaging) found on a cluster of bones belonging to an adult giraffe in Chone National Park. This is relevant to see the extend of damage that carnivores can do on megafaunal remains.

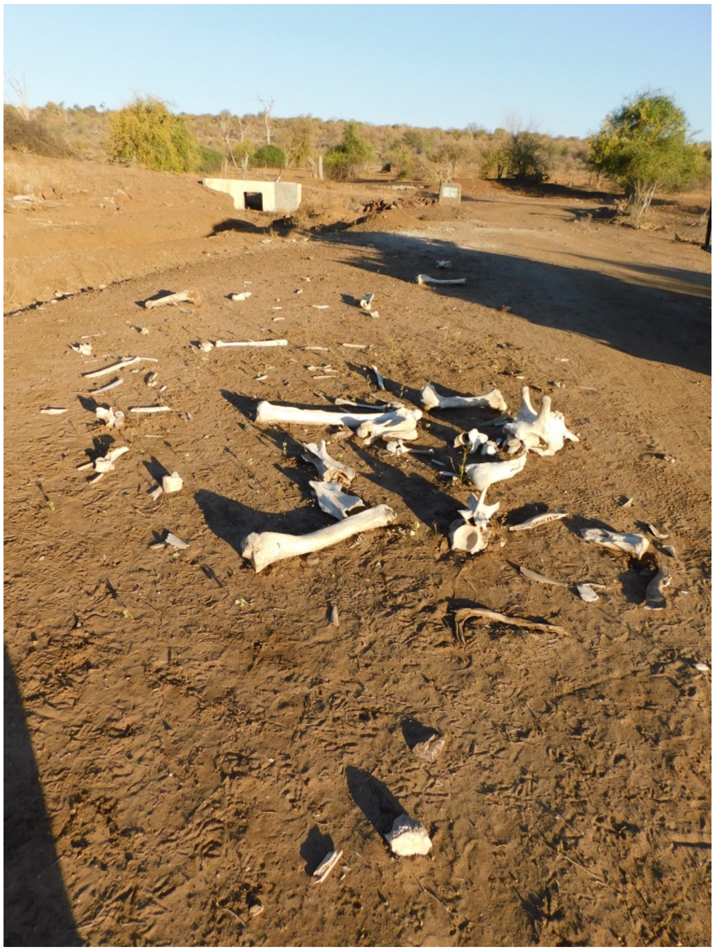

Figure S29. Cluster of hyena-ravaged bones from a giraffe at Chobe (Botswana).

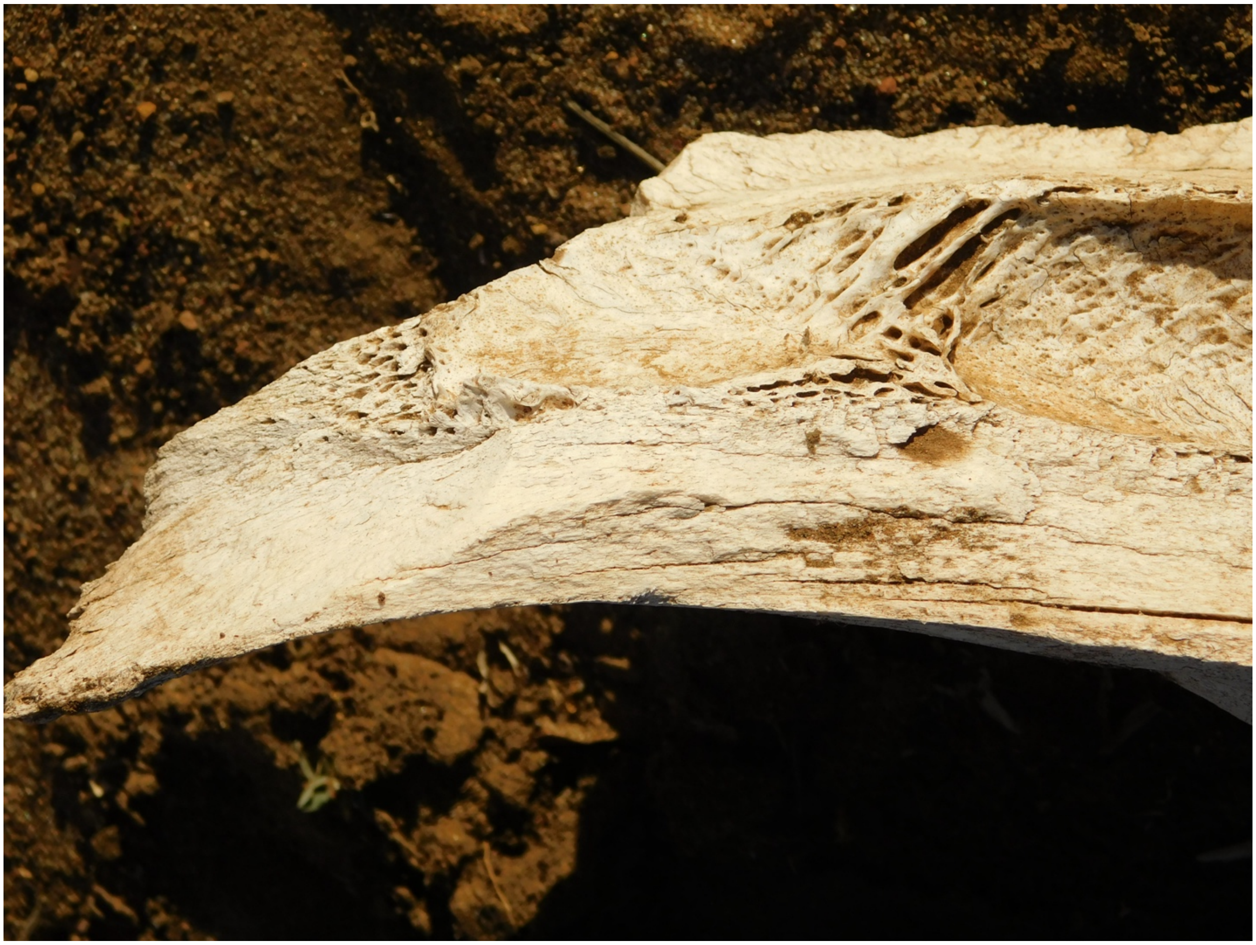

Figure S30. Spiral and green-broken planes on both sides of a giraffe humerus (broken by hyenas) from the cluster displayed in Fig. S22.

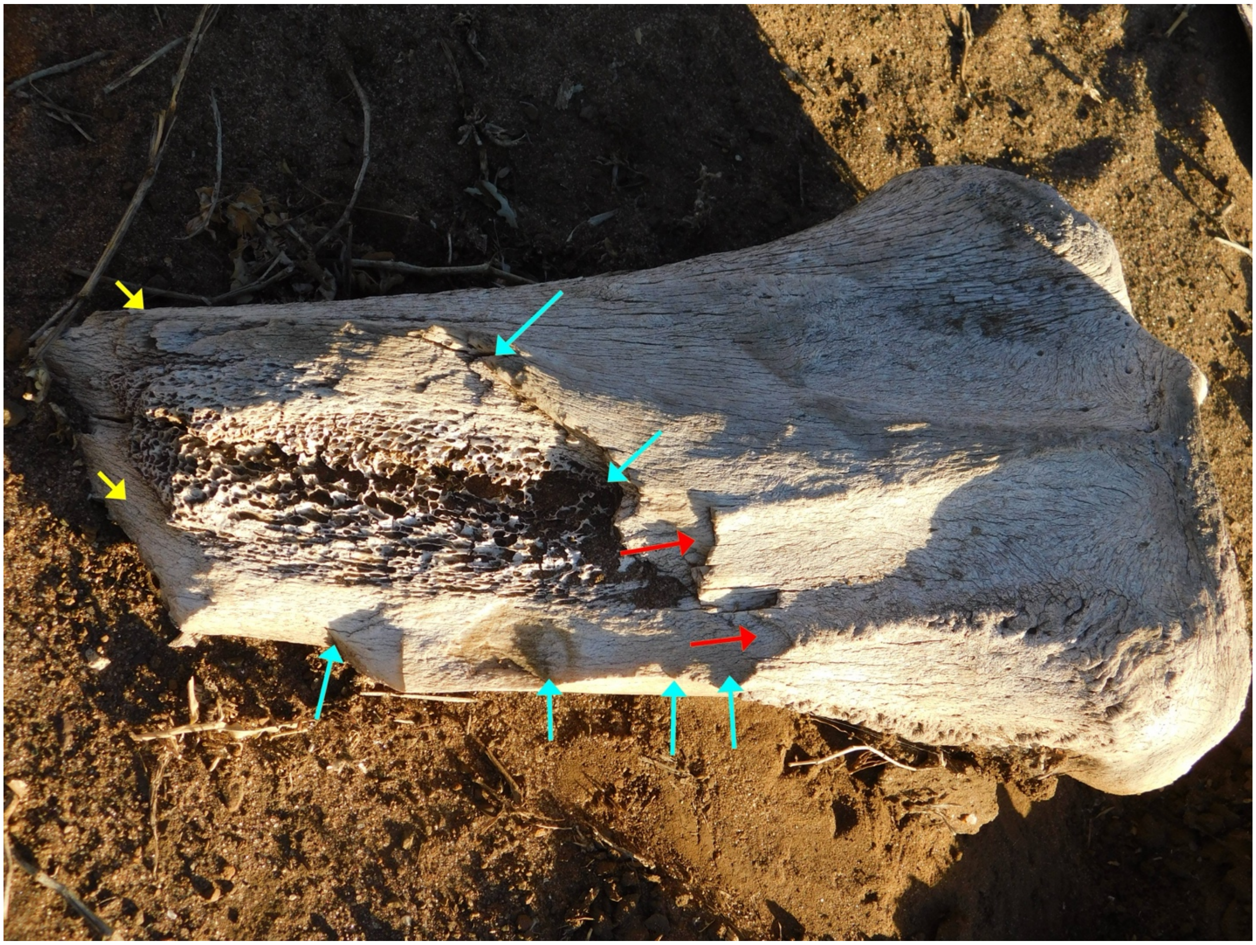

*Figure S31. Giraffe scapula ravaged by hyenas displaying overlapping non-invasive notches (blue arrows), continuous green breaks (yellow arrows), and axial reflected scars parallel to the axis of the bone (also documented in hammerstone broken megafaunal limb bones; see Fig S4).*

**THE ISSUE OF BONE TOOLS**

Here, we show some examples of additional carnivore damage on large-sized animals, displaying features that could be mistaken with anthropogenic agency.

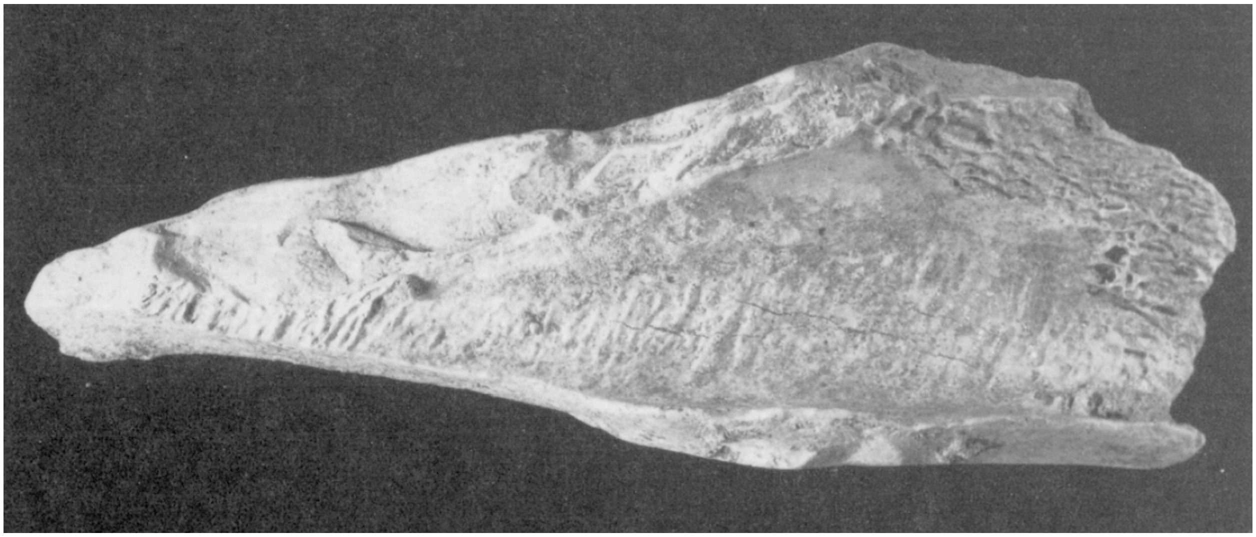

*Figure S32. Example of Bos/Bison femoral shaft from the hyena den of Bois Roche (from Villa & Bartram, 1996, Flaked Bone from a Hyena Den, Paleo 8: 143-159). Notice the amount of scars, their overlapping and incomplete nature, and their continuous trajectory on what seems a pointed bone fragment.*

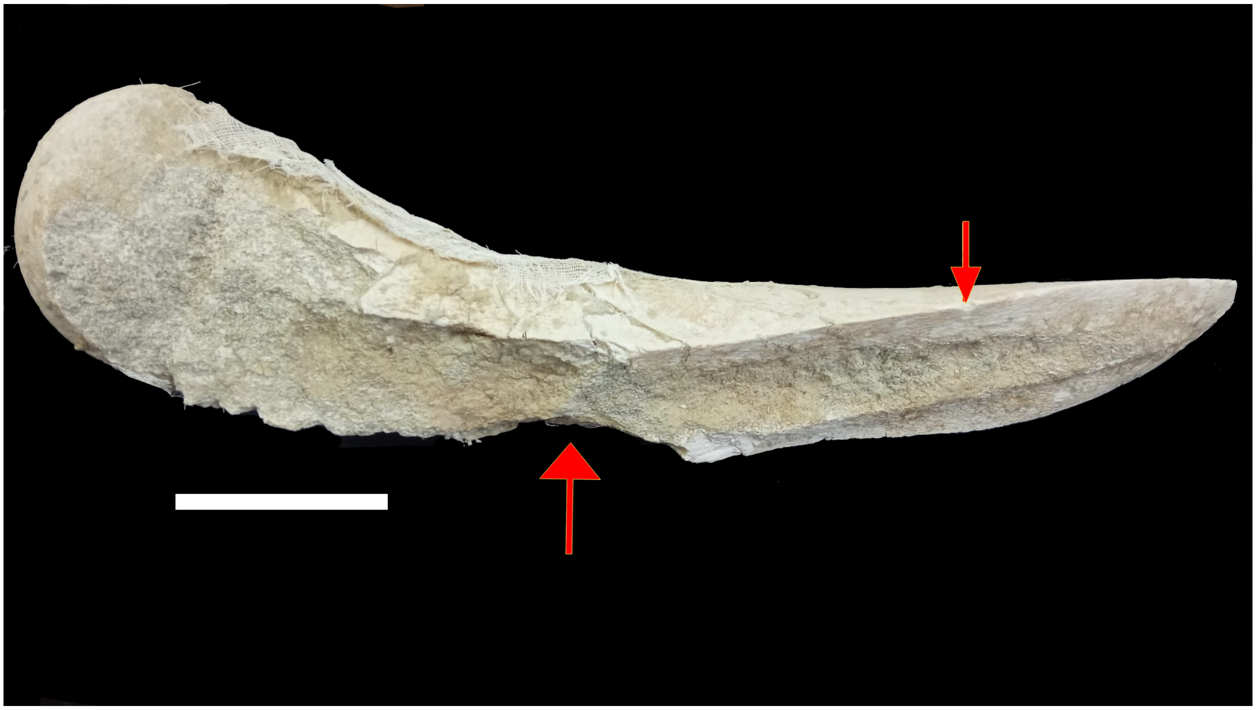

*Figure S33. Example of green-breakage on a proboscidean femur (Gomphoterium) from the Middle Miocene site of Virgen del Puerto (Madrid, Spain).*

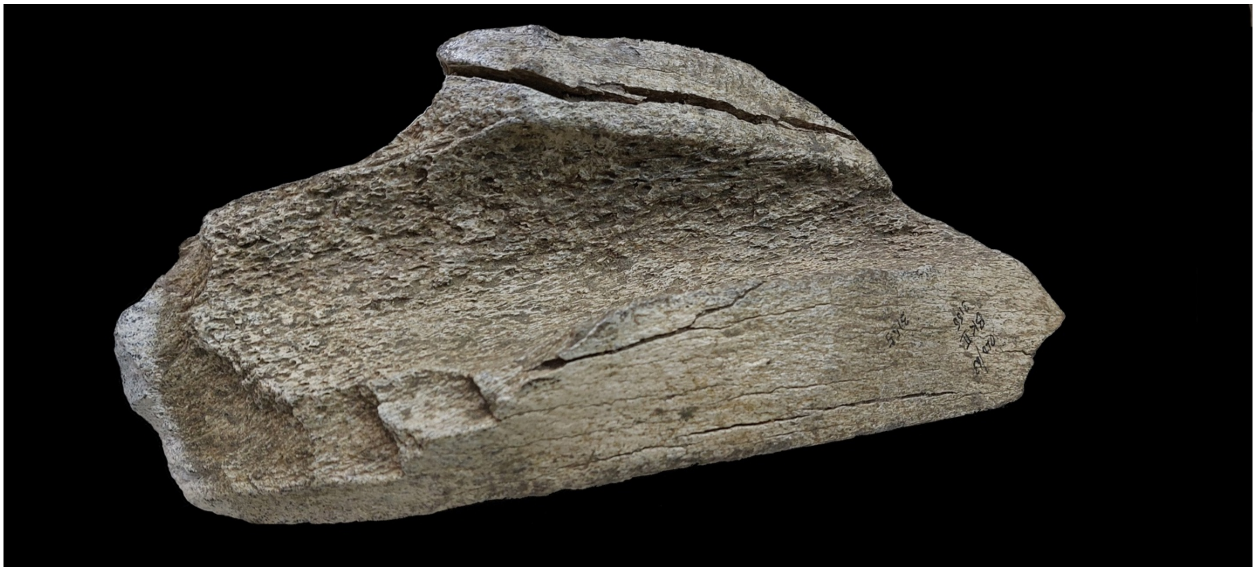

*Figure S34. Example of elephant green-broken femoral shaft fragment from BK (Olduvai, Upper Bed II) presented by Leakey as a case of bone tool. Notice the similar scar overlap and smooth green fracture surfaces resulting from bone breaking. The specimen´s length is 46 cm. Currently displayed at the Olduvai Museum.*

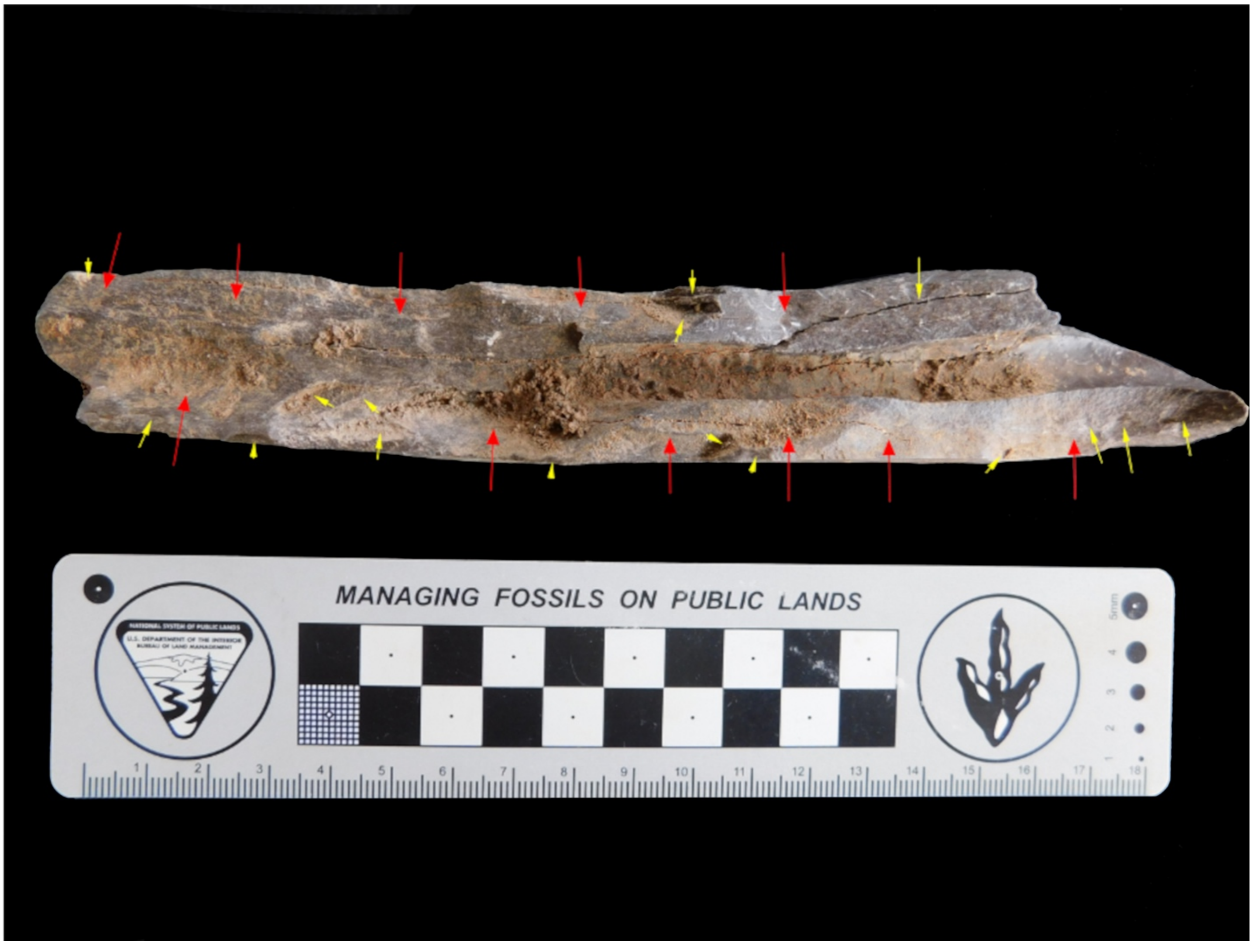

*Figure S35. Tibia fragment from a large medium-sized bovid displaying multiple overlapping scars on both breakage planes inflicted by carnivore damage (red arrows). Within those scars, there are abundant reflected/incomplete smaller scars (yellow arrows). The specimen was found within the base of the LAS stratigraphic unit, where the most abundant number of megafaunal green-broken bones have been found.*

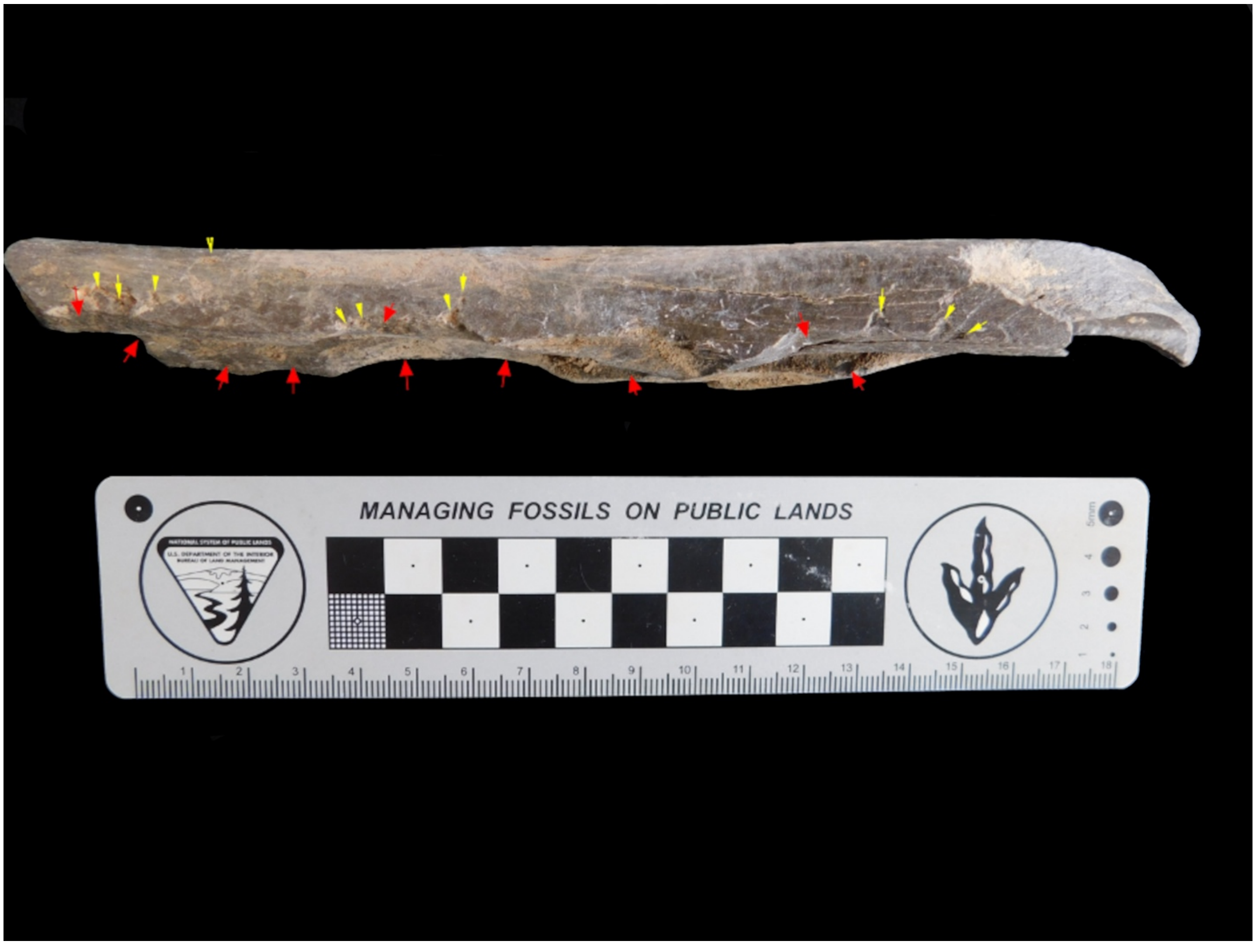

*Figure S36. Same specimen as shown in Figure S30, viewed on its medial side. The profile of the overlapping scars and notches can be clearly seen (red arrows), together with traces of tooth marks (yellow arrows).*
